# The Human Dendritic Cell Atlas: An integrated transcriptional tool to study human dendritic cell biology

**DOI:** 10.1101/2022.05.12.491745

**Authors:** Zahra Elahi, Paul W. Angel, Suzanne Butcher, Nadia Rajab, Jarny Choi, Justine D. Mintern, Kristen J. Radford, Christine A. Wells

## Abstract

Dendritic cells (DCs) are functionally diverse and are present in most adult tissues, however progress in understanding human DC biology is hampered by a relatively small number of these in circulation and by limited access to human tissues. We built a transcriptional atlas of human DCs by combining samples from 14 expression profiling studies derived from 10 laboratories. We identified significant gene expression variation of DC subset-defining markers across tissue-type and upon viral or bacterial stimulation. We further highlight critical gaps between in vitro-derived DC subsets and their in vivo counterparts and provide evidence that monocytes or cord blood progenitor in vitro-differentiated DCs fail to capture the repertoire of primary DC subsets or behaviours. In constructing a reference DC atlas, we provide an important resource for the community wishing to identify and annotate tissue-specific DC subsets from single-cell datasets, or benchmark new in vitro models of DC biology.

**Key Points:**

- A reference atlas of human DC that allows benchmarking of in vitro DC models
- Meta-analysis of 14 integrated studies demonstrate that human conventional dendritic cells have distinct tissue-of-origin phenotypes
- User uploads allow tissue-relevant annotation of human DC subsets from single cell datasets
- Key subset markers are altered by tissue or activation status
- Gaps between in vitro-differentiated DC and in vivo counterparts are partially rescued by humanized mouse models, or coculture with NOTCH-ligands.

## Introduction

Resident in most tissues, dendritic cells (DCs) are specialized antigen-presenting components of the immune system that play a key role in the mounting and regulating of antigen-specific responses by T and B lymphocytes. Their role in bridging innate and adaptive immunity has made them a popular target in vaccine development, immunotherapies, and autoimmune disease treatments (reviewed by Calmeiro et al., 2020; Wylie et al., 2019). There are at least three main classes of DCs that are distinct in morphology, phenotype, and function. These classes include plasmacytoid DCs (pDC), two subsets of conventional DCs (cDC1 and cDC2) and monocyte-derived DCs (MoDC). The latter are not present at steady state but can be differentiated from monocytes during inflammatory conditions. Individual transcriptome and functional studies have provided a detailed insight into DC biology, but these show little agreement with respect to the subset-specific transcriptional markers. For example, others have shown that only 4.2% of the signature genes associated to cDC1 were common to 3 recently published datasets (reviewed by Balan et al., 2020 for Heidkamp et al., 2016; See et al., 2017; Villani et al., 2017).

As DCs are a rare cell population in human tissues, our understanding of their biology has largely arisen from model organisms, but cross-species comparisons have revealed the distinctiveness of mouse DC subsets from human equivalents. These include the absence of shared expression of subset-defining markers (reviewed by Macri et al., 2018), toll-like receptor 8 (TLR8) mediated responses to single-strand RNA (Heil et al., 2004) and expression of co-stimulatory molecules after stimulator of interferon genes (STING)-dependent activation (Pang et al., 2022). To study DC heterogeneity and complexity in humans, many studies have relied on putative DC isolation from blood. For example, one of the first single cell transcriptome experiments of blood DC (Villani et al., 2017) described a new DC subset termed AXL^+^ SIGLEC6^+^ (AS) DCs. Although some individual studies have described DC subsets from other tissues including spleen (Brown et al., 2019; Heidkamp et al., 2016; McGovern et al., 2017), intestine (Watchmaker et al., 2014), skin (Haniffa et al., 2012), or bone marrow (van Leeuwen-Kerkhoff et al., 2018), comparisons between these individual datasets are *ad hoc* in the absence of a reference that can integrate information from diverse sources.

Given the importance of DCs in immunity, and their distinctive phenotype and functions compared to other myeloid components including macrophages and monocytes, the lack of laboratory models of these cells is becoming a critical bottleneck for the field. The current *in vitro* models of human DC biology rely on the differentiation of CD34+ hematopoietic stem cells (HSC) from blood or bone marrow (Balan et al., 2014), are differentiated *in vitro* from peripheral blood monocytes (Pacis et al., 2015), or use induced pluripotent stem cells (iPSC)(Monkley et al., 2020) or directed differentiation from fibroblasts (Rosa et al., 2018, 2022). Concordance between cord-blood derived cDC1 with their *in vivo* blood counterparts was previously reported (Balan et al., 2018). However, the experimental setup did not assess both sources simultaneously, and similarity was evaluated with a limited set of markers for each subset. Therefore, introducing a platform that incorporates different models of DCs and includes freshly isolated tissue-resident cells (*in vivo*), primary DC cultured after isolation (*ex vivo*), DC models generated *in vitro*, or isolated from humanized mouse models (*in vivo* HuMouse) (Minoda et al., 2017) benefits the community by providing the opportunity to compare and evaluate subsets of DCs across multiple sources.

Here we undertake the first systematic evaluation of tissue-resident, *ex vivo* and *in vitro* models of human DC biology by constructing a reference human DC atlas that integrates data from multiple laboratories, derivation methods, and measurement platforms. We observe differential expression of subset-specific genes between tissues of origin and status of activation which suggests being cautious with over-reliance on a small set of markers to identify DC subsets. The DC atlas reveals a transcriptional gap between *in vitro*-derived DCs and their primary counterparts, highlighting the need for improvement of *in vitro* differentiation models.

## Results

### Assessing the reproducibility of DC expression phenotypes in the Stemformatics DC atlas

As DCs are relatively rare cell types, it has been difficult to directly compare subsets derived from different tissues and evaluate how disease or antigen activation alter their molecular phenotypes. Most published studies on human DC biology are described for a specific experimental context, such as profiling blood DCs (See et al., 2017) or comparing the transcriptional phenotype of *in vitro*-derived DCs with those from blood (Balan et al., 2014). Comparing DC phenotypes derived from different studies remains a challenge for the field. We set out to develop a unified description of DC biology by assessing which transcriptional phenotypes were generalisable across multiple studies. The resulting DC atlas has been constructed using 342 human samples from 14 studies including tissue-resident, *ex vivo* and *in-vitro* generated conventional and non-conventional DCs (Figure 1, Supplementary Figure 1, Supplementary Table 1). Samples were annotated using evidence from the original study consisting of fluorescence-activated cell sorting (FACS) accompanying each DC subset, tissue of origin, and where relevant, culture conditions, disease and activation status.

**Figure 1.**
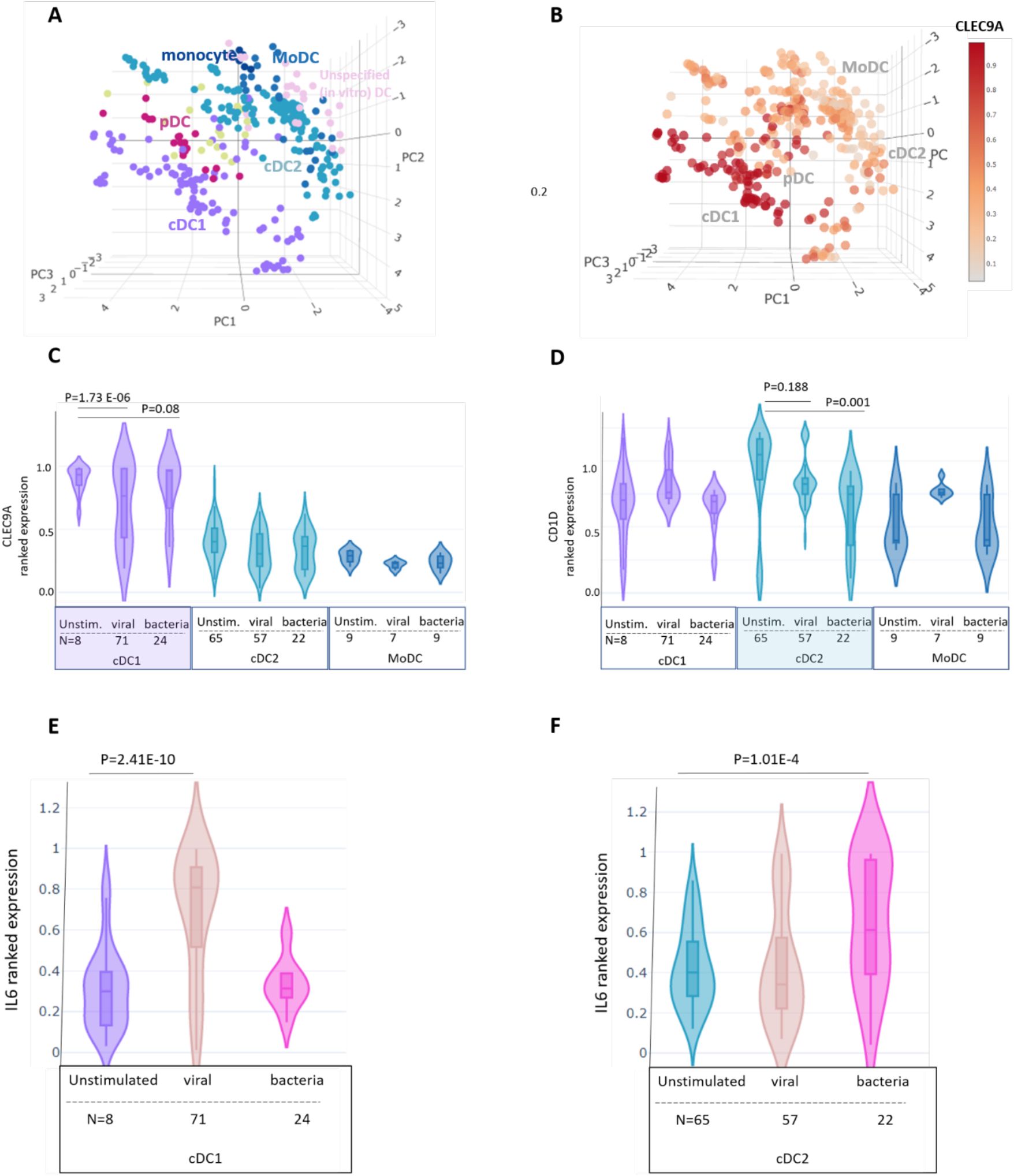
The Dendritic Cell Atlas in Stemformatics.org consists of 342 samples derived from 14 transcriptome studies and 10 laboratories. (A) Annotation of samples by DC subtype - purple cDC1, curious blue cDC2, red-violet pDC, denim MoDC, congress blue monocyte, yellow-green DC precursor, pink unspecified dendritic cell. (B) Overlay of *CLEC9A* expression highlights behaviour as a marker of cDC1. Colour intensity ranges from high expression (dark red) to low/no expression (pale red). (C) Ranked gene expression (median, interquartile range) of (C) *CLEC9A* and (D) *CD1D* comparing activation status between bacterial, unstimulated or viral agonists. Sample sizes (N) are given under each combined category. p value: independent t-test. (E) Ranked gene expression (median, interquartile range) of *IL6* for the subsets of (E) cDC1 and (F) cDC2 comparing activation status between bacterial, unstimulated or viral agonists. Sample sizes (N) are given under each combined category. p value: student t-test. See also Figure S1.

The dominant pattern shared by all studies was the clustering of known DC subtypes, which allowed us to assess the expression of genes commonly used to discriminate between these groups (Figure 1A). Monocyte-derived dendritic cells (MoDC), for example, clustered with monocytes and were closely associated with the conventional dendritic cell 2 (cDC2) cluster. A generic “dendritic cell” label was assigned to the samples that lacked sufficient subtype information, and as these were predominantly derived *in vitro* from CD34+ cells, they were closely associated with cDC2 and MoDC subsets. Plasmacytoid dendritic cells (pDC) grouped closely with progenitors as well as conventional dendritic cell 1 (cDC1) subsets. Examination of technical variables such as study ID or platform (Supplementary Figure 1A-B) showed that these were not contributing to clustering of individual samples, or the DC subsets.

cDC1 cells have been described in both blood and primary tissues and are important for immune responses to pathogen and tumour (reviewed by Merad et al., 2013). Samples were annotated as cDC1 if the original study had provided FACS data using current benchmarks of cDC1, including *CD141, CLEC9A* and *CADM1* (Figure 1B, Supplementary Figure 1C). cDC2 are more abundant and are specialized in major histocompatibility complex (MHC) class II antigen presentation (same citation, Merad). *CD1C* is a common marker of cDC2 population and is widely used to isolate this cell type from blood and other tissues. In evaluating marker distribution across the cDC subsets, we noted high expression of *CD1C* mRNA by *in vitro*-differentiated cDC1 and pDC, but not tissue-resident equivalents (Supplementary Figure 1D-E), which is in agreement with previous research showing FLT3L-driven differentiation of progenitors induces expression of *CD1C* marker in cDC1s (Kirkling et al., 2018; Poulin et al., 2010).

Many of the classical markers of cDC1 and cDC2 cells displayed varying levels of expression on activation with bacterial or viral ligands. For example, cDC1 significant markers *CLEC9A* and *TLR3* and cDC2 markers *CLEC10A* and *CD1D* were downregulated by bacterial or viral activation (Figure 1C-D, Supplementary Figure 2A-B). Activation markers such as *IL6* and *TNFAIP6* demonstrated subset-specific activation profiles: bacterial ligands induced *IL6* and *TNFAIP6* expression in cDC2, whereas these factors were induced by viral ligands in cDC1 (Fig 1E-F, Supplementary Figure 2C-F). This is consistent with the reciprocal expression pattern of the bacterial and viral adjuvants’ receptors in cDC1 and cDC2; viral adjuvant receptor, TLR3, is highly expressed by cDC1 while bacterial adjuvant receptor, TLR4, is highly expressed by cDC2 (Leal Rojas et al., 2017). Thus, the ability to look across multiple experimental attributes allows users of the DC atlas to assess the conditions likely to reproducibly alter cDC subset-specific behaviours.

**Figure 2.**
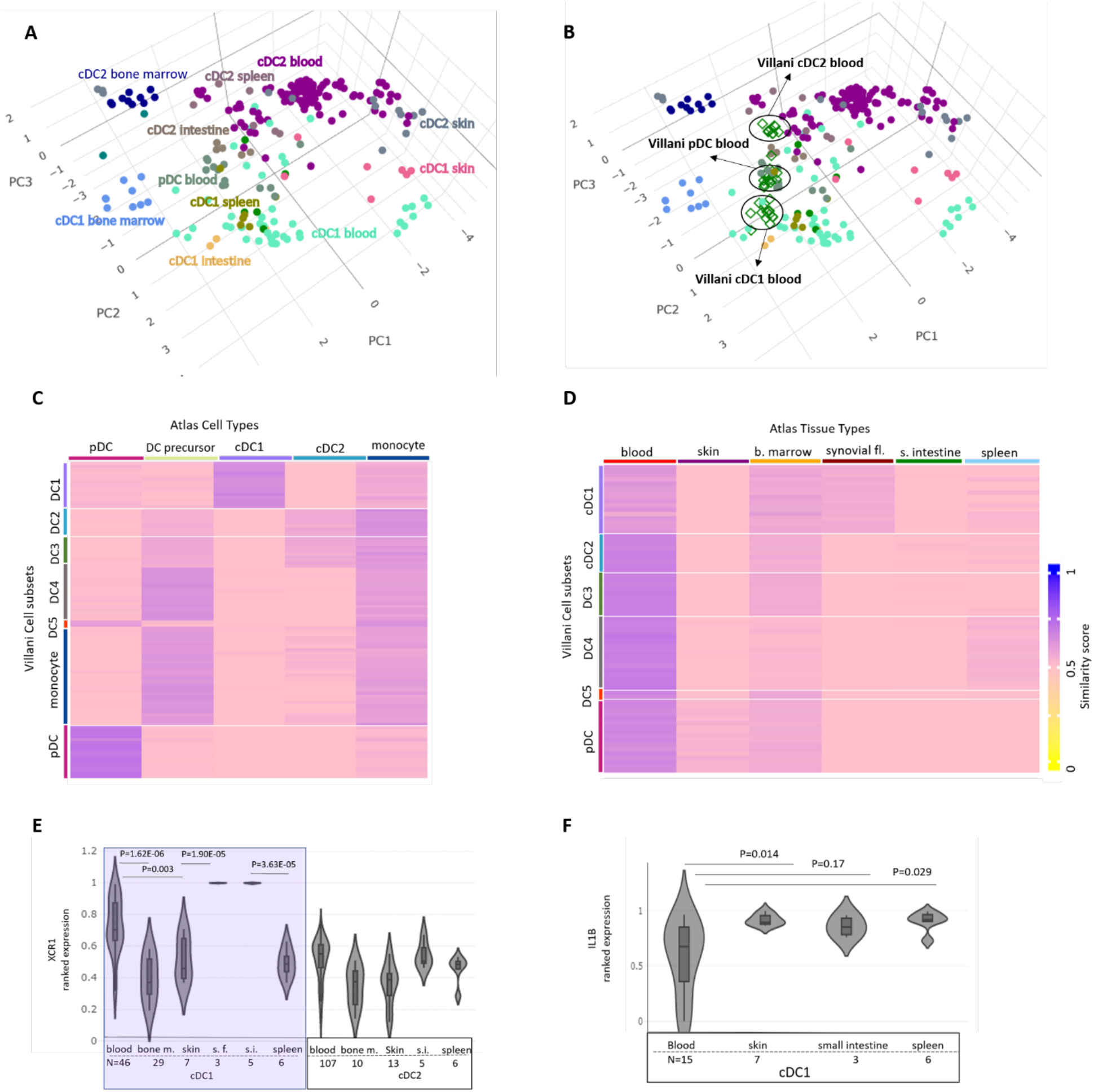
Tissue-specific behaviour of DC. (A) Stemformatics DC atlas cluster by subset and tissue of origin. (B) Projection of pseudo-bulk samples from single-cell data describing blood-derived DC from (Villani et al., 2017) demonstrates reproducible grouping of new samples on the reference atlas. (C) Annotation of cell type from Villani single-cell libraries using the Capybara similarity score against the reference Stemformatics DC atlas predicting DC subsets and (D) tissue of origin. Similarity scores high(blue) to low (yellow). (E)Ranked gene expression of *XCR1* (median, interquartile range) in samples selected for DC subtype cDC1 and cDC2 from blood, bone marrow (bone m.), skin, small intestine(s.i.), spleen and synovial fluid(s.f.). Sample size (N) for each combined category listed under x-axis. (F) Ranked expression of *IL1B* gene (median, interquartile range) in normal (disease status) *in vivo* cDC1 samples isolated from blood, skin, small intestine (s.i.) and spleen. Sample size (N) for each combined category listed under x-axis. p-value: student t-test.

### Tissue of origin alters DC behaviour

The tissue environment shapes human DC phenotypes (reviewed by Roquilly et al., 2022), and indeed, the DC atlas demonstrated reproducible patterns of DC subsets grouping by related tissue types. For example (Figure 2A), spleen and small intestine groups overlapped substantially with blood clusters, but DC isolated from primary human skin biopsies grouped together and were distinct from DC isolated from humanised mouse bone marrow. In order to assess the reproducibility of tissue clusters of DC subsets, we projected pseudo-bulk samples of an external single-cell dataset of blood-derived DC subsets (Villani et al., 2017) on the reference atlas. Projection of these samples showed the association of this data with the reference DC type, and tissue (Figure 2B-D). The Capybara similarity analysis predicts the specific cell types of the Villani dataset and the tissue of isolation, blood, by showing a high similarity score with DC atlas samples from blood. This is further evidenced by lower Capybara similarity scores for bone marrow, spleen, skin and intestinal DCs. Projection of high-resolution single-cell data like the Villani dataset on the DC atlas can be carried out on the Stemformatics platform and provides users with the opportunity to compare and annotate their own samples against the reference atlas.

Most isolation methods rely on specific markers to isolate DC subsets from varied tissue types. For example, CLEC9A, XCR1 and CADM1 are the most common markers for isolating blood and tissue-resident cDC1. However, a highly variable gene expression pattern of *XCR1* was identified across multiple tissue types (Figure 2E). This has implications when choosing a reference to identify and annotate innate immune cells isolated from different tissues. cDC1 isolated from peripheral blood have differentially low expression of pro-inflammatory factors interleukin 1-beta (*IL1B*), suggesting a less mature profile of blood cDC1 in comparison with cDC1 isolated from other tissues (Figure 2F). Therefore, the DC atlas is a useful reference tool to study the tissue-specific or activation behaviour of DC subsets.

### *in vitro*-derived dendritic cells do not recapitulate the biology of their *in vivo* equivalents

Human DCs develop from the HSCs resident in the bone marrow that later differentiate to common myeloid progenitors (CMP) and common dendritic cell progenitors (CDP). Although the ontogeny of human DCs is still being elucidated, many *in vitro* DC differentiation protocols have been developed in line with what is currently understood with regards to the DC developmental trajectory. Most human models of DC biology rely on differentiation from peripheral blood monocytes(Pacis et al., 2015) or CD34+ cord blood progenitors(Balan et al., 2014). It is critical to understand how faithfully these capture the repertoire of DC subsets or behaviours.

The transcriptional phenotypes of *in vitro*-differentiated cDCs from CD34+ cord blood progenitors (n=60) or monocyte-derived DCs (n=25) were distinct, shown by clustering away from tissue-isolated DCs (Figure 3A). We wondered if this lack of similarity is due to the presence of MoDC models *in vitro* but not *in vivo* conditions but looking at the smaller subsets of pDCs and cDC1s derived from FLT3L-dependant differentiation methods showed the same pattern (Supplementary Figure 3A-B). For example, B- and T-lymphocyte attenuator (*BTLA*), a significant marker of human cDC1 mediating immune regulation (Jones et al., 2016), is missing from the cDC1s differentiated from cord blood CD34+ progenitors, but its expression is rescued in cDC1s differentiated in the HuMouse models (Figure 3B). The improved methods of differentiation using a feeder layer of mouse OP9 stromal cells expressing Notch ligand Delta-like 1 (DLL1) (Balan et al., 2018) pushed the *in vitro*-derived DCs more toward the DCs derived *in vivo* in the HuMouse models, nevertheless these still sat in a different transcriptional cluster to their in vivo counterparts (Figure 2C, Supplementary Figure 3C).

**Figure 3:**
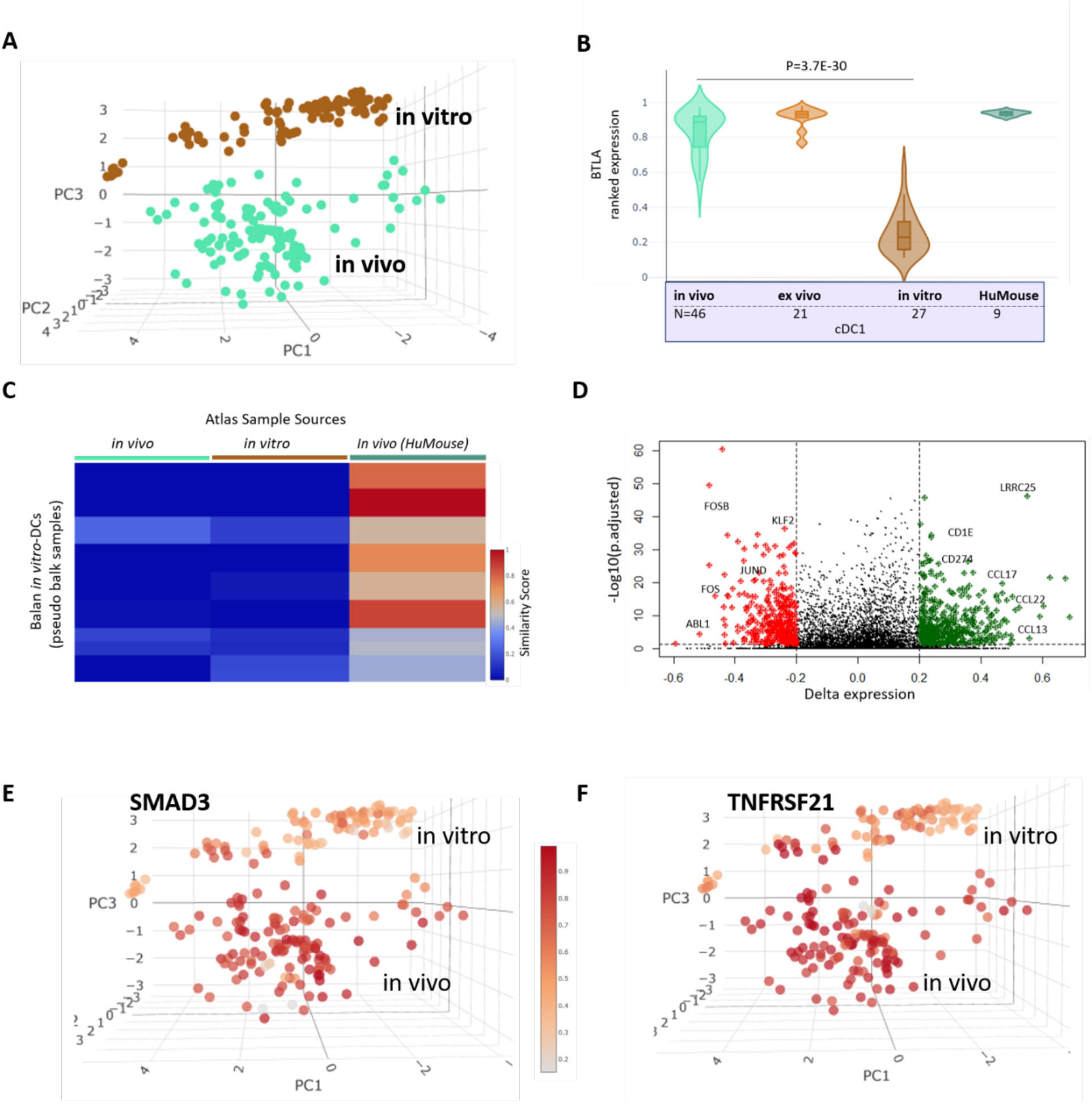
*In vitro* derived DC from CD34+ cord blood progenitors (n=60) and monocyte-derived DC (n=25) are transcriptionally distinct from primary cell types. (A) Stemformatics DC atlas coloured by *in vitro* (n=85) and *in vivo* (n=125) DC sources. (B) Ranked gene expression of cDC1 marker, *BTLA*, in *in vivo* cDC1, *ex vivo* cDC1, *in vitro-*derived from cord blood CD34+ progenitors and in HuMouse-differentiated cDC1. Sample size (n) for each combined category listed under x-axis. (C) Heatmap of similarity score of *in vitro*-derived DCs from single cell dataset by Balan et al. (2014) against the reference Stemformatics DC atlas using the Capybara analysis. Similarity scores high(red) to low (blue). (D) Volcano plot of differentially expressed genes lost (red) or gained (green) in *in vitro* DC compared to primary cells. (E) Overlay of *SMAD3* and (F) *TNFRSF21* gene expression between *in vivo* and *in vitro* samples. Colour intensity ranges from high expression (dark red) to low/no expression (pale red). p-value: student t-test.

We identified 3139 differentially expressed genes (DEG) between *in vitro*-generated DCs and *in vivo*, DCs (Supp. Table 2, listing top 1500 DEG), where key receptors such as *CSF3R, CCR9* and *TLR10* were amongst the downregulated DEG by *in vitro* samples (Supp. Figure 3D-E). The gene set enrichment analysis on top-ranked genes missing *in vitro* revealed that the most impacted biological processes are cell activation and communication, including interleukin 18 receptor 1 (*IL18R1*), tumour necrosis factor receptor, *TNFRSF21*, important for activating NF-KB pathway (Pan et al., 1998), signaling lymphocyte activation molecule (SLAM) family immunoregulatory receptor *CD244*, and *SMAD3* controlling responsiveness to transforming TGF-B1 cytokine(Peng et al., 2013) (Figure 3E_F, Supplementary Figure 3F-G). The missing factors from the transcriptional phenotype of the current *in vitro* DC models highlight the necessity to develop improved differentiation methods of *in vitro* DC derivation.

To better understand the phenotype captured *in vitro*, and to what extent this phenotype is different from *ex vivo* models, we compared the gene expression profile of stimulated and unstimulated cultured groups with *in vivo* samples. Four broad gene clusters were identified among the genes significantly captured by *in vitro* DCs. The genes that were upregulated by both *in vitro* and *ex vivo* models (clusters 3 and 4) were mostly enriched with metabolite and energy-related processes (Figure 4A), with a subset of these genes induced by viral activation specifically by the *in vitro* group (cluster 3).

**Figure 4.**
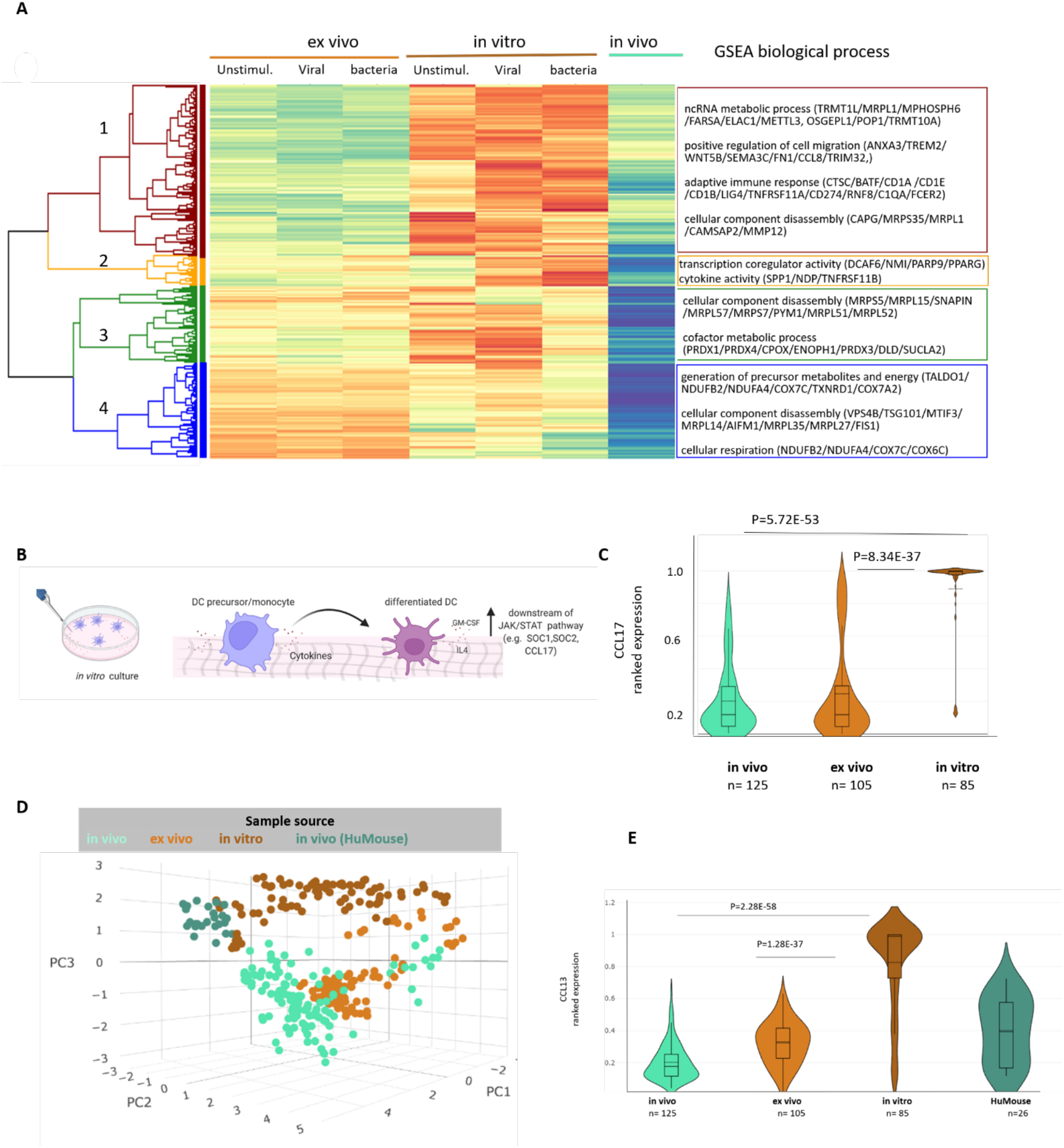
The phenotype captured by *in vitro* derived DC is distinct from *primary cells*. (A) Heatmap of differentially upregulated genes by *in vitro* DCs (p value <0.05) compared within varied activation statuses of cultured samples versus *in vivo* DCs, with gene set enrichment analysis (GSEA) of gene ontology terms (biological process) for each cluster including examples of some of the genes associated with each process (B) Schematic of the impact of differentiation cytokines on the transcriptional profile of *in vitro*-derived DCs. (C) Ranked gene expression of *CCL17* chemokine compared between *in vivo* (n=125), *ex vivo* (n=105) and *in vitro* (n=85) DC samples. (D) Stemformatics DC atlas coloured by sample source to highlight the resemblance of *ex vivo* (light brown) DCs to their *in vivo* (green) models but apart from *in vitro* (dark brown) differentiated DCs. (E) Ranked gene expression of *CCL13* chemokine in *in vivo* DCs (n=125), *ex vivo* DCs (n=105), *in vitro*-derived DCs (n=85) and humanised mouse (HuMouse)-differentiated DCs (n=26). p value: student t-test.

Genes that were upregulated *in vitro* but remained at a low expression level under *ex vivo* conditions and *in vivo* DC (cluster 1 and 2) were associated with the regulation of cell migration (e.g. *CCL8* and *WNT5B*), immunoregulatory functions (e.g. *CD274* and *PDCD1LG2*), communication with adaptive immune cells (e.g. *BATF* and *FCER2)* and the genes encoding CD1 family antigen-presenting molecules (*CD1A, CD1B, CD1E*) (Supplementary Figure 4A-B). As these proteins are associated with maturation phenotype of DCs, these data suggest that *in vitro*-generated DCs represent a partially activated profile, most likely explained by the presence of IL-4 and GM-CSF cytokines in most DC differentiation culture media (Fig 4B). For example, highly expressed genes by *in vitro* DCs such as *CCL17, CCL22* and *CCL1* chemokines are downstream of GM-CSF signaling (Globisch et al., 2014; Ushach and Zlotnik, 2016), and *SOCS1* and *SOCS2* genes are downstream of IL-4 signaling pathway (Jackson et al., 2004) (Figure 4C, Supplementary Figure 4C). This *in vitro* activated profile might have implications for their application in immunotherapy, as activated DCs have a limited lifespan (De Smedt et al., 1996). Certainly, we observed low expression of anti-apoptotic marker, *BCL-2*, by *in vitro* models (Supplementary Figure 4D).

### Cultured blood DCs remember their tissue environment

We noted above that cultured blood DC clustered closely with cell profiled directly from blood (Figure 4D). For example, the expression of chemokine *CCL13* is significantly high *in vitro* but *ex vivo* samples do not upregulate this gene after being in a culture environment (Figure 4E). This suggests that although *ex vivo* blood DCs have experienced a culture environment, they appear to maintain memory of their tissue environment. We don’t know how generalizable this is, but it suggests that the derivation strategy is a more important than the culture environment when considering DC behaviours such as their activation or maturation status.

## Discussion

DCs are specialized antigen-processing and antigen-presenters of the human immune system. However, our understanding of their biology is limited by their paucity in human tissues, and a lack of appropriate *in vitro* models. Our aim here was to develop a resource that allowed systematic benchmarking and assessment of diverse DC behaviours, by developing an integrated transcriptional atlas of human DCs that combined datasets from individual laboratories. We used a method of batch correction that does not require prior delegation of samples to biological categories, thereby avoiding normalization that predicates the analysis outcomes. In integrating many datasets together, emergent biological properties were reproducible across several independently derived studies, often from different laboratories. In doing so, we identified previously hidden variation of subset-defining markers across tissues or upon activation by viral or bacterial stimuli.

The DC atlas is implemented in Stemformatics.org, offering a readily accessible platform for benchmarking *in vitro*-generated models of *in vivo* biology. The human DC atlas is publicly available as an interactive 3D PCA graph that can be explored across a single or a combination of conditions such as cell type, tissue type, derivation source, disease or activation status. Users may project their own gene expression dataset, including single-cell RNA sequencing datasets, against the atlas for annotating subsets or benchmarking their models. The atlas is scalable and will grow as new datasets of DC biology become available.

Although individual studies emphasized the homology of their *in vitro*-generated models of DCs compared to putative DCs, here we identified a transcriptional gap between these two. The genes over-expressed by *in vitro* models can be explained by the presence of growth factors and cytokines in the dish environment and indicate an activated phenotype that may impact their longevity in clinical settings. The missing genes from the *in vitro* models were associated with their ability to develop, differentiate, and communicate with T cells. Nevertheless, cultured DC models did partially recapitulate *in vivo* DC biology. For example, cultured blood DCs clustered tightly with primary blood DCs, suggesting that they remember their tissue environment, or rather, have not yet been exposed to maturation factors in the tissue niche (reviewed by Merad et al., 2013; Roquilly et al., 2022). These observations suggest that DC require additional signals from the tissue stroma that are absent in a culture dish. While co-culture with stromal cells such as the OP1 line, or addition of NOTCH-signaling to the culture environment seek to address this gap, the DC atlas phenotypes show that these strategies only partially rescued the cord-blood derived DC phenotypes, as did reconstitution in HuMouse models. Altogether there are further opportunities to improve *in vitro*-DC derivation methods. The use of alternative progenitor sources, such as iPSCs, may assist in deconstructing these environmental gaps and provide an opportunity to understand the requirements, and impact, of various factors in shaping DC-development, subset specificity and function.

## Acknowledgements

This study was funded by NHMRC Synergy grant APP1186371 to C.A.W.

## Author contributions

Conceptualization C.A.W, Z.E; Data curation Z.E., P.W.A., S.B., J.C., N.R.; Formal analysis Z.E., P.A.W; Investigation Z.E., C.A.W; Methodology Z.E., P.W.A, J.C. ; Project administration C.A.W, J.C.; Resources C.A.W; Software P.A.W., J.C.; Supervision J.D.M., K.R., C.A.W; Validation Z.E., P.W.A., C.A.W; Visualization J.C; Writing – original draft Z.E., C.A.W; Writing – review and editing Z.E., C.A.W., S.B., N.R., J.D.M., K.R.

## Methods

### Data collection and processing

For atlas construction, publicly available datasets were collected from database repositories such as NCBI’s Gene Expression Omnibus (GEO) and EBI’s Array Express.

Mapping and analysis of microarray and RNA sequencing datasets were done using the standard Stemformatics processing pipeline described by (Choi et al., 2019). All datasets passed multiple stringent quality control (QC) steps required for hosting on the Stemformatics platform. Dataset processing scripts are available at Stemformatics GitHub. Datasets that have failed the QC steps were removed from the integration step. Finally, 14 datasets consisting of 342 samples were selected for inclusion in the atlas (Supplementary Table 1).

### Dataset integration and gene selection for PCA

For the integration step, the common genes between all datasets were selected and assessed for their platform dependency as described in detail by (Angel et al., 2020) and (Rajab et al., 2021). Briefly, the gene expression values from RNA Sequencing and Microarray were transformed into percentile values. Then using a linear mixed model each gene’s variance composition was determined regarding to variance explained by or dependent on the platform. Then a threshold for gene selection was empirically determined based on a platform variance ratio of 0.16 to remove genes with platform effect for the first three principal components. A total of 2417 genes with low platform-dependent variance proportion and high variance explained by the samples’ biology were kept for generating atlas and visualizing by PCA. Codes for constructing atlas are available at Wells Lab GitHub.

### Pseudo-bulk samples from external single-cell dataset

To project external single-cell data on to DC atlas, first single cells were aggregated to build pseudo-bulk samples in order to mitigate library size differences. For each cluster defined by the author of single-cell data, the cells were randomly separated into subgroups of a size 15. Thus, every pseudo-bulk sample was aggregated from 14 samples and the expression values of each aggregated group was a summation of the subgroup’s expression values for each gene.

### Capybara similarity analysis

Capybara (Kong et al. 2020) identity scores were used to measure the similarity of the external pseudo-bulk samples to the reference atlas. Capybara cell scores were calculated by performing a restricted linear regression of atlas samples on each of the pseudo-bulk samples’ expression profiles as described previously by (Rajab et al., 2021).

### Differential expression (DE) analysis

For DE analysis between *in vivo* and *in vitro* models, the linear mixed model fitting from the Lme4 R package was used to estimate the parameters of the formula, including fixed- and random-effects as described by (Bates et al. 2014). For each gene, the fitted model to the expression values includes the variable of interest, sample source, and the platform as the batch variable. The p-values and proportion of explained variance by each parameter were extracted from the model and were used to find the significant differentially expressed genes explained by the sample source (p-value <0.05) with lower batch variance proportion (<0.5). For adjusted p-values, the Benjamini–Hochberg (BH) method was used. p-values recorded for two-group comparisons in violin plots was done using student t-test.

### Enrichment analysis

Gene ontology (GO) biological process enrichment analysis was conducted on the top 200 genes missing from *in vitro*-derived DCs. Genes were ranked by their significance of BH adjusted p-value (<0.5), the difference of mean expression of *in vivo* and *in vitro* cells (>0.2) and the variance explained by sample source (>0.5). Enriched pathways were identified using these genes by ClusterProfile R package 3.12.0 in Bioconductor. The same analysis was conducted on the genes upregulated by *in vitro* DCs.

### Graphing and illustration

Violin plots of ranked gene expressions were generated using Plotly Python Graphing Library. PCA graphs were created through www.stemformatics.org platform. The schematic illustration was created by Biorender.com.

## 1. Supplementary Materials

**Figure S1.**
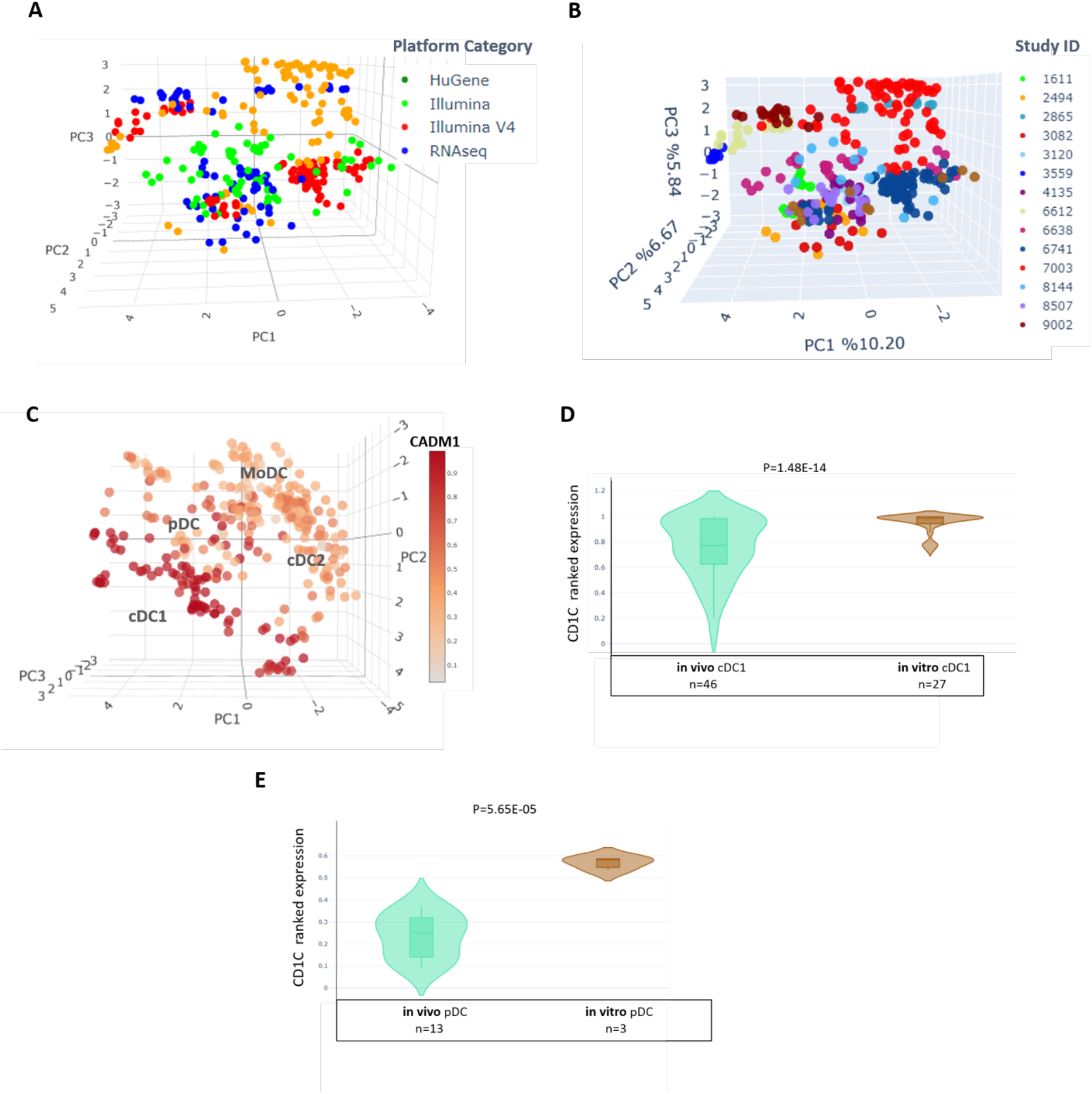
The Dendritic Cell (DC) Atlas assessing DC expression phenotypes across multiple experimental attributes. (A) Stemformatics DC atlas samples distribution across multiple (A) platform categories and (B) individual studies specified by Stemformatics datasets’ IDs. (C) Overlay of *CADM1* expression as a highly expressed marker by cDC1. Colour intensity ranges from high expression (dark red) to low/no expression (pale red). (D) Ranked gene expression (median, interquartile range) of *CD1C*, a marker of putative cDC2, in samples selected for subtype (D) cDC1 and (E) plasmocytoid dendritic cells (pDC) comparing *in vitro*-generated models and their tissue-resident *in vivo* counterparts. Sample sizes (n) are given under each combined category. p value: t-test.

**Figure S2.**
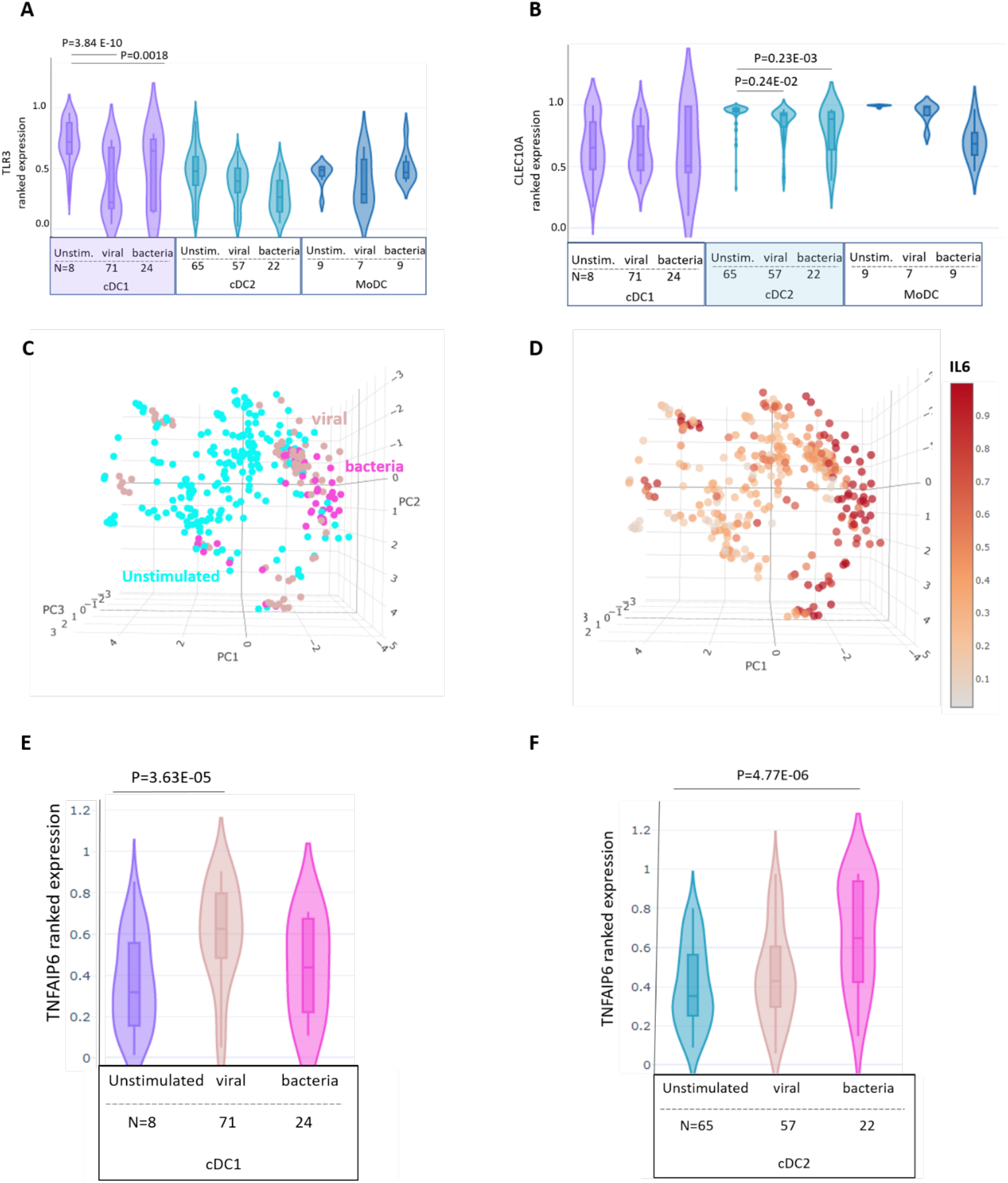
Changes in gene expression of DC subsets as a result of activation. Ranked gene expression (median, interquartile range) of (A) *TLR3*, a marker of putative cDC1 and (B) *CLEC10A*, a marker of putative cDC2, in samples selected for DC subtype cDC1, cDC2 or moDC comparing activation status between bacterial, unstimulated or viral agonists. Sample sizes (N) given under each combine category. (C) Annotation of DC samples by activation highlighting the activation axis along the PC1 in the DC atlas (D) Overlay of *IL6* expression as a highly expressed gene in activated condition. Colour intensity ranges from high expression (dark red) to low/no expression (pale red). (E) Ranked gene expression (median, interquartile range) of *TNFAIP6* for the subsets of (E) cDC1 and (F) cDC2 comparing activation status between bacterial, unstimulated or viral agonists. Sample sizes (N) are given under each combined category. p value: t-test.

**Figure S3:**
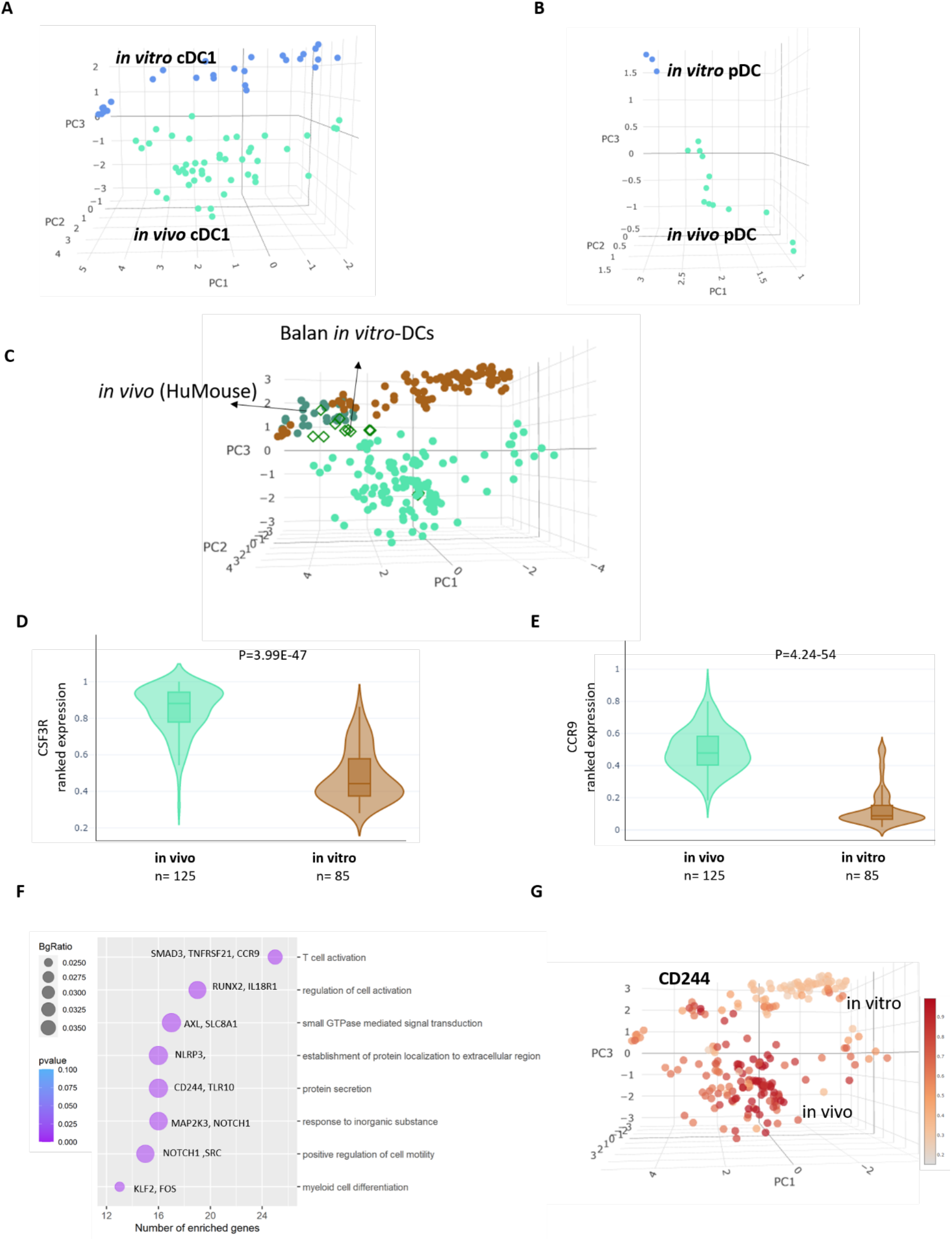
Key factors missing from *in vitro*-derived DCs compared to their *in* vivo counterparts. (A) Stemformatics DC atlas cDC1 subset samples coloured by in vitro (n=27) and in vivo (n=46) DC sources. (B) Stemformatics DC atlas pDC subset samples coloured by in vitro (n=3) and in vivo (n=14) DC sources. (C) Projection of pseudo-bulk samples from single-cell data describing in vitro-derived DC from (Balan et al., 2018) demonstrates similarity of these models with models derived in vivo HuMouse. (D) Ranked gene expression of receptors *CSF3R* and (E) *CCR9* between *in vivo* (n=125) and *in vitro* (n=85) models of DCs. Sample size (n) for each combined category listed under x-axis. (F) Gene set enrichment analysis of GO terms (biological process) lost in *in vitro*-derived DCs. Gene ratio (DE member/GO term membership), p-value adjusted calculated using Benjamini Hoschberg method; Gene symbol of DE genes overlapping GO term. (G) Overlay of ranked gene expression of *CD244* between *in vivo* and *in vitro* samples. Colour intensity ranges from high expression (dark red) to low/no expression (pale red). p value: t-test.

**Supplementary Figure 4:**
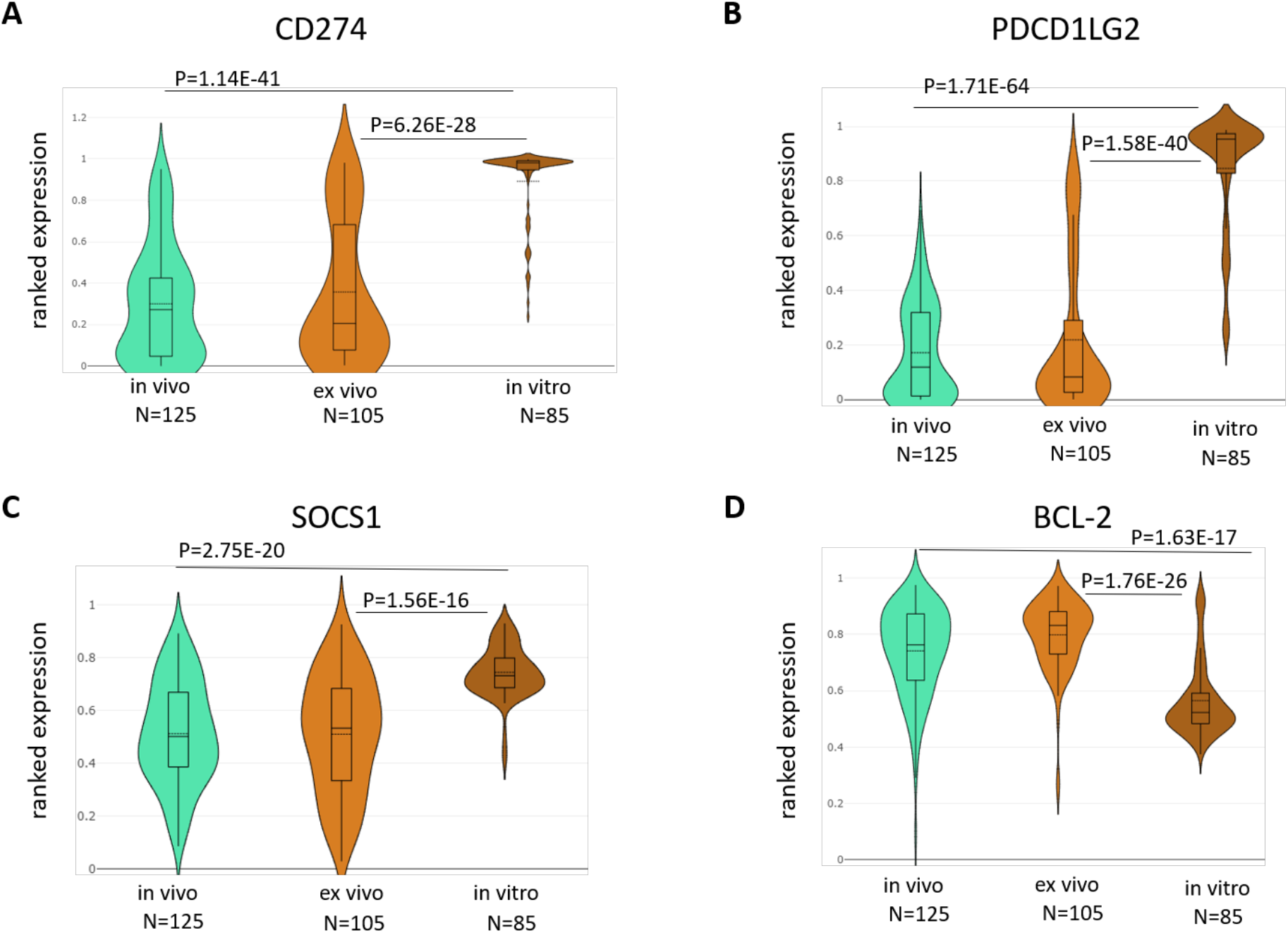
Similar pattern of gene expression between *ex vivo* and *in vivo* samples, and different from *in vitro* models. Ranked gene expression of immunoregulatory markers of (A) *CD274* and (B) *PDCD1LG2* between *in vivo, ex vivo* and *in vitro* samples. (C) Ranked gene expression of SOCS1 gene, downstream from IL-4 signaling pathway, across samples from *in vivo, ex vivo* and *in vitro* sources. (D). Ranked gene expression of antiapoptotic marker, BCL-2, across samples from *in vivo, ex vivo* and *in vitro* sources. Sample sizes (N) are given under each combined category. p value: t-test.

## References

Angel, P.W., Rajab, N., Deng, Y., Pacheco, C.M., Chen, T., Lê Cao, K.A., Choi, J., and Wells, C.A. (2020). A simple, scalable approach to building a cross-platform transcriptome atlas. PLoS Computational Biology 16, e1008219.

Balan, S., Ollion, V., Colletti, N., Chelbi, R., Montanana-Sanchis, F., Liu, H., Vu Manh, T.-P., Sanchez, C., Savoret, J., Perrot, I., et al. (2014). Human XCR1 + Dendritic Cells Derived In Vitro from CD34 + Progenitors Closely Resemble Blood Dendritic Cells, Including Their Adjuvant Responsiveness, Contrary to Monocyte-Derived Dendritic Cells. The Journal of Immunology 193, 1622–1635.

Balan, S., Arnold-Schrauf, C., Abbas, A., Couespel, N., Savoret, J., Imperatore, F., Villani, A.C., Vu Manh, T.P., Bhardwaj, N., and Dalod, M. (2018). Large-Scale Human Dendritic Cell Differentiation Revealing Notch-Dependent Lineage Bifurcation and Heterogeneity. Cell Reports 24, 1902–1915.e6.

Balan, S., Radford, K.J., and Bhardwaj, N. (2020). Unexplored horizons of cDC1 in immunity and tolerance. Advances in Immunology 148, 49–91.

Brown, C.C., Gudjonson, H., Pritykin, Y., Deep, D., Lavallée, V.P., Mendoza, A., Fromme, R., Mazutis, L., Ariyan, C., Leslie, C., et al. (2019). Transcriptional Basis of Mouse and Human Dendritic Cell Heterogeneity. Cell 179, 846–863.e24.

Calmeiro, J., Carrascal, M.A., Tavares, A.R., Ferreira, D.A., Gomes, C., Falcão, A., Cruz, M.T., and Neves, B.M. (2020). Dendritic cell vaccines for cancer immunotherapy: The role of human conventional type 1 dendritic cells. Pharmaceutics 12, 158.

Choi, J., Pacheco, C.M., Mosbergen, R., Korn, O., Chen, T., Nagpal, I., Englart, S., Angel, P.W., and Wells, C.A. (2019). Stemformatics: Visualize and download curated stem cell data. Nucleic Acids Research 47, D841–D846.

Globisch, T., Steiner, N., Fülle, L., Lukacs-Kornek, V., Degrandi, D., Dresing, P., Alferink, J., Lang, R., Pfeffer, K., Beyer, M., et al. (2014). Cytokine-dependent regulation of dendritic cell differentiation in the splenic microenvironment. European Journal of Immunology 44, 500–510.

Haniffa, M., Shin, A., Bigley, V., McGovern, N., Teo, P., See, P., Wasan, P.S., Wang, X.N., Malinarich, F., Malleret, B., et al. (2012). Human Tissues Contain CD141 hi Cross-Presenting Dendritic Cells with Functional Homology to Mouse CD103 + Nonlymphoid Dendritic Cells. Immunity 37, 60–73.

Heidkamp, G.F., Sander, J., Lehmann, C.H.K., Heger, L., Eissing, N., Baranska, A., Lühr, J.J., Hoffmann, A., Reimer, K.C., and Lux, A. (2016). Human lymphoid organ dendritic cell identity is predominantly dictated by ontogeny, not tissue microenvironment. Science Immunology 1, eaai7677.

Heil, F., Hemmi, H., Hochrein, H., Ampenberger, F., Kirschning, C., Akira, S., Lipford, G., Wagner, H., and Bauer, S. (2004). Species-Specific Recognition of Single-Stranded RNA via Till-like Receptor 7 and 8. Science 303, 1526–1529.

Jackson, S.H., Yu, C.-R., Mahdi, R.M., Ebong, S., and Egwuagu, C.E. (2004). Dendritic Cell Maturation Requires STAT1 and Is under Feedback Regulation by Suppressors of Cytokine Signaling. The Journal of Immunology 172, 2307–2315.

Jones, A., Bourque, J., Kuehm, L., Opejin, A., Teague, R.M., Gross, C., and Hawiger, D. (2016). Immunomodulatory Functions of BTLA and HVEM Govern Induction of Extrathymic Regulatory T Cells and Tolerance by Dendritic Cells. Immunity 45, 1066–1077.

Kirkling, M.E., Cytlak, U., Lau, C.M., Lewis, K.L., Resteu, A., Khodadadi-Jamayran, A., Siebel, C.W., Salmon, H., Merad, M., Tsirigos, A., et al. (2018). Notch Signaling Facilitates In Vitro Generation of Cross-Presenting Classical Dendritic Cells. Cell Reports 23, 3658–3672.e6.

Leal Rojas, I.M., Mok, W.H., Pearson, F.E., Minoda, Y., Kenna, T.J., Barnard, R.T., and Radford, K.J. (2017). Human blood cD1c+ dendritic cells promote Th1 and Th17 effector function in memory cD4+ T cells. Frontiers in Immunology 8, 971.

van Leeuwen-Kerkhoff, N., Lundberg, K., Westers, T.M., Kordasti, S., Bontkes, H.J., Lindstedt, M., de Gruijl, T.D., and van de Loosdrecht, A.A. (2018). Human bone marrow-derived myeloid dendritic cells show an immature transcriptional and functional profile compared to their peripheral blood counterparts and separate from slan+ non-classical monocytes. Frontiers in Immunology 9, 1619.

Macri, C., Pang, E.S., Patton, T., and O’Keeffe, M. (2018). Dendritic cell subsets. Seminars in Cell & Developmental Biology 84, 11–21.

McGovern, N., Shin, A., Low, G., Low, D., Duan, K., Yao, L.J., Msallam, R., Low, I., Shadan, N.B., Sumatoh, H.R., et al. (2017). Human fetal dendritic cells promote prenatal T-cell immune suppression through arginase-2. Nature 546, 662–666.

Merad, M., Sathe, P., Helft, J., Miller, J., and Mortha, A. (2013). The dendritic cell lineage: Ontogeny and function of dendritic cells and their subsets in the steady state and the inflamed setting. Annual Review of Immunology 31, 563–604.

Minoda, Y., Virshup, I., Rojas, I.L., Haigh, O., Wong, Y., Miles, J.J., Wells, C.A., and Radford, K.J. (2017). Human CD141+ dendritic cell and CD1c+ dendritic cell undergo concordant early genetic programming after activation in humanized mice in vivo. Frontiers in Immunology 8, 1419.

Monkley, S., Krishnaswamy, J.K., Göransson, M., Clausen, M., Meuller, J., Thörn, K., Hicks, R., Delaney, S., and Stjernborg, L. (2020). Optimised generation of iPSC-derived macrophages and dendritic cells that are functionally and transcriptionally similar to their primary counterparts. PLoS ONE 15, e0243807.

Pacis, A., Tailleux, L., Morin, A.M., Lambourne, J., MacIsaac, J.L., Yotova, V., Dumaine, A., Danckært, A., Luca, F., Grenier, J.C., et al. (2015). Bacterial infection remodels the DNA methylation landscape of human dendritic cells. Genome Research 25, 1801–1811.

Pan, G., Bauer, J.H., Haridas, V., Wang, S., Liu, D., Yu, G., Vincenz, C., Aggarwal, B.B., Ni, J., and Dixit, V.M. (1998). Identification and functional characterization of DR6, a novel death domain-containing TNF receptor. FEBS Letters 431, 351–356.

Pang, E.S., Daraj, G., Balka, K.R., De Nardo, D., Macri, C., Hochrein, H., Masterman, K.-A., Tan, P.S., Shoppee, A., Magill, Z., et al. (2022). Discordance in STING-Induced Activation and Cell Death Between Mouse and Human Dendritic Cell Populations. Frontiers in Immunology 13, 794776–794776.

Peng, W.M., Maintz, L., Allam, J.P., and Novak, N. (2013). Attenuated TGF-β1 responsiveness of dendritic cells and their precursors in atopic dermatitis. European Journal of Immunology 43, 1374–1382.

Poulin, L.F., Salio, M., Griessinger, E., Anjos-Afonso, F., Craciun, L., Chen, J.L., Keller, A.M., Joffre, O., Zelenay, S., Nye, E., et al. (2010). Characterization of human DNGR-1+ BDCA3+ leukocytes as putative equivalents of mouse CD8α+ dendritic cells. Journal of Experimental Medicine 207, 1261–1271.

Rajab, N., Angel, P.W., Deng, Y., Gu, J., Jameson, V., Kurowska-Stolarska, M., Milling, S., Pacheco, C.M., Rutar, M., Laslett, A.L., et al. (2021). An integrated analysis of human myeloid cells identifies gaps in in vitro models of in vivo biology. Stem Cell Reports 16, 1629–1643.

Roquilly, A., Mintern, J.D., and Villadangos, J.A. (2022). Spatiotemporal Adaptations of Macrophage and Dendritic Cell Development and Function. Annual Review of Immunology 40, 525–557.

Rosa, F.F., Pires, C.F., Kurochkin, I., Ferreira, A.G., Gomes, A.M., Palma, L.G., Shaiv, K., Solanas, L., Azenha, C., Papatsenko, D., et al. (2018). Direct reprogramming of fibroblasts into antigen-presenting dendritic cells. Science Immunology 3, eaau4292.

Rosa, F.F., Pires, C.F., Kurochkin, I., Halitzki, E., Zahan, T., Arh, N., Zimmermannová, O., Ferreira, A.G., Li, H., Karlsson, S., et al. (2022). Single-cell transcriptional profiling informs efficient reprogramming of human somatic cells to cross-presenting dendritic cells. Science Immunology 7, eabg5539.

See, P., Dutertre, C.A., Chen, J., Günther, P., McGovern, N., Irac, S.E., Gunawan, M., Beyer, M., Händler, K., Duan, K., et al. (2017). Mapping the human DC lineage through the integration of high-dimensional techniques. Science 356, eaag3009.

De Smedt, T., Pajak, B., Muraille, E., Lespagnard, L., Heinen, E., De Baetselier, P., Urbain, J., Leo, O., and Moser, M. (1996). Regulation of dendritic cell numbers and maturation by lipopolysaccharide in vivo. Journal of Experimental Medicine 184, 1413–1424.

Ushach, I., and Zlotnik, A. (2016). Biological role of granulocyte macrophage colony-stimulating factor (GM-CSF) and macrophage colony-stimulating factor (M-CSF) on cells of the myeloid lineage. Journal of Leukocyte Biology 100, 481–489.

Villani, A.-C., Satija, R., Reynolds, G., Sarkizova, S., Shekhar, K., Fletcher, J., Griesbeck, M., Butler, A., Zheng, S., and Lazo, S. (2017). Single-cell RNA-seq reveals new types of human blood dendritic cells, monocytes, and progenitors. Science 356, eaah4573.

Watchmaker, P.B., Lahl, K., Lee, M., Baumjohann, D., Morton, J., Kim, S.J., Zeng, R., Dent, A., Ansel, K.M., Diamond, B., et al. (2014). Comparative transcriptional and functional profiling defines conserved programs of intestinal DC differentiation in humans and mice. Nature Immunology 15, 98–108.

Wylie, B., Macri, C., Mintern, J.D., and Waithman, J. (2019). Dendritic cells and cancer: From biology to therapeutic intervention. Cancers 11, 521.

