## Supplementary Table 1 for "The Human Dendritic Cell Atlas: An integrated transcriptional tool to study human dendritic cell biology"

**Supplementary Table 1.** List of studies processed for inclusion in the Human Dendritic Cell Atlas and their detailed information

| **Dataset name/**  **PMID code** | **Cell type** | **Sample source** | **Tissue** | **Activation** | **Platform** | **accession code/Inclusion in DC atlas** |
| --- | --- | --- | --- | --- | --- | --- |
| Anselmi G, PMID: 32345968 | DC1, DC2, pDC | *in vivo*  and *in vitro* | blood (*in vivo* samples) | healthy | RNA-seq | GSE144435/not included (failed QC)  GSE145803/ included |
| Balan S, PMID: 25009205 | DC1, MoDC | *in vitro*  and *ex vivo* | *in vitro* differentiated cells | untreated  &  treated | Microarray, Affymetrix (HuGene-1_0-st) | GSE57671/ included |
| BanchereauPMID: 25335753 | DC1, DC2 | *ex vivo* | blood | untreated  &  treated | Microarray Illumina HumanHT-12 V4.0 | GSE56744/ included |
| Haniffa M, PMID: 22795876 | DC1, DC2. pDC, monocyte | *in vivo* | blood skin | healthy | Microarray, Illumina HT12 V4 | GSE35457/ included |
| Helft J, PMID: 28723558 | DC1 | *in vitro* | cord blood | healthy | Microarray, Affymetrix (HuGene-1_0-st) | GSE98957/ included |
| Rojas L, PMID: 28878767 | DC1, DC2 | *in vivo* (humanised mice) | *in vitro* differentiated cells | untreated  &  treated | Microarray,  Illumina HumanHT-12 v4 | GSE99666/ included |
| See P,  PMID: 28473638 | DC1, DC2, pDC, DC precursor | *in vivo* | blood | disease:  PHS syndrome | Microarray,  Illumina HumanHT-12 v4 | GSE80171/ included |
| Silvin A, PMID: 28783704 | DC1, DC2, pDC | *in vivo* | blood | healthy | RNA-seq | GSE76511/ included |
| Ma W, PMID: 31253076 | DC1, DC2, DC precursor | *in vivo* | blood | healthy | RNA-seq | GSE89322 |
| Canavan M, PMID: 30518680 | DC1, DC2 | *in vivo* | synovial tissue, blood | healthy  &  inflamed synovial joint fluid | RNA-seq | GSE108174/ included |
| Gabrielsen ISM,  PMC: 6602790 | DC1, pDC | *in vivo* | thymus | healthy | RNA-seq | included* |
| Yin X, PMID: 28087664 | DC2 | *in vivo* | blood | healthy | RNA-seq | GSE77649 |
| Pacis A, PMID: 26392366 | MoDC | *in vitro* | *in vitro* differentiated cells | treated  &  untreated | RNA-seq | GSE64179 |
| Watchmaker PB,  PMID: 24292363 | DC1  DC2 | *in vivo* | Intestine | healthy | Microarray Affymetrix (HuGene-1_0-st) | GSE50380/ included |
| Bourdely P, PMID: 32610077 | DC1, DC2 | *in vitro* | *in vitro* differentiated cells | untreated | RNA-seq | GSE151095/ not included (failed QC) |
| McGovern N, PMID: 28614294 | DC1, DC2, monocyte | *in vivo* | skin  & spleen | healthy  & disease:  pancreatic tumours | Microarray Illumina HumanHT-12 V4.0 | GSE85305/ included GSE85304/ included |
