## Supplementary Table 2 for "The Human Dendritic Cell Atlas: An integrated transcriptional tool to study human dendritic cell biology"

Supplementary Table 2. Supplementary Table 2: Differentially expressed genes between *in vitro-*generated DCs and in vivo DCs

| ENSEMBL ID | Gene Symbol | Mean ranked expression, *in vitro* DC | Mean ranked expression, *in vivo* DC | p value |
| --- | --- | --- | --- | --- |
| ENSG00000091651 | ORC6 | 0.505831 | 0.744487 | 5.73E-07 |
| ENSG00000251192 | ZNF674 | 0.212612 | 0.721584 | 0.044677 |
| ENSG00000238172 | RPS2P35 | 0.244604 | 0.670279 | 5.42E-21 |
| ENSG00000171241 | SHCBP1 | 0.517182 | 0.773564 | 9.65E-06 |
| ENSG00000237190 | CDKN2AIPNL | 0.52347 | 0.760102 | 2.82E-09 |
| ENSG00000130522 | JUND | 0.782713 | 0.98976 | 8.41E-37 |
| ENSG00000150782 | IL18 | 0.591839 | 0.862867 | 0.000652 |
| ENSG00000026297 | RNASET2 | 0.616916 | 0.961595 | 1.14E-20 |
| ENSG00000236533 | AC009413.1 | 0.469069 | 0.675818 | 0.000565 |
| ENSG00000170421 | KRT8 | 0.260578 | 0.745029 | 4.01E-30 |
| ENSG00000182287 | AP1S2 | 0.666068 | 0.915238 | 0.005838 |
| ENSG00000198155 | ZNF876P | 0.148995 | 0.672194 | 0.02555 |
| ENSG00000228502 | EEF1A1P11 | 0.523722 | 0.737672 | 2.53E-09 |
| ENSG00000177350 | RPL13AP3 | 0.447907 | 0.69756 | 1.75E-11 |
| ENSG00000105650 | PDE4C | 0.429727 | 0.689671 | 7.62E-05 |
| ENSG00000122378 | FAM213A | 0.506214 | 0.742794 | 0.008319 |
| ENSG00000146223 | RPL7L1 | 0.551072 | 0.846482 | 3.20E-07 |
| ENSG00000185163 | DDX51 | 0.530161 | 0.863271 | 7.82E-11 |
| ENSG00000156172 | C8orf37 | 0.318713 | 0.697448 | 8.33E-05 |
| ENSG00000166762 | CATSPER2 | 0.316896 | 0.730652 | 0.001059 |
| ENSG00000197124 | ZNF682 | 0.35367 | 0.702137 | 1.14E-09 |
| ENSG00000226836 | AC018437.1 | 0.186272 | 0.623014 | 1.46E-13 |
| ENSG00000178074 | C2orf69 | 0.621787 | 0.869035 | 1.59E-18 |
| ENSG00000183287 | CCBE1 | 0.121932 | 0.608239 | 0.000493 |
| ENSG00000243779 | AP001086.1 | 0.453863 | 0.679297 | 3.89E-12 |
| ENSG00000178460 | MCMDC2 | 0.156614 | 0.751291 | 4.30E-06 |
| ENSG00000125885 | MCM8 | 0.42114 | 0.791342 | 8.15E-07 |
| ENSG00000179709 | NLRP8 | 0.110031 | 0.599951 | 7.48E-06 |
| ENSG00000163283 | ALPP | 0.206717 | 0.638441 | 3.33E-06 |
| ENSG00000156515 | HK1 | 0.643473 | 0.916825 | 0.020267 |
| ENSG00000172530 | BANP | 0.614089 | 0.869716 | 5.36E-13 |
| ENSG00000144655 | CSRNP1 | 0.673828 | 0.975642 | 3.80E-36 |
| ENSG00000196502 | SULT1A1 | 0.556783 | 0.805058 | 0.002141 |
| ENSG00000180172 | RPS12P23 | 0.070433 | 0.62567 | 0.001556 |
| ENSG00000142609 | CFAP74 | 0.269438 | 0.606188 | 2.32E-05 |
| ENSG00000186889 | TMEM17 | 0.15414 | 0.635132 | 2.33E-05 |
| ENSG00000198740 | ZNF652 | 0.722595 | 0.923937 | 7.70E-11 |
| ENSG00000139433 | GLTP | 0.660445 | 0.877448 | 1.01E-12 |
| ENSG00000106144 | CASP2 | 0.696827 | 0.915119 | 3.06E-23 |
| ENSG00000063244 | U2AF2 | 0.671457 | 0.929258 | 1.08E-27 |
| ENSG00000170852 | KBTBD2 | 0.698743 | 0.902733 | 3.83E-07 |
| ENSG00000186063 | AIDA | 0.457386 | 0.759155 | 8.01E-07 |
| ENSG00000157514 | TSC22D3 | 0.478218 | 0.962877 | 1.97E-54 |
| ENSG00000182022 | CHST15 | 0.508204 | 0.797857 | 1.43E-06 |
| ENSG00000144802 | NFKBIZ | 0.699123 | 0.95484 | 2.40E-13 |
| ENSG00000034152 | MAP2K3 | 0.679483 | 0.935083 | 6.82E-33 |
| ENSG00000144730 | IL17RD | 0.230464 | 0.612733 | 0.004257 |
| ENSG00000172216 | CEBPB | 0.589672 | 0.845179 | 0.000585 |
| ENSG00000151881 | TMEM267 | 0.38913 | 0.729256 | 3.06E-10 |
| ENSG00000100034 | PPM1F | 0.660348 | 0.911449 | 1.44E-11 |
| ENSG00000087206 | UIMC1 | 0.571123 | 0.864755 | 4.98E-10 |
| ENSG00000164663 | USP49 | 0.343723 | 0.776714 | 8.86E-12 |
| ENSG00000143977 | SNRPG | 0.512457 | 0.762469 | 0.026268 |
| ENSG00000129667 | RHBDF2 | 0.506945 | 0.845683 | 0.006496 |
| ENSG00000103168 | TAF1C | 0.499297 | 0.844171 | 2.31E-13 |
| ENSG00000177169 | ULK1 | 0.650806 | 0.870336 | 4.53E-05 |
| ENSG00000028310 | BRD9 | 0.624246 | 0.87222 | 1.07E-06 |
| ENSG00000160298 | C21orf58 | 0.40934 | 0.678093 | 8.59E-11 |
| ENSG00000197912 | SPG7 | 0.673702 | 0.880983 | 0.040355 |
| ENSG00000184588 | PDE4B | 0.675849 | 0.936722 | 6.86E-12 |
| ENSG00000078177 | N4BP2 | 0.695585 | 0.896187 | 1.13E-06 |
| ENSG00000063245 | EPN1 | 0.657665 | 0.901806 | 2.95E-33 |
| ENSG00000079739 | PGM1 | 0.579377 | 0.834351 | 0.045805 |
| ENSG00000196584 | XRCC2 | 0.247589 | 0.63579 | 0.000431 |
| ENSG00000170381 | SEMA3E | 0.067411 | 0.565101 | 0.006703 |
| ENSG00000131876 | SNRPA1 | 0.458402 | 0.826875 | 0.001894 |
| ENSG00000162775 | RBM15 | 0.640049 | 0.878806 | 2.92E-41 |
| ENSG00000105122 | RASAL3 | 0.559519 | 0.847266 | 1.13E-17 |
| ENSG00000162711 | NLRP3 | 0.520724 | 0.902839 | 2.29E-25 |
| ENSG00000166501 | PRKCB | 0.682481 | 0.925032 | 7.22E-23 |
| ENSG00000126391 | FRMD8 | 0.538123 | 0.825709 | 0.032473 |
| ENSG00000269335 | IKBKG | 0.381621 | 0.750458 | 0.000228 |
| ENSG00000160299 | PCNT | 0.623551 | 0.882605 | 0.002349 |
| ENSG00000099904 | ZDHHC8 | 0.5615 | 0.825818 | 1.14E-09 |
| ENSG00000127528 | KLF2 | 0.520177 | 0.846271 | 1.76E-39 |
| ENSG00000160584 | SIK3 | 0.676146 | 0.892861 | 4.24E-36 |
| ENSG00000159086 | PAXBP1 | 0.634976 | 0.87154 | 2.52E-05 |
| ENSG00000149499 | EML3 | 0.614896 | 0.869322 | 3.23E-13 |
| ENSG00000061938 | TNK2 | 0.662027 | 0.9104 | 2.38E-23 |
| ENSG00000122490 | PQLC1 | 0.661341 | 0.888104 | 6.37E-13 |
| ENSG00000138439 | FAM117B | 0.613067 | 0.844711 | 1.26E-19 |
| ENSG00000170525 | PFKFB3 | 0.738533 | 0.943736 | 3.73E-17 |
| ENSG00000135372 | NAT10 | 0.590596 | 0.850396 | 6.68E-10 |
| ENSG00000180448 | ARHGAP45 | 0.700383 | 0.945323 | 1.90E-17 |
| ENSG00000136861 | CDK5RAP2 | 0.603564 | 0.85541 | 1.78E-09 |
| ENSG00000097007 | ABL1 | 0.700433 | 0.912532 | 2.51E-21 |
| ENSG00000106003 | LFNG | 0.413238 | 0.82046 | 6.40E-23 |
| ENSG00000085265 | FCN1 | 0.565025 | 0.767823 | 0.008588 |
| ENSG00000156508 | EEF1A1 | 0.547362 | 0.902572 | 1.90E-17 |
| ENSG00000135924 | DNAJB2 | 0.576771 | 0.826995 | 2.65E-07 |
| ENSG00000227747 | AC096631.2 | 0.342733 | 0.6049 | 2.03E-08 |
| ENSG00000197362 | ZNF786 | 0.373433 | 0.665917 | 5.73E-08 |
| ENSG00000172322 | CLEC12A | 0.357648 | 0.762303 | 5.99E-06 |
| ENSG00000225405 | RPS15AP17 | 0.379798 | 0.606384 | 0.000767 |
| ENSG00000169710 | FASN | 0.645172 | 0.884169 | 1.01E-12 |
| ENSG00000125735 | TNFSF14 | 0.373725 | 0.721903 | 1.06E-14 |
| ENSG00000170345 | FOS | 0.724756 | 0.959066 | 4.15E-23 |
| ENSG00000185504 | FAAP100 | 0.559478 | 0.831087 | 4.78E-13 |
| ENSG00000140941 | MAP1LC3B | 0.616146 | 0.887279 | 3.68E-10 |
| ENSG00000165802 | NSMF | 0.668485 | 0.887868 | 1.88E-14 |
| ENSG00000160796 | NBEAL2 | 0.706102 | 0.920998 | 5.50E-19 |
| ENSG00000003147 | ICA1 | 0.293209 | 0.663531 | 0.029814 |
| ENSG00000010810 | FYN | 0.638017 | 0.889913 | 3.41E-12 |
| ENSG00000070501 | POLB | 0.623065 | 0.824694 | 3.52E-10 |
| ENSG00000158315 | RHBDL2 | 0.376109 | 0.609023 | 4.39E-07 |
| ENSG00000111913 | RIPOR2 | 0.477582 | 0.8876 | 3.01E-17 |
| ENSG00000137070 | IL11RA | 0.515277 | 0.783885 | 0.000114 |
| ENSG00000142409 | ZNF787 | 0.628384 | 0.840452 | 3.82E-18 |
| ENSG00000184216 | IRAK1 | 0.675792 | 0.901033 | 9.22E-11 |
| ENSG00000100241 | SBF1 | 0.645689 | 0.898066 | 7.85E-17 |
| ENSG00000134086 | VHL | 0.637962 | 0.854299 | 4.43E-11 |
| ENSG00000133030 | MPRIP | 0.508284 | 0.899 | 2.37E-37 |
| ENSG00000163482 | STK36 | 0.411743 | 0.725485 | 0.023691 |
| ENSG00000125740 | FOSB | 0.518773 | 0.961409 | 1.91E-65 |
| ENSG00000179409 | GEMIN4 | 0.525127 | 0.771401 | 0.000265 |
| ENSG00000100814 | CCNB1IP1 | 0.492368 | 0.822325 | 8.61E-08 |
| ENSG00000167967 | E4F1 | 0.601727 | 0.81402 | 0.00224 |
| ENSG00000179335 | CLK3 | 0.673666 | 0.893135 | 1.18E-13 |
| ENSG00000126775 | ATG14 | 0.606278 | 0.828761 | 5.23E-08 |
| ENSG00000105643 | ARRDC2 | 0.541604 | 0.881813 | 2.04E-25 |
| ENSG00000090376 | IRAK3 | 0.47056 | 0.772457 | 0.017662 |
| ENSG00000119138 | KLF9 | 0.657261 | 0.859873 | 1.17E-33 |
| ENSG00000148400 | NOTCH1 | 0.630568 | 0.831121 | 2.24E-07 |
| ENSG00000153574 | RPIA | 0.553943 | 0.7611 | 0.015944 |
| ENSG00000073756 | PTGS2 | 0.532428 | 0.785306 | 7.13E-09 |
| ENSG00000135211 | TMEM60 | 0.861082 | 0.62385 | 1.18E-38 |
| ENSG00000181444 | ZNF467 | 0.594839 | 0.817029 | 1.91E-07 |
| ENSG00000168329 | CX3CR1 | 0.241757 | 0.617438 | 0.000498 |
| ENSG00000109670 | FBXW7 | 0.546077 | 0.869372 | 6.76E-28 |
| ENSG00000130052 | STARD8 | 0.452014 | 0.788802 | 1.51E-07 |
| ENSG00000128284 | APOL3 | 0.508007 | 0.796146 | 0.035347 |
| ENSG00000198612 | COPS8 | 0.423142 | 0.793114 | 1.02E-07 |
| ENSG00000152348 | ATG10 | 0.46373 | 0.810167 | 9.98E-07 |
| ENSG00000152242 | C18orf25 | 0.583866 | 0.80964 | 8.35E-05 |
| ENSG00000168404 | MLKL | 0.463975 | 0.676064 | 0.024189 |
| ENSG00000100918 | REC8 | 0.500132 | 0.871172 | 4.18E-35 |
| ENSG00000175489 | LRRC25 | 0.577587 | 0.807832 | 2.46E-09 |
| ENSG00000148841 | ITPRIP | 0.656024 | 0.881475 | 1.28E-35 |
| ENSG00000083457 | ITGAE | 0.472746 | 0.765467 | 0.006255 |
| ENSG00000134215 | VAV3 | 0.628862 | 0.842195 | 3.25E-08 |
| ENSG00000180354 | MTURN | 0.515345 | 0.776083 | 0.003581 |
| ENSG00000131381 | RBSN | 0.407173 | 0.761379 | 0.002456 |
| ENSG00000154930 | ACSS1 | 0.510664 | 0.808285 | 8.07E-15 |
| ENSG00000156925 | ZIC3 | 0.18486 | 0.544633 | 0.005458 |
| ENSG00000182162 | P2RY8 | 0.628001 | 0.83948 | 0.000106 |
| ENSG00000160613 | PCSK7 | 0.639818 | 0.882322 | 1.29E-29 |
| ENSG00000169252 | ADRB2 | 0.488746 | 0.689656 | 3.79E-05 |
| ENSG00000169398 | PTK2 | 0.588985 | 0.812869 | 0.013283 |
| ENSG00000142864 | SERBP1 | 0.536639 | 0.865003 | 1.76E-08 |
| ENSG00000138413 | IDH1 | 0.927227 | 0.725713 | 1.43E-23 |
| ENSG00000100068 | LRP5L | 0.526813 | 0.774385 | 5.33E-07 |
| ENSG00000228929 | RPS13P2 | 0.51679 | 0.731876 | 0.003289 |
| ENSG00000119866 | BCL11A | 0.659669 | 0.86542 | 1.76E-06 |
| ENSG00000186940 | CHCHD2P9 | 0.205305 | 0.58877 | 0.000289 |
| ENSG00000099326 | MZF1 | 0.516005 | 0.781954 | 0.045432 |
| ENSG00000163050 | COQ8A | 0.547899 | 0.772713 | 0.001213 |
| ENSG00000088247 | KHSRP | 0.686751 | 0.8913 | 6.47E-34 |
| ENSG00000132952 | USPL1 | 0.654023 | 0.855884 | 9.87E-18 |
| ENSG00000128050 | PAICS | 0.533391 | 0.790297 | 0.000161 |
| ENSG00000189403 | HMGB1 | 0.532764 | 0.806362 | 2.20E-16 |
| ENSG00000146350 | TBC1D32 | 0.174498 | 0.661116 | 0.001124 |
| ENSG00000063169 | BICRA | 0.557148 | 0.799959 | 1.95E-20 |
| ENSG00000198873 | GRK5 | 0.331189 | 0.641255 | 5.42E-10 |
| ENSG00000146409 | SLC18B1 | 0.344953 | 0.779559 | 6.25E-09 |
| ENSG00000005844 | ITGAL | 0.592654 | 0.873796 | 1.31E-05 |
| ENSG00000140044 | JDP2 | 0.604833 | 0.805527 | 9.37E-14 |
| ENSG00000139679 | LPAR6 | 0.903133 | 0.670954 | 0.000169 |
| ENSG00000164880 | INTS1 | 0.660316 | 0.866141 | 6.74E-19 |
| ENSG00000105339 | DENND3 | 0.551672 | 0.807251 | 0.00016 |
| ENSG00000090238 | YPEL3 | 0.571896 | 0.780772 | 0.004085 |
| ENSG00000179604 | CDC42EP4 | 0.501314 | 0.735306 | 2.45E-08 |
| ENSG00000141401 | IMPA2 | 0.522662 | 0.790941 | 1.16E-11 |
| ENSG00000139197 | PEX5 | 0.426956 | 0.715824 | 5.39E-07 |
| ENSG00000181722 | ZBTB20 | 0.5114 | 0.802057 | 7.72E-19 |
| ENSG00000091490 | SEL1L3 | 0.626698 | 0.887081 | 1.46E-08 |
| ENSG00000041880 | PARP3 | 0.478897 | 0.732326 | 6.69E-07 |
| ENSG00000136717 | BIN1 | 0.631878 | 0.90369 | 8.31E-22 |
| ENSG00000172716 | SLFN11 | 0.480006 | 0.775952 | 5.32E-08 |
| ENSG00000109944 | JHY | 0.445445 | 0.702241 | 1.39E-12 |
| ENSG00000163820 | FYCO1 | 0.482234 | 0.712476 | 1.15E-05 |
| ENSG00000112242 | E2F3 | 0.546924 | 0.756859 | 4.02E-05 |
| ENSG00000082074 | FYB1 | 0.575047 | 0.800816 | 7.01E-07 |
| ENSG00000196844 | PATE2 | 0.190769 | 0.501944 | 0.043947 |
| ENSG00000175857 | GAPT | 0.34281 | 0.662223 | 7.32E-05 |
| ENSG00000089820 | ARHGAP4 | 0.670619 | 0.910232 | 8.63E-19 |
| ENSG00000225787 | LSM3P3 | 0.25887 | 0.578881 | 1.52E-05 |
| ENSG00000141577 | CEP131 | 0.482335 | 0.814582 | 4.78E-36 |
| ENSG00000240370 | RPL13P5 | 0.440108 | 0.674621 | 0.0091 |
| ENSG00000121895 | TMEM156 | 0.216908 | 0.733125 | 4.71E-09 |
| ENSG00000143344 | RGL1 | 0.871971 | 0.615046 | 6.10E-10 |
| ENSG00000121671 | CRY2 | 0.547615 | 0.813145 | 5.18E-19 |
| ENSG00000182378 | PLCXD1 | 0.387353 | 0.746087 | 1.62E-09 |
| ENSG00000125089 | SH3TC1 | 0.652868 | 0.894702 | 9.42E-17 |
| ENSG00000165355 | FBXO33 | 0.56908 | 0.811939 | 1.96E-17 |
| ENSG00000140403 | DNAJA4 | 0.573821 | 0.785612 | 1.35E-11 |
| ENSG00000176438 | SYNE3 | 0.553687 | 0.803038 | 9.68E-09 |
| ENSG00000091592 | NLRP1 | 0.573287 | 0.822632 | 0.0237 |
| ENSG00000132432 | SEC61G | 0.948022 | 0.714827 | 1.60E-31 |
| ENSG00000135473 | PAN2 | 0.534813 | 0.782461 | 2.34E-05 |
| ENSG00000108773 | KAT2A | 0.614946 | 0.840646 | 4.65E-10 |
| ENSG00000163964 | PIGX | 0.537968 | 0.747502 | 0.000756 |
| ENSG00000250510 | GPR162 | 0.418541 | 0.648 | 0.000933 |
| ENSG00000163545 | NUAK2 | 0.511403 | 0.794323 | 9.90E-13 |
| ENSG00000144283 | PKP4 | 0.531003 | 0.740722 | 9.85E-08 |
| ENSG00000188404 | SELL | 0.457851 | 0.850439 | 2.12E-11 |
| ENSG00000147394 | ZNF185 | 0.296376 | 0.622209 | 0.001611 |
| ENSG00000114812 | VIPR1 | 0.378454 | 0.666942 | 7.17E-12 |
| ENSG00000164010 | ERMAP | 0.478676 | 0.691205 | 3.66E-06 |
| ENSG00000101247 | NDUFAF5 | 0.370747 | 0.701451 | 0.047621 |
| ENSG00000196459 | TRAPPC2 | 0.376256 | 0.683048 | 0.002804 |
| ENSG00000135503 | ACVR1B | 0.457642 | 0.699078 | 6.38E-11 |
| ENSG00000132326 | PER2 | 0.561138 | 0.796909 | 8.43E-28 |
| ENSG00000159788 | RGS12 | 0.520666 | 0.768713 | 0.00186 |
| ENSG00000167207 | NOD2 | 0.538242 | 0.744906 | 0.003087 |
| ENSG00000103126 | AXIN1 | 0.550099 | 0.781299 | 3.56E-08 |
| ENSG00000119535 | CSF3R | 0.483367 | 0.842336 | 7.27E-21 |
| ENSG00000104907 | TRMT1 | 0.517292 | 0.787966 | 1.79E-11 |
| ENSG00000144589 | STK11IP | 0.590505 | 0.814247 | 0.000674 |
| ENSG00000197021 | CXorf40B | 0.551755 | 0.753009 | 0.001582 |
| ENSG00000132005 | RFX1 | 0.485562 | 0.718579 | 3.72E-09 |
| ENSG00000107902 | LHPP | 0.497398 | 0.748539 | 9.14E-21 |
| ENSG00000103264 | FBXO31 | 0.534682 | 0.763883 | 4.83E-05 |
| ENSG00000078081 | LAMP3 | 0.922162 | 0.709925 | 1.69E-08 |
| ENSG00000165195 | PIGA | 0.384243 | 0.748189 | 2.85E-16 |
| ENSG00000077984 | CST7 | 0.96602 | 0.751266 | 4.84E-06 |
| ENSG00000127922 | SEM1 | 0.94896 | 0.741935 | 7.82E-13 |
| ENSG00000181631 | P2RY13 | 0.422743 | 0.705496 | 1.23E-10 |
| ENSG00000010803 | SCMH1 | 0.472725 | 0.769501 | 2.65E-14 |
| ENSG00000111639 | MRPL51 | 0.958405 | 0.710259 | 8.96E-19 |
| ENSG00000014914 | MTMR11 | 0.35434 | 0.556071 | 0.022792 |
| ENSG00000142459 | EVI5L | 0.434939 | 0.701487 | 5.06E-12 |
| ENSG00000185187 | SIGIRR | 0.50948 | 0.759424 | 1.01E-14 |
| ENSG00000108515 | ENO3 | 0.600588 | 0.827827 | 2.33E-07 |
| ENSG00000146063 | TRIM41 | 0.40303 | 0.758617 | 4.80E-13 |
| ENSG00000146072 | TNFRSF21 | 0.573848 | 0.853303 | 1.40E-11 |
| ENSG00000198945 | L3MBTL3 | 0.49296 | 0.704608 | 8.57E-06 |
| ENSG00000047578 | KIAA0556 | 0.520274 | 0.755931 | 5.96E-16 |
| ENSG00000145287 | PLAC8 | 0.425752 | 0.746373 | 5.15E-10 |
| ENSG00000150990 | DHX37 | 0.516688 | 0.774518 | 8.17E-18 |
| ENSG00000101220 | C20orf27 | 0.531297 | 0.779878 | 0.000517 |
| ENSG00000164674 | SYTL3 | 0.467873 | 0.700188 | 0.002276 |
| ENSG00000165359 | INTS6L | 0.536713 | 0.777241 | 1.16E-18 |
| ENSG00000027075 | PRKCH | 0.372654 | 0.758773 | 3.59E-21 |
| ENSG00000076641 | PAG1 | 0.468777 | 0.754682 | 1.10E-14 |
| ENSG00000026950 | BTN3A1 | 0.580572 | 0.792816 | 7.19E-15 |
| ENSG00000186376 | ZNF75D | 0.38814 | 0.6789 | 2.18E-05 |
| ENSG00000153234 | NR4A2 | 0.451085 | 0.876913 | 3.01E-39 |
| ENSG00000184428 | TOP1MT | 0.621935 | 0.838024 | 3.40E-11 |
| ENSG00000101544 | ADNP2 | 0.56766 | 0.790346 | 3.70E-18 |
| ENSG00000114841 | DNAH1 | 0.461345 | 0.794464 | 1.19E-25 |
| ENSG00000186866 | POFUT2 | 0.491161 | 0.720804 | 1.76E-06 |
| ENSG00000173852 | DPY19L1 | 0.379926 | 0.669674 | 1.64E-09 |
| ENSG00000021762 | OSBPL5 | 0.415808 | 0.684693 | 0.0011 |
| ENSG00000140090 | SLC24A4 | 0.283007 | 0.747582 | 1.02E-20 |
| ENSG00000116815 | CD58 | 0.922813 | 0.708588 | 4.52E-06 |
| ENSG00000138834 | MAPK8IP3 | 0.607227 | 0.865221 | 3.29E-21 |
| ENSG00000166839 | ANKDD1A | 0.486372 | 0.747768 | 1.35E-12 |
| ENSG00000002933 | TMEM176A | 0.827312 | 0.523854 | 3.42E-07 |
| ENSG00000130147 | SH3BP4 | 0.523849 | 0.832254 | 1.37E-11 |
| ENSG00000166949 | SMAD3 | 0.5806 | 0.812363 | 3.87E-14 |
| ENSG00000111859 | NEDD9 | 0.402305 | 0.771464 | 1.47E-21 |
| ENSG00000162302 | RPS6KA4 | 0.609107 | 0.825877 | 2.86E-11 |
| ENSG00000134571 | MYBPC3 | 0.431708 | 0.65678 | 2.58E-06 |
| ENSG00000167280 | ENGASE | 0.52921 | 0.794747 | 2.11E-14 |
| ENSG00000173917 | HOXB2 | 0.344205 | 0.639214 | 2.98E-06 |
| ENSG00000106804 | C5 | 0.23828 | 0.643965 | 2.01E-06 |
| ENSG00000179981 | TSHZ1 | 0.502375 | 0.745693 | 1.58E-17 |
| ENSG00000196664 | TLR7 | 0.232065 | 0.549518 | 0.001434 |
| ENSG00000196693 | ZNF33B | 0.431015 | 0.661506 | 0.000211 |
| ENSG00000225185 | PPIAP8 | 0.123441 | 0.504417 | 0.000203 |
| ENSG00000125356 | NDUFA1 | 0.981325 | 0.765801 | 1.38E-26 |
| ENSG00000170006 | TMEM154 | 0.495133 | 0.748673 | 6.33E-15 |
| ENSG00000115604 | IL18R1 | 0.40978 | 0.800998 | 4.58E-10 |
| ENSG00000184481 | FOXO4 | 0.433175 | 0.657604 | 3.61E-10 |
| ENSG00000100532 | CGRRF1 | 0.407001 | 0.614458 | 0.030848 |
| ENSG00000127530 | OR7C1 | 0.205054 | 0.542323 | 0.012274 |
| ENSG00000184076 | UQCR10 | 0.920909 | 0.703945 | 1.06E-15 |
| ENSG00000167283 | ATP5MG | 0.978751 | 0.775878 | 8.85E-05 |
| ENSG00000178449 | COX14 | 0.930356 | 0.692194 | 2.72E-39 |
| ENSG00000183828 | NUDT14 | 0.794836 | 0.588224 | 5.07E-16 |
| ENSG00000140259 | MFAP1 | 0.841408 | 0.616388 | 1.65E-31 |
| ENSG00000167601 | AXL | 0.405951 | 0.827961 | 1.13E-20 |
| ENSG00000012232 | EXTL3 | 0.468605 | 0.717025 | 9.06E-13 |
| ENSG00000134590 | RTL8C | 0.909536 | 0.674012 | 1.24E-31 |
| ENSG00000169241 | SLC50A1 | 0.911591 | 0.693767 | 2.07E-20 |
| ENSG00000114023 | FAM162A | 0.908117 | 0.700698 | 1.53E-30 |
| ENSG00000128203 | ASPHD2 | 0.486028 | 0.72938 | 5.05E-07 |
| ENSG00000187210 | GCNT1 | 0.855442 | 0.643475 | 9.85E-06 |
| ENSG00000065518 | NDUFB4 | 0.946275 | 0.731792 | 1.31E-16 |
| ENSG00000181704 | YIPF6 | 0.823951 | 0.608862 | 0.004342 |
| ENSG00000162654 | GBP4 | 0.463528 | 0.669183 | 0.000187 |
| ENSG00000111269 | CREBL2 | 0.928705 | 0.725507 | 8.31E-05 |
| ENSG00000196154 | S100A4 | 0.96681 | 0.705143 | 1.92E-06 |
| ENSG00000139055 | ERP27 | 0.26977 | 0.585768 | 7.66E-07 |
| ENSG00000198105 | ZNF248 | 0.294185 | 0.589873 | 0.015517 |
| ENSG00000176533 | GNG7 | 0.501254 | 0.821199 | 2.33E-15 |
| ENSG00000167612 | ANKRD33 | 0.233251 | 0.462825 | 0.024172 |
| ENSG00000166825 | ANPEP | 0.97155 | 0.76897 | 0.02145 |
| ENSG00000163913 | IFT122 | 0.408302 | 0.668005 | 9.13E-06 |
| ENSG00000100483 | VCPKMT | 0.380127 | 0.698341 | 2.17E-16 |
| ENSG00000197530 | MIB2 | 0.517418 | 0.740329 | 4.32E-11 |
| ENSG00000158321 | AUTS2 | 0.529546 | 0.766779 | 2.71E-06 |
| ENSG00000144504 | ANKMY1 | 0.463206 | 0.713695 | 3.67E-07 |
| ENSG00000035115 | SH3YL1 | 0.295598 | 0.636563 | 1.50E-08 |
| ENSG00000175198 | PCCA | 0.35355 | 0.628483 | 0.012627 |
| ENSG00000132182 | NUP210 | 0.574677 | 0.849277 | 1.15E-11 |
| ENSG00000169504 | CLIC4 | 0.931244 | 0.705042 | 0.01441 |
| ENSG00000125482 | TTF1 | 0.388681 | 0.611539 | 0.004489 |
| ENSG00000107736 | CDH23 | 0.428222 | 0.760598 | 1.44E-09 |
| ENSG00000082781 | ITGB5 | 0.452649 | 0.654142 | 1.90E-05 |
| ENSG00000137103 | TMEM8B | 0.435443 | 0.729737 | 4.15E-06 |
| ENSG00000281005 | LINC00921 | 0.306505 | 0.543359 | 7.78E-05 |
| ENSG00000143224 | PPOX | 0.415598 | 0.623023 | 0.002101 |
| ENSG00000169891 | REPS2 | 0.549348 | 0.783825 | 2.23E-15 |
| ENSG00000148660 | CAMK2G | 0.582647 | 0.806419 | 1.54E-20 |
| ENSG00000139946 | PELI2 | 0.442623 | 0.662448 | 2.00E-12 |
| ENSG00000004961 | HCCS | 0.871033 | 0.621961 | 1.01E-19 |
| ENSG00000179361 | ARID3B | 0.531273 | 0.732893 | 2.75E-06 |
| ENSG00000158481 | CD1C | 0.937862 | 0.700743 | 4.10E-05 |
| ENSG00000171208 | NETO2 | 0.756575 | 0.544966 | 0.022889 |
| ENSG00000137996 | RTCA | 0.900232 | 0.681815 | 3.23E-07 |
| ENSG00000189159 | JPT1 | 0.949899 | 0.698991 | 4.27E-09 |
| ENSG00000110768 | GTF2H1 | 0.855234 | 0.623901 | 5.76E-09 |
| ENSG00000120963 | ZNF706 | 0.495288 | 0.698022 | 0.006106 |
| ENSG00000183508 | FAM46C | 0.455811 | 0.789432 | 1.41E-20 |
| ENSG00000139974 | SLC38A6 | 0.740538 | 0.49313 | 1.50E-07 |
| ENSG00000100100 | PIK3IP1 | 0.395671 | 0.657022 | 2.04E-05 |
| ENSG00000103549 | RNF40 | 0.584538 | 0.790186 | 2.61E-15 |
| ENSG00000138764 | CCNG2 | 0.939411 | 0.69309 | 0.000257 |
| ENSG00000173207 | CKS1B | 0.80162 | 0.592538 | 4.38E-08 |
| ENSG00000071794 | HLTF | 0.473991 | 0.689767 | 6.92E-05 |
| ENSG00000172766 | NAA16 | 0.457454 | 0.689156 | 1.16E-12 |
| ENSG00000107719 | PALD1 | 0.496976 | 0.746846 | 1.59E-07 |
| ENSG00000187554 | TLR5 | 0.215901 | 0.495448 | 0.000799 |
| ENSG00000108405 | P2RX1 | 0.474113 | 0.72959 | 6.28E-06 |
| ENSG00000079263 | SP140 | 0.462348 | 0.722795 | 1.30E-08 |
| ENSG00000075142 | SRI | 0.9131 | 0.698471 | 1.80E-05 |
| ENSG00000158477 | CD1A | 0.832948 | 0.511563 | 1.84E-08 |
| ENSG00000118507 | AKAP7 | 0.217465 | 0.537912 | 0.001173 |
| ENSG00000173409 | ARV1 | 0.72604 | 0.509909 | 0.001561 |
| ENSG00000182636 | NDN | 0.238637 | 0.535836 | 0.038445 |
| ENSG00000015133 | CCDC88C | 0.486742 | 0.745481 | 4.41E-19 |
| ENSG00000161980 | POLR3K | 0.884956 | 0.612498 | 3.37E-12 |
| ENSG00000185085 | INTS5 | 0.834805 | 0.61165 | 1.74E-11 |
| ENSG00000254004 | ZNF260 | 0.804809 | 0.513495 | 1.88E-11 |
| ENSG00000132359 | RAP1GAP2 | 0.471863 | 0.75449 | 3.07E-24 |
| ENSG00000183023 | SLC8A1 | 0.484487 | 0.691961 | 2.79E-10 |
| ENSG00000151136 | BTBD11 | 0.188968 | 0.513859 | 0.000383 |
| ENSG00000146070 | PLA2G7 | 0.779279 | 0.457216 | 7.32E-05 |
| ENSG00000138050 | THUMPD2 | 0.430448 | 0.678937 | 0.009583 |
| ENSG00000127838 | PNKD | 0.718718 | 0.491565 | 3.92E-05 |
| ENSG00000138756 | BMP2K | 0.943523 | 0.739688 | 9.00E-09 |
| ENSG00000102962 | CCL22 | 0.965235 | 0.465545 | 2.45E-07 |
| ENSG00000115137 | DNAJC27 | 0.365899 | 0.638523 | 0.000197 |
| ENSG00000149639 | SOGA1 | 0.559218 | 0.792568 | 9.48E-11 |
| ENSG00000137710 | RDX | 0.883724 | 0.671334 | 0.015154 |
| ENSG00000164167 | LSM6 | 0.860354 | 0.626942 | 1.69E-11 |
| ENSG00000143772 | ITPKB | 0.451944 | 0.683376 | 5.58E-12 |
| ENSG00000140497 | SCAMP2 | 0.968932 | 0.760546 | 3.29E-05 |
| ENSG00000161395 | PGAP3 | 0.430405 | 0.635043 | 6.83E-05 |
| ENSG00000149573 | MPZL2 | 0.284143 | 0.579596 | 6.03E-05 |
| ENSG00000134001 | EIF2S1 | 0.905262 | 0.692556 | 1.98E-11 |
| ENSG00000155307 | SAMSN1 | 0.903916 | 0.683707 | 0.000211 |
| ENSG00000188676 | IDO2 | 0.359828 | 0.647537 | 0.000255 |
| ENSG00000187474 | FPR3 | 0.838782 | 0.486005 | 0.000445 |
| ENSG00000008394 | MGST1 | 0.80655 | 0.51625 | 4.51E-13 |
| ENSG00000206190 | ATP10A | 0.437445 | 0.685725 | 2.50E-12 |
| ENSG00000077157 | PPP1R12B | 0.406153 | 0.69797 | 7.41E-21 |
| ENSG00000048052 | HDAC9 | 0.533311 | 0.781411 | 2.17E-14 |
| ENSG00000181019 | NQO1 | 0.773775 | 0.524212 | 1.31E-23 |
| ENSG00000133739 | LRRCC1 | 0.342101 | 0.691085 | 7.17E-13 |
| ENSG00000144744 | UBA3 | 0.91746 | 0.714251 | 5.32E-20 |
| ENSG00000171812 | COL8A2 | 0.410486 | 0.67669 | 3.41E-20 |
| ENSG00000120833 | SOCS2 | 0.702456 | 0.478634 | 8.76E-08 |
| ENSG00000185950 | IRS2 | 0.448338 | 0.718346 | 7.50E-36 |
| ENSG00000116266 | STXBP3 | 0.877527 | 0.64851 | 0.031716 |
| ENSG00000134460 | IL2RA | 0.62429 | 0.422763 | 0.000187 |
| ENSG00000100372 | SLC25A17 | 0.752739 | 0.544209 | 2.83E-11 |
| ENSG00000115828 | QPCT | 0.87689 | 0.606917 | 9.06E-05 |
| ENSG00000159714 | ZDHHC1 | 0.300121 | 0.549015 | 2.78E-08 |
| ENSG00000184205 | TSPYL2 | 0.386715 | 0.822086 | 3.51E-27 |
| ENSG00000243147 | MRPL33 | 0.897973 | 0.679149 | 3.68E-21 |
| ENSG00000047346 | FAM214A | 0.524682 | 0.748902 | 8.20E-18 |
| ENSG00000117479 | SLC19A2 | 0.416978 | 0.649384 | 6.37E-19 |
| ENSG00000166979 | EVA1C | 0.303457 | 0.538372 | 3.32E-06 |
| ENSG00000102445 | RUBCNL | 0.518773 | 0.763917 | 0.000697 |
| ENSG00000177575 | CD163 | 0.314231 | 0.618942 | 0.00036 |
| ENSG00000115594 | IL1R1 | 0.907946 | 0.557803 | 0.001363 |
| ENSG00000134909 | ARHGAP32 | 0.402012 | 0.709702 | 2.46E-33 |
| ENSG00000168685 | IL7R | 0.863073 | 0.584554 | 0.000144 |
| ENSG00000137642 | SORL1 | 0.42991 | 0.715331 | 2.38E-11 |
| ENSG00000162222 | TTC9C | 0.789281 | 0.572868 | 1.12E-50 |
| ENSG00000102898 | NUTF2 | 0.933743 | 0.694996 | 0.001295 |
| ENSG00000137267 | TUBB2A | 0.831901 | 0.618962 | 4.89E-08 |
| ENSG00000198286 | CARD11 | 0.354538 | 0.79206 | 1.89E-17 |
| ENSG00000136828 | RALGPS1 | 0.348757 | 0.583513 | 0.005475 |
| ENSG00000131931 | THAP1 | 0.781212 | 0.560297 | 3.39E-10 |
| ENSG00000009844 | VTA1 | 0.830893 | 0.608521 | 3.14E-05 |
| ENSG00000132423 | COQ3 | 0.270399 | 0.495398 | 1.89E-05 |
| ENSG00000152217 | SETBP1 | 0.415722 | 0.702394 | 1.39E-25 |
| ENSG00000134575 | ACP2 | 0.803919 | 0.556306 | 3.01E-07 |
| ENSG00000149489 | ROM1 | 0.345533 | 0.573934 | 0.00017 |
| ENSG00000115607 | IL18RAP | 0.161025 | 0.626695 | 0.00431 |
| ENSG00000213903 | LTB4R | 0.414051 | 0.719036 | 1.29E-13 |
| ENSG00000100099 | HPS4 | 0.514921 | 0.731752 | 1.90E-05 |
| ENSG00000155868 | MED7 | 0.7951 | 0.592225 | 1.43E-42 |
| ENSG00000188266 | HYKK | 0.250416 | 0.504211 | 0.007524 |
| ENSG00000254838 | GVINP1 | 0.231698 | 0.509358 | 6.91E-07 |
| ENSG00000105246 | EBI3 | 0.764308 | 0.42011 | 0.000139 |
| ENSG00000135541 | AHI1 | 0.456194 | 0.664197 | 1.49E-05 |
| ENSG00000071205 | ARHGAP10 | 0.832008 | 0.541793 | 1.34E-09 |
| ENSG00000025770 | NCAPH2 | 0.482916 | 0.710117 | 0.000159 |
| ENSG00000174842 | GLMN | 0.353049 | 0.566454 | 0.000319 |
| ENSG00000122223 | CD244 | 0.312157 | 0.624528 | 1.78E-11 |
| ENSG00000176225 | RTTN | 0.358839 | 0.641406 | 0.003707 |
| ENSG00000100784 | RPS6KA5 | 0.313075 | 0.612676 | 2.47E-14 |
| ENSG00000104951 | IL4I1 | 0.871744 | 0.647184 | 0.000144 |
| ENSG00000136897 | MRPL50 | 0.798799 | 0.539368 | 3.94E-08 |
| ENSG00000163823 | CCR1 | 0.769264 | 0.545175 | 0.000262 |
| ENSG00000150779 | TIMM8B | 0.838572 | 0.624512 | 1.35E-30 |
| ENSG00000158411 | MITD1 | 0.729279 | 0.521829 | 3.42E-17 |
| ENSG00000124813 | RUNX2 | 0.56015 | 0.765404 | 2.06E-08 |
| ENSG00000198270 | TMEM116 | 0.322129 | 0.559328 | 0.000302 |
| ENSG00000087884 | AAMDC | 0.763914 | 0.550238 | 2.95E-27 |
| ENSG00000111802 | TDP2 | 0.832129 | 0.618312 | 4.94E-11 |
| ENSG00000171202 | TMEM126A | 0.800787 | 0.589629 | 1.67E-12 |
| ENSG00000173369 | C1QB | 0.894117 | 0.483276 | 1.26E-05 |
| ENSG00000116791 | CRYZ | 0.771759 | 0.537978 | 3.56E-16 |
| ENSG00000123700 | KCNJ2 | 0.6415 | 0.436337 | 0.027584 |
| ENSG00000143499 | SMYD2 | 0.430769 | 0.633521 | 0.000235 |
| ENSG00000169490 | TM2D2 | 0.811393 | 0.560065 | 5.64E-16 |
| ENSG00000115514 | TXNDC9 | 0.773389 | 0.482342 | 1.29E-17 |
| ENSG00000120437 | ACAT2 | 0.882265 | 0.667731 | 1.78E-21 |
| ENSG00000109919 | MTCH2 | 0.948089 | 0.704781 | 4.40E-15 |
| ENSG00000103540 | CCP110 | 0.352612 | 0.572849 | 0.04108 |
| ENSG00000276045 | ORAI1 | 0.890876 | 0.650708 | 3.81E-08 |
| ENSG00000174123 | TLR10 | 0.279598 | 0.649506 | 1.24E-07 |
| ENSG00000085465 | OVGP1 | 0.240573 | 0.615524 | 5.52E-11 |
| ENSG00000158488 | CD1E | 0.902253 | 0.582504 | 1.32E-06 |
| ENSG00000113845 | TIMMDC1 | 0.895337 | 0.584573 | 1.07E-05 |
| ENSG00000168303 | MPLKIP | 0.815517 | 0.603059 | 1.86E-19 |
| ENSG00000117480 | FAAH | 0.375508 | 0.651652 | 2.87E-07 |
| ENSG00000165898 | ISCA2 | 0.780861 | 0.446685 | 9.24E-07 |
| ENSG00000163485 | ADORA1 | 0.370093 | 0.570643 | 0.000105 |
| ENSG00000134809 | TIMM10 | 0.832688 | 0.536327 | 4.07E-13 |
| ENSG00000159618 | ADGRG5 | 0.428724 | 0.63452 | 0.00203 |
| ENSG00000178175 | ZNF366 | 0.907759 | 0.672897 | 0.000464 |
| ENSG00000134755 | DSC2 | 0.795073 | 0.548015 | 0.000585 |
| ENSG00000008283 | CYB561 | 0.353219 | 0.605221 | 6.80E-08 |
| ENSG00000157445 | CACNA2D3 | 0.235609 | 0.53007 | 0.000217 |
| ENSG00000142405 | NLRP12 | 0.202876 | 0.426752 | 0.005683 |
| ENSG00000118200 | CAMSAP2 | 0.783107 | 0.492009 | 1.02E-12 |
| ENSG00000121552 | CSTA | 0.85061 | 0.469937 | 5.87E-05 |
| ENSG00000197077 | KIAA1671 | 0.848628 | 0.443171 | 8.19E-19 |
| ENSG00000146859 | TMEM140 | 0.746174 | 0.524219 | 0.011673 |
| ENSG00000099246 | RAB18 | 0.882238 | 0.680929 | 0.00057 |
| ENSG00000164743 | C8orf48 | 0.106537 | 0.417035 | 9.48E-07 |
| ENSG00000137288 | UQCC2 | 0.835065 | 0.620865 | 7.67E-05 |
| ENSG00000153208 | MERTK | 0.321597 | 0.544075 | 0.00056 |
| ENSG00000169075 | Z99496.1 | 0.141119 | 0.482826 | 0.001587 |
| ENSG00000196220 | SRGAP3 | 0.38992 | 0.618503 | 3.68E-05 |
| ENSG00000168653 | NDUFS5 | 0.956584 | 0.676828 | 9.92E-10 |
| ENSG00000057252 | SOAT1 | 0.908463 | 0.655731 | 0.001752 |
| ENSG00000255587 | RAB44 | 0.256838 | 0.494336 | 8.07E-07 |
| ENSG00000169439 | SDC2 | 0.913663 | 0.56966 | 7.53E-15 |
| ENSG00000167193 | CRK | 0.873363 | 0.631768 | 4.79E-08 |
| ENSG00000135698 | MPHOSPH6 | 0.824829 | 0.571235 | 7.94E-27 |
| ENSG00000111696 | NT5DC3 | 0.330004 | 0.657721 | 1.31E-06 |
| ENSG00000118257 | NRP2 | 0.848896 | 0.403606 | 5.04E-13 |
| ENSG00000076555 | ACACB | 0.264958 | 0.471767 | 0.032586 |
| ENSG00000128833 | MYO5C | 0.208339 | 0.533391 | 0.000393 |
| ENSG00000074706 | IPCEF1 | 0.821654 | 0.575686 | 3.09E-11 |
| ENSG00000163154 | TNFAIP8L2 | 0.767823 | 0.463854 | 6.67E-08 |
| ENSG00000154319 | FAM167A | 0.28753 | 0.564004 | 2.53E-09 |
| ENSG00000173404 | INSM1 | 0.662947 | 0.43409 | 0.000317 |
| ENSG00000109390 | NDUFC1 | 0.900756 | 0.556885 | 8.86E-28 |
| ENSG00000129521 | EGLN3 | 0.858703 | 0.541605 | 2.49E-08 |
| ENSG00000160446 | ZDHHC12 | 0.855486 | 0.629765 | 1.37E-08 |
| ENSG00000111875 | ASF1A | 0.803726 | 0.576809 | 0.00197 |
| ENSG00000123353 | ORMDL2 | 0.931375 | 0.580947 | 6.86E-15 |
| ENSG00000137502 | RAB30 | 0.891273 | 0.678933 | 0.000356 |
| ENSG00000096093 | EFHC1 | 0.274602 | 0.609441 | 0.00077 |
| ENSG00000168310 | IRF2 | 0.938144 | 0.667871 | 7.70E-10 |
| ENSG00000189221 | MAOA | 0.699147 | 0.275377 | 1.17E-10 |
| ENSG00000270804 | AC010326.4 | 0.761549 | 0.560777 | 1.22E-13 |
| ENSG00000206560 | ANKRD28 | 0.417086 | 0.659578 | 1.61E-12 |
| ENSG00000110660 | SLC35F2 | 0.255915 | 0.613497 | 3.57E-12 |
| ENSG00000181924 | COA4 | 0.839612 | 0.618025 | 3.63E-33 |
| ENSG00000163646 | CLRN1 | 0.081652 | 0.432369 | 0.006613 |
| ENSG00000171174 | RBKS | 0.264135 | 0.562863 | 6.71E-06 |
| ENSG00000066294 | CD84 | 0.932209 | 0.701755 | 7.55E-07 |
| ENSG00000156313 | RPGR | 0.292404 | 0.499759 | 0.009048 |
| ENSG00000165959 | CLMN | 0.369219 | 0.599417 | 0.008282 |
| ENSG00000262406 | MMP12 | 0.847886 | 0.29086 | 7.39E-08 |
| ENSG00000163520 | FBLN2 | 0.421923 | 0.716461 | 2.81E-12 |
| ENSG00000143155 | TIPRL | 0.881309 | 0.644015 | 2.38E-28 |
| ENSG00000169385 | RNASE2 | 0.685441 | 0.46557 | 7.95E-06 |
| ENSG00000111860 | CEP85L | 0.346128 | 0.603494 | 7.86E-10 |
| ENSG00000150768 | DLAT | 0.821891 | 0.591107 | 9.00E-09 |
| ENSG00000158485 | CD1B | 0.874241 | 0.442017 | 5.49E-15 |
| ENSG00000103544 | VPS35L | 0.86157 | 0.609775 | 0.011634 |
| ENSG00000052126 | PLEKHA5 | 0.834202 | 0.610534 | 4.25E-05 |
| ENSG00000123560 | PLP1 | 0.16172 | 0.431235 | 0.011537 |
| ENSG00000064763 | FAR2 | 0.360463 | 0.563111 | 0.000608 |
| ENSG00000007350 | TKTL1 | 0.248856 | 0.505236 | 1.34E-05 |
| ENSG00000162373 | BEND5 | 0.300218 | 0.601715 | 2.49E-05 |
| ENSG00000166068 | SPRED1 | 0.762635 | 0.467636 | 0.001169 |
| ENSG00000196975 | ANXA4 | 0.942458 | 0.565675 | 0.000576 |
| ENSG00000166670 | MMP10 | 0.545454 | 0.306953 | 9.82E-07 |
| ENSG00000137501 | SYTL2 | 0.208423 | 0.496287 | 0.00126 |
| ENSG00000152219 | ARL14EP | 0.88138 | 0.564305 | 4.02E-06 |
| ENSG00000182557 | SPNS3 | 0.393663 | 0.720725 | 1.30E-12 |
| ENSG00000107742 | SPOCK2 | 0.385473 | 0.679823 | 1.73E-10 |
| ENSG00000090263 | MRPS33 | 0.791473 | 0.577194 | 1.36E-21 |
| ENSG00000147155 | EBP | 0.862141 | 0.657188 | 3.45E-10 |
| ENSG00000099204 | ABLIM1 | 0.33358 | 0.535477 | 0.003759 |
| ENSG00000166477 | LEO1 | 0.855788 | 0.544809 | 1.67E-16 |
| ENSG00000137285 | TUBB2B | 0.719587 | 0.311235 | 2.89E-05 |
| ENSG00000142552 | RCN3 | 0.405469 | 0.646625 | 1.68E-07 |
| ENSG00000128534 | LSM8 | 0.876073 | 0.553589 | 6.91E-10 |
| ENSG00000164294 | GPX8 | 0.179874 | 0.400344 | 0.001067 |
| ENSG00000123268 | ATF1 | 0.839722 | 0.600474 | 0.041724 |
| ENSG00000118785 | SPP1 | 0.835136 | 0.452465 | 2.62E-16 |
| ENSG00000135094 | SDS | 0.351516 | 0.632426 | 6.06E-12 |
| ENSG00000106853 | PTGR1 | 0.651073 | 0.390954 | 9.71E-18 |
| ENSG00000151012 | SLC7A11 | 0.861214 | 0.465443 | 1.98E-06 |
| ENSG00000119599 | DCAF4 | 0.365829 | 0.567447 | 4.91E-06 |
| ENSG00000205755 | CRLF2 | 0.713178 | 0.439565 | 0.00395 |
| ENSG00000181896 | ZNF101 | 0.88821 | 0.522439 | 0.019822 |
| ENSG00000198937 | CCDC167 | 0.779247 | 0.55907 | 2.30E-15 |
| ENSG00000161835 | GRASP | 0.576918 | 0.796573 | 2.55E-14 |
| ENSG00000162924 | REL | 0.938438 | 0.729458 | 0.037631 |
| ENSG00000203782 | LOR | 0.5331 | 0.332153 | 8.01E-09 |
| ENSG00000174405 | LIG4 | 0.807539 | 0.592565 | 1.39E-26 |
| ENSG00000156299 | TIAM1 | 0.852108 | 0.594939 | 0.002009 |
| ENSG00000164164 | OTUD4 | 0.351383 | 0.723274 | 1.76E-31 |
| ENSG00000186265 | BTLA | 0.27756 | 0.616411 | 7.19E-08 |
| ENSG00000130649 | CYP2E1 | 0.348329 | 0.584719 | 6.33E-05 |
| ENSG00000198954 | KIF1BP | 0.766985 | 0.530359 | 1.38E-27 |
| ENSG00000187091 | PLCD1 | 0.39053 | 0.654242 | 1.20E-13 |
| ENSG00000256223 | ZNF10 | 0.264309 | 0.56706 | 5.04E-05 |
| ENSG00000155016 | CYP2U1 | 0.330045 | 0.579969 | 7.87E-12 |
| ENSG00000106565 | TMEM176B | 0.83098 | 0.380667 | 4.44E-08 |
| ENSG00000153130 | SCOC | 0.685553 | 0.456348 | 0.000104 |
| ENSG00000124145 | SDC4 | 0.734572 | 0.484936 | 0.00241 |
| ENSG00000123572 | NRK | 0.071398 | 0.409209 | 0.010219 |
| ENSG00000163739 | CXCL1 | 0.713589 | 0.385353 | 0.000702 |
| ENSG00000185338 | SOCS1 | 0.743671 | 0.51151 | 1.06E-07 |
| ENSG00000196218 | RYR1 | 0.354127 | 0.585763 | 4.72E-16 |
| ENSG00000169255 | B3GALNT1 | 0.610948 | 0.410443 | 1.71E-14 |
| ENSG00000126016 | AMOT | 0.279993 | 0.521888 | 1.57E-07 |
| ENSG00000165899 | OTOGL | 0.063106 | 0.400917 | 0.002619 |
| ENSG00000138382 | METTL5 | 0.858461 | 0.565575 | 0.003581 |
| ENSG00000178789 | CD300LB | 0.327126 | 0.63876 | 2.52E-12 |
| ENSG00000103196 | CRISPLD2 | 0.178314 | 0.400603 | 0.020264 |
| ENSG00000116133 | DHCR24 | 0.852821 | 0.579349 | 0.000294 |
| ENSG00000074416 | MGLL | 0.817057 | 0.614125 | 0.028561 |
| ENSG00000163735 | CXCL5 | 0.51566 | 0.288664 | 0.001438 |
| ENSG00000136522 | MRPL47 | 0.837383 | 0.580257 | 3.05E-05 |
| ENSG00000118518 | RNF146 | 0.765298 | 0.559264 | 4.21E-13 |
| ENSG00000198692 | EIF1AY | 0.83372 | 0.45832 | 1.13E-15 |
| ENSG00000105829 | BET1 | 0.707371 | 0.4213 | 4.53E-13 |
| ENSG00000114923 | SLC4A3 | 0.489127 | 0.745579 | 4.49E-10 |
| ENSG00000133104 | SPART | 0.859028 | 0.586983 | 2.27E-09 |
| ENSG00000196083 | IL1RAP | 0.75681 | 0.538425 | 7.26E-05 |
| ENSG00000188015 | S100A3 | 0.451255 | 0.237815 | 1.45E-05 |
| ENSG00000188505 | NCCRP1 | 0.760797 | 0.393346 | 6.17E-05 |
| ENSG00000105755 | ETHE1 | 0.836924 | 0.600847 | 1.27E-07 |
| ENSG00000175334 | BANF1 | 0.851774 | 0.626835 | 0.000942 |
| ENSG00000184465 | WDR27 | 0.312798 | 0.581927 | 1.00E-07 |
| ENSG00000116852 | KIF21B | 0.406762 | 0.623039 | 4.06E-15 |
| ENSG00000154134 | ROBO3 | 0.345547 | 0.620352 | 1.84E-05 |
| ENSG00000179115 | FARSA | 0.892352 | 0.635516 | 2.14E-09 |
| ENSG00000111711 | GOLT1B | 0.825996 | 0.543673 | 0.000154 |
| ENSG00000112304 | ACOT13 | 0.923127 | 0.577336 | 2.16E-06 |
| ENSG00000160961 | ZNF333 | 0.371043 | 0.611201 | 1.07E-06 |
| ENSG00000198829 | SUCNR1 | 0.747508 | 0.401777 | 8.63E-22 |
| ENSG00000159228 | CBR1 | 0.791122 | 0.47297 | 9.29E-16 |
| ENSG00000136244 | IL6 | 0.598846 | 0.372563 | 0.001823 |
| ENSG00000113263 | ITK | 0.230388 | 0.442229 | 0.000544 |
| ENSG00000169855 | ROBO1 | 0.78422 | 0.53309 | 0.013621 |
| ENSG00000229205 | LINC00200 | 0.181073 | 0.440269 | 4.37E-05 |
| ENSG00000143479 | DYRK3 | 0.221563 | 0.475681 | 1.36E-11 |
| ENSG00000157404 | KIT | 0.389678 | 0.637658 | 5.45E-07 |
| ENSG00000198730 | CTR9 | 0.834715 | 0.62743 | 0.014908 |
| ENSG00000136960 | ENPP2 | 0.856061 | 0.582559 | 0.005166 |
| ENSG00000207863 | MIR125B2 | 0.066949 | 0.407987 | 0.015387 |
| ENSG00000086289 | EPDR1 | 0.629968 | 0.413065 | 1.55E-11 |
| ENSG00000125869 | LAMP5 | 0.195588 | 0.441238 | 0.014033 |
| ENSG00000010327 | STAB1 | 0.735429 | 0.532105 | 0.038914 |
| ENSG00000154188 | ANGPT1 | 0.116871 | 0.450751 | 0.000588 |
| ENSG00000022267 | FHL1 | 0.292979 | 0.562716 | 1.68E-05 |
| ENSG00000186008 | RPS4XP21 | 0.091462 | 0.423212 | 4.86E-09 |
| ENSG00000067048 | DDX3Y | 0.774491 | 0.558768 | 1.90E-10 |
| ENSG00000119714 | GPR68 | 0.670835 | 0.299116 | 1.71E-12 |
| ENSG00000174776 | WDR49 | 0.219787 | 0.526734 | 3.28E-25 |
| ENSG00000118242 | MREG | 0.777115 | 0.499368 | 1.22E-25 |
| ENSG00000130038 | CRACR2A | 0.36179 | 0.597031 | 7.73E-20 |
| ENSG00000124491 | F13A1 | 0.75456 | 0.393526 | 7.26E-05 |
| ENSG00000134375 | TIMM17A | 0.924957 | 0.619832 | 8.58E-22 |
| ENSG00000122507 | BBS9 | 0.320599 | 0.546292 | 1.58E-05 |
| ENSG00000168528 | SERINC2 | 0.756287 | 0.422351 | 1.72E-06 |
| ENSG00000063127 | SLC6A16 | 0.243978 | 0.479231 | 5.39E-05 |
| ENSG00000112414 | ADGRG6 | 0.347307 | 0.579754 | 0.001864 |
| ENSG00000178573 | MAF | 0.720104 | 0.449933 | 0.000226 |
| ENSG00000198133 | TMEM229B | 0.338124 | 0.561574 | 0.000195 |
| ENSG00000141655 | TNFRSF11A | 0.816475 | 0.506025 | 1.37E-12 |
| ENSG00000140939 | NOL3 | 0.649968 | 0.441269 | 0.013815 |
| ENSG00000107789 | MINPP1 | 0.752167 | 0.465087 | 6.69E-13 |
| ENSG00000023330 | ALAS1 | 0.911453 | 0.611225 | 8.54E-12 |
| ENSG00000171643 | S100Z | 0.212841 | 0.428107 | 1.98E-09 |
| ENSG00000173065 | FAM222B | 0.823894 | 0.609299 | 2.53E-05 |
| ENSG00000164023 | SGMS2 | 0.325355 | 0.538068 | 1.79E-07 |
| ENSG00000198932 | GPRASP1 | 0.149653 | 0.484539 | 0.00158 |
| ENSG00000186001 | LRCH3 | 0.869135 | 0.613941 | 0.000406 |
| ENSG00000137124 | ALDH1B1 | 0.704985 | 0.454916 | 2.39E-08 |
| ENSG00000115306 | SPTBN1 | 0.391543 | 0.683684 | 7.61E-35 |
| ENSG00000183960 | KCNH8 | 0.2035 | 0.484418 | 0.017359 |
| ENSG00000142102 | PGGHG | 0.428481 | 0.635946 | 2.40E-09 |
| ENSG00000108352 | RAPGEFL1 | 0.282606 | 0.528003 | 0.000609 |
| ENSG00000165996 | HACD1 | 0.648896 | 0.414542 | 4.53E-22 |
| ENSG00000176771 | NCKAP5 | 0.350313 | 0.573076 | 3.14E-07 |
| ENSG00000108671 | PSMD11 | 0.907623 | 0.694795 | 4.55E-06 |
| ENSG00000104412 | EMC2 | 0.783679 | 0.527605 | 6.45E-09 |
| ENSG00000135045 | C9orf40 | 0.661394 | 0.401074 | 0.009663 |
| ENSG00000276644 | DACH1 | 0.240613 | 0.474345 | 0.001187 |
| ENSG00000117834 | SLC5A9 | 0.225439 | 0.452939 | 3.47E-07 |
| ENSG00000134962 | KLB | 0.234126 | 0.506057 | 2.19E-09 |
| ENSG00000165507 | DEPP1 | 0.775867 | 0.454919 | 2.83E-10 |
| ENSG00000155115 | GTF3C6 | 0.826431 | 0.572128 | 2.52E-11 |
| ENSG00000163840 | DTX3L | 0.862918 | 0.57893 | 0.001526 |
| ENSG00000106034 | CPED1 | 0.245559 | 0.575655 | 1.44E-17 |
| ENSG00000120217 | CD274 | 0.891531 | 0.300273 | 1.76E-14 |
| ENSG00000108423 | TUBD1 | 0.588554 | 0.384246 | 4.42E-11 |
| ENSG00000163618 | CADPS | 0.083806 | 0.398144 | 6.55E-06 |
| ENSG00000129538 | RNASE1 | 0.846433 | 0.396762 | 2.61E-16 |
| ENSG00000163596 | ICA1L | 0.228569 | 0.5683 | 1.53E-13 |
| ENSG00000106991 | ENG | 0.867856 | 0.589986 | 0.038984 |
| ENSG00000102057 | KCND1 | 0.307487 | 0.556843 | 8.31E-07 |
| ENSG00000159189 | C1QC | 0.911628 | 0.431875 | 5.63E-09 |
| ENSG00000158270 | COLEC12 | 0.703618 | 0.242751 | 7.76E-12 |
| ENSG00000120159 | CAAP1 | 0.689233 | 0.431021 | 0.002572 |
| ENSG00000167380 | ZNF226 | 0.681233 | 0.477221 | 1.19E-27 |
| ENSG00000079101 | CLUL1 | 0.154887 | 0.390246 | 1.61E-06 |
| ENSG00000151789 | ZNF385D | 0.105138 | 0.388479 | 0.019656 |
| ENSG00000165457 | FOLR2 | 0.558773 | 0.353935 | 0.020295 |
| ENSG00000165525 | NEMF | 0.897576 | 0.688017 | 0.009403 |
| ENSG00000242485 | MRPL20 | 0.926943 | 0.636312 | 1.12E-07 |
| ENSG00000151150 | ANK3 | 0.237723 | 0.522741 | 1.03E-12 |
| ENSG00000128596 | CCDC136 | 0.227177 | 0.448961 | 0.048715 |
| ENSG00000256683 | ZNF350 | 0.694655 | 0.494366 | 0.016336 |
| ENSG00000088543 | C3orf18 | 0.685144 | 0.421808 | 2.13E-11 |
| ENSG00000255302 | EID1 | 0.956644 | 0.658355 | 0.036279 |
| ENSG00000065833 | ME1 | 0.663027 | 0.331933 | 2.97E-11 |
| ENSG00000168273 | SMIM4 | 0.816118 | 0.510829 | 1.75E-10 |
| ENSG00000152582 | SPEF2 | 0.231901 | 0.451616 | 0.000656 |
| ENSG00000112130 | RNF8 | 0.72107 | 0.510593 | 1.07E-18 |
| ENSG00000168026 | TTC21A | 0.296306 | 0.525811 | 0.003344 |
| ENSG00000205060 | SLC35B4 | 0.716952 | 0.491537 | 0.000544 |
| ENSG00000163347 | CLDN1 | 0.838326 | 0.23538 | 1.03E-17 |
| ENSG00000005075 | POLR2J | 0.843743 | 0.544361 | 1.81E-11 |
| ENSG00000173585 | CCR9 | 0.132328 | 0.496651 | 2.99E-17 |
| ENSG00000100342 | APOL1 | 0.794834 | 0.5022 | 0.00021 |
| ENSG00000133134 | BEX2 | 0.276721 | 0.53235 | 0.000974 |
| ENSG00000131475 | VPS25 | 0.799087 | 0.576181 | 3.83E-08 |
| ENSG00000162437 | RAVER2 | 0.37931 | 0.674742 | 6.58E-20 |
| ENSG00000138772 | ANXA3 | 0.52907 | 0.279166 | 1.91E-11 |
| ENSG00000103855 | CD276 | 0.844229 | 0.394241 | 6.02E-21 |
| ENSG00000196440 | ARMCX4 | 0.171902 | 0.503464 | 1.38E-06 |
| ENSG00000173372 | C1QA | 0.737006 | 0.506602 | 3.78E-06 |
| ENSG00000203760 | CENPW | 0.660709 | 0.44288 | 3.72E-10 |
| ENSG00000160013 | PTGIR | 0.764788 | 0.412754 | 1.38E-09 |
| ENSG00000183454 | GRIN2A | 0.117727 | 0.397452 | 0.001022 |
| ENSG00000164107 | HAND2 | 0.179042 | 0.38392 | 9.92E-08 |
| ENSG00000116096 | SPR | 0.638582 | 0.267904 | 4.10E-09 |
| ENSG00000154262 | ABCA6 | 0.711726 | 0.375397 | 7.18E-21 |
| ENSG00000163029 | SMC6 | 0.706603 | 0.457619 | 0.002445 |
| ENSG00000163577 | EIF5A2 | 0.685269 | 0.446842 | 1.98E-16 |
| ENSG00000196184 | OR10J1 | 0.080724 | 0.366804 | 0.011364 |
| ENSG00000119965 | C10orf88 | 0.553964 | 0.341956 | 1.07E-06 |
| ENSG00000136051 | WASHC4 | 0.924922 | 0.549765 | 0.02256 |
| ENSG00000122176 | FMOD | 0.122523 | 0.386331 | 0.016996 |
| ENSG00000162891 | IL20 | 0.083908 | 0.383733 | 0.006272 |
| ENSG00000133069 | TMCC2 | 0.458958 | 0.216446 | 3.24E-06 |
| ENSG00000117595 | IRF6 | 0.203582 | 0.439653 | 0.005473 |
| ENSG00000227234 | SPANXB1 | 0.051357 | 0.368767 | 0.001995 |
| ENSG00000147655 | RSPO2 | 0.129645 | 0.371175 | 0.039439 |
| ENSG00000172243 | CLEC7A | 0.894275 | 0.519024 | 0.000152 |
| ENSG00000178718 | RPP25 | 0.655415 | 0.408625 | 2.00E-05 |
| ENSG00000139343 | SNRPF | 0.826559 | 0.579186 | 3.81E-09 |
| ENSG00000157450 | RNF111 | 0.840322 | 0.55865 | 0.000224 |
| ENSG00000147592 | LACTB2 | 0.56627 | 0.334154 | 7.35E-06 |
| ENSG00000237514 | PTP4A1P7 | 0.132527 | 0.41408 | 1.62E-16 |
| ENSG00000178665 | ZNF713 | 0.287254 | 0.51137 | 0.019543 |
| ENSG00000151917 | BEND6 | 0.191776 | 0.464156 | 8.12E-05 |
| ENSG00000180479 | ZNF571 | 0.283313 | 0.509291 | 0.000187 |
| ENSG00000177807 | KCNJ10 | 0.135516 | 0.413965 | 0.000113 |
| ENSG00000185518 | SV2B | 0.114877 | 0.376142 | 0.000299 |
| ENSG00000010278 | CD9 | 0.86905 | 0.438213 | 1.94E-11 |
| ENSG00000132329 | RAMP1 | 0.878422 | 0.409789 | 3.26E-08 |
| ENSG00000137270 | GCM1 | 0.11733 | 0.414817 | 0.000283 |
| ENSG00000167191 | GPRC5B | 0.255707 | 0.47679 | 9.37E-12 |
| ENSG00000102471 | NDFIP2 | 0.652441 | 0.414164 | 7.50E-17 |
| ENSG00000184005 | ST6GALNAC3 | 0.159032 | 0.379984 | 0.000416 |
| ENSG00000093134 | VNN3 | 0.120993 | 0.335947 | 0.027982 |
| ENSG00000114491 | UMPS | 0.740888 | 0.480326 | 1.64E-30 |
| ENSG00000143297 | FCRL5 | 0.196908 | 0.446564 | 0.019888 |
| ENSG00000182566 | CLEC4G | 0.657237 | 0.391479 | 2.18E-05 |
| ENSG00000122224 | LY9 | 0.413003 | 0.662329 | 1.40E-08 |
| ENSG00000112837 | TBX18 | 0.07221 | 0.364367 | 0.006365 |
| ENSG00000081087 | OSTM1 | 0.787193 | 0.463748 | 0.001 |
| ENSG00000152990 | ADGRA3 | 0.229246 | 0.533631 | 2.57E-06 |
| ENSG00000077616 | NAALAD2 | 0.125633 | 0.410453 | 0.007895 |
| ENSG00000108587 | GOSR1 | 0.818611 | 0.581688 | 8.80E-07 |
| ENSG00000233828 | LINC01949 | 0.145495 | 0.352162 | 4.06E-05 |
| ENSG00000100557 | CCDC198 | 0.04813 | 0.3414 | 0.000796 |
| ENSG00000137491 | SLCO2B1 | 0.740016 | 0.428036 | 0.000403 |
| ENSG00000120051 | CFAP58 | 0.116388 | 0.414623 | 0.003986 |
| ENSG00000256704 | SDCCAG3P1 | 0.121233 | 0.379706 | 0.001722 |
| ENSG00000117155 | SSX2IP | 0.284467 | 0.53532 | 1.59E-10 |
| ENSG00000173801 | JUP | 0.787658 | 0.552176 | 0.022958 |
| ENSG00000132170 | PPARG | 0.633713 | 0.334931 | 1.84E-10 |
| ENSG00000146263 | MMS22L | 0.307519 | 0.543947 | 0.003328 |
| ENSG00000135976 | ANKRD36 | 0.294444 | 0.515793 | 0.000405 |
| ENSG00000102243 | VGLL1 | 0.168253 | 0.373407 | 9.98E-06 |
| ENSG00000087842 | PIR | 0.587108 | 0.307569 | 7.74E-16 |
| ENSG00000177570 | SAMD12 | 0.178614 | 0.523908 | 9.78E-19 |
| ENSG00000154548 | SRSF12 | 0.53233 | 0.262237 | 3.68E-06 |
| ENSG00000196376 | SLC35F1 | 0.157775 | 0.363055 | 0.023413 |
| ENSG00000263002 | ZNF234 | 0.654195 | 0.41374 | 8.95E-20 |
| ENSG00000130032 | PRRG3 | 0.126994 | 0.366185 | 0.001129 |
| ENSG00000178795 | GDPD4 | 0.104288 | 0.342573 | 0.002033 |
| ENSG00000146938 | NLGN4X | 0.119168 | 0.338083 | 1.12E-06 |
| ENSG00000203688 | LINC02487 | 0.069625 | 0.357136 | 0.035636 |
| ENSG00000181218 | HIST3H2A | 0.256487 | 0.531493 | 4.26E-07 |
| ENSG00000196581 | AJAP1 | 0.208223 | 0.428805 | 0.028955 |
| ENSG00000002726 | AOC1 | 0.589152 | 0.271962 | 5.03E-12 |
| ENSG00000123178 | SPRYD7 | 0.745195 | 0.478057 | 4.76E-19 |
| ENSG00000145439 | CBR4 | 0.663706 | 0.398835 | 4.19E-08 |
| ENSG00000137204 | SLC22A7 | 0.199147 | 0.41719 | 0.000447 |
| ENSG00000187257 | RSBN1L | 0.756089 | 0.515211 | 0.000767 |
| ENSG00000169379 | ARL13B | 0.696914 | 0.439106 | 2.24E-13 |
| ENSG00000185737 | NRG3 | 0.123245 | 0.33261 | 7.87E-05 |
| ENSG00000124203 | ZNF831 | 0.207414 | 0.416374 | 3.26E-08 |
| ENSG00000095970 | TREM2 | 0.661819 | 0.402702 | 2.33E-10 |
| ENSG00000144868 | TMEM108 | 0.120529 | 0.401497 | 0.017736 |
| ENSG00000197279 | ZNF165 | 0.245915 | 0.541873 | 3.57E-19 |
| ENSG00000182333 | LIPF | 0.043149 | 0.355463 | 0.006322 |
| ENSG00000137731 | FXYD2 | 0.686669 | 0.465939 | 3.23E-07 |
| ENSG00000144840 | RABL3 | 0.843679 | 0.559194 | 0.000682 |
| ENSG00000151322 | NPAS3 | 0.166307 | 0.38605 | 0.007326 |
| ENSG00000063438 | AHRR | 0.805577 | 0.334966 | 2.90E-13 |
| ENSG00000153347 | FAM81B | 0.162905 | 0.393638 | 0.000236 |
| ENSG00000116157 | GPX7 | 0.20933 | 0.485377 | 0.000501 |
| ENSG00000177337 | DLGAP1-AS1 | 0.601069 | 0.371211 | 1.06E-08 |
| ENSG00000112182 | BACH2 | 0.235175 | 0.536706 | 2.58E-16 |
| ENSG00000134057 | CCNB1 | 0.672802 | 0.426778 | 2.72E-17 |
| ENSG00000165113 | GKAP1 | 0.152622 | 0.437266 | 5.64E-05 |
| ENSG00000008256 | CYTH3 | 0.302443 | 0.59345 | 9.34E-23 |
| ENSG00000145388 | METTL14 | 0.746419 | 0.545906 | 0.005681 |
| ENSG00000185031 | SLC2A3P2 | 0.092343 | 0.378301 | 0.032171 |
| ENSG00000164082 | GRM2 | 0.217255 | 0.438871 | 2.74E-13 |
| ENSG00000069702 | TGFBR3 | 0.163794 | 0.413954 | 0.007618 |
| ENSG00000092068 | SLC7A8 | 0.730204 | 0.446866 | 8.77E-07 |
| ENSG00000161405 | IKZF3 | 0.214294 | 0.494646 | 7.17E-06 |
| ENSG00000170370 | EMX2 | 0.103567 | 0.365671 | 1.94E-10 |
| ENSG00000064309 | CDON | 0.189693 | 0.420614 | 0.000411 |
| ENSG00000160208 | RRP1B | 0.765019 | 0.546569 | 0.040282 |
| ENSG00000153064 | BANK1 | 0.315877 | 0.530535 | 0.000545 |
| ENSG00000111186 | WNT5B | 0.615521 | 0.378183 | 9.55E-14 |
| ENSG00000109787 | KLF3 | 0.862865 | 0.649444 | 1.31E-05 |
| ENSG00000164347 | GFM2 | 0.76567 | 0.554813 | 2.79E-14 |
| ENSG00000208892 | SNORA49 | 0.130413 | 0.44502 | 2.04E-17 |
| ENSG00000144847 | IGSF11 | 0.183811 | 0.397015 | 0.010294 |
| ENSG00000173421 | CCDC36 | 0.096175 | 0.395172 | 4.92E-06 |
| ENSG00000115289 | PCGF1 | 0.694574 | 0.488943 | 2.82E-07 |
| ENSG00000099282 | TSPAN15 | 0.783122 | 0.538019 | 2.98E-05 |
| ENSG00000134028 | ADAMDEC1 | 0.575292 | 0.279334 | 0.00145 |
| ENSG00000215018 | COL28A1 | 0.148741 | 0.363705 | 3.40E-05 |
| ENSG00000145721 | LIX1 | 0.116674 | 0.361719 | 0.037119 |
| ENSG00000064989 | CALCRL | 0.8915 | 0.554105 | 0.000245 |
| ENSG00000158683 | PKD1L1 | 0.203345 | 0.413904 | 0.012437 |
| ENSG00000197980 | LEKR1 | 0.147152 | 0.431197 | 0.010983 |
| ENSG00000196632 | WNK3 | 0.113188 | 0.381559 | 0.010739 |
| ENSG00000118418 | HMGN3 | 0.950582 | 0.597508 | 7.81E-23 |
| ENSG00000178093 | TSSK6 | 0.208002 | 0.437216 | 0.002526 |
| ENSG00000167014 | TERB2 | 0.035481 | 0.322448 | 0.04403 |
| ENSG00000185924 | RTN4RL1 | 0.177751 | 0.385578 | 0.000514 |
| ENSG00000138385 | SSB | 0.951198 | 0.611474 | 0.014297 |
| ENSG00000115425 | PECR | 0.547368 | 0.333693 | 0.002551 |
| ENSG00000104760 | FGL1 | 0.069371 | 0.327936 | 0.02614 |
| ENSG00000100439 | ABHD4 | 0.737705 | 0.505072 | 9.38E-06 |
| ENSG00000143369 | ECM1 | 0.69471 | 0.397079 | 5.19E-09 |
| ENSG00000166750 | SLFN5 | 0.808402 | 0.531529 | 0.022253 |
| ENSG00000011009 | LYPLA2 | 0.85036 | 0.596734 | 0.001808 |
| ENSG00000117507 | FMO6P | 0.077808 | 0.365388 | 1.73E-05 |
| ENSG00000140623 | 12-Sep | 0.143008 | 0.403958 | 1.57E-09 |
| ENSG00000198093 | ZNF649 | 0.696264 | 0.420446 | 2.66E-11 |
| ENSG00000172987 | HPSE2 | 0.127188 | 0.347608 | 0.005373 |
| ENSG00000196166 | C8orf86 | 0.108496 | 0.347967 | 0.000456 |
| ENSG00000166167 | BTRC | 0.689722 | 0.44609 | 1.24E-05 |
| ENSG00000163900 | TMEM41A | 0.855872 | 0.496994 | 0.003053 |
| ENSG00000105205 | CLC | 0.643399 | 0.41154 | 2.29E-05 |
| ENSG00000079112 | CDH17 | 0.122421 | 0.466304 | 2.24E-09 |
| ENSG00000172000 | ZNF556 | 0.106808 | 0.33525 | 7.65E-05 |
| ENSG00000187037 | GPR141 | 0.803281 | 0.451895 | 0.008353 |
| ENSG00000140848 | CPNE2 | 0.723108 | 0.521744 | 0.033285 |
| ENSG00000157483 | MYO1E | 0.858944 | 0.620793 | 3.18E-07 |
| ENSG00000126545 | CSN1S1 | 0.07824 | 0.324397 | 0.007618 |
| ENSG00000145817 | YIPF5 | 0.849352 | 0.527148 | 6.47E-10 |
| ENSG00000120075 | HOXB5 | 0.142604 | 0.370801 | 0.007705 |
| ENSG00000090402 | SI | 0.044247 | 0.34514 | 0.038062 |
| ENSG00000113578 | FGF1 | 0.093366 | 0.340415 | 9.49E-07 |
| ENSG00000176697 | BDNF | 0.137795 | 0.373412 | 0.026495 |
| ENSG00000163606 | CD200R1 | 0.720861 | 0.479889 | 0.028052 |
| ENSG00000243064 | ABCC13 | 0.133264 | 0.345045 | 0.03473 |
| ENSG00000165417 | GTF2A1 | 0.882412 | 0.586157 | 0.022707 |
| ENSG00000154678 | PDE1C | 0.145846 | 0.429684 | 8.25E-10 |
| ENSG00000168994 | PXDC1 | 0.749753 | 0.544142 | 0.018247 |
| ENSG00000155621 | C9orf85 | 0.706109 | 0.458824 | 1.04E-06 |
| ENSG00000131018 | SYNE1 | 0.335666 | 0.591191 | 1.68E-18 |
| ENSG00000214653 | HNRNPA3P3 | 0.122369 | 0.383102 | 0.044233 |
| ENSG00000176563 | CNTD1 | 0.321639 | 0.521824 | 0.000763 |
| ENSG00000086717 | PPEF1 | 0.082132 | 0.347703 | 0.000206 |
| ENSG00000172339 | ALG14 | 0.641358 | 0.432883 | 3.94E-13 |
| ENSG00000176343 | RPL37AP8 | 0.706691 | 0.467725 | 9.92E-06 |
| ENSG00000176246 | OR4L1 | 0.114385 | 0.333119 | 0.017783 |
| ENSG00000203664 | OR2W5 | 0.136548 | 0.34743 | 1.44E-07 |
| ENSG00000173809 | TDRD12 | 0.103206 | 0.363071 | 0.026492 |
| ENSG00000142156 | COL6A1 | 0.681006 | 0.416986 | 0.000139 |
| ENSG00000230992 | FAM201B | 0.60473 | 0.368777 | 0.047364 |
| ENSG00000100722 | ZC3H14 | 0.754731 | 0.542045 | 0.010516 |
| ENSG00000205981 | DNAJC19 | 0.739787 | 0.365867 | 2.08E-14 |
| ENSG00000112530 | PACRG | 0.118807 | 0.348245 | 0.001744 |
| ENSG00000124422 | USP22 | 0.939619 | 0.70542 | 0.002674 |
| ENSG00000162620 | LRRIQ3 | 0.084022 | 0.339844 | 0.029556 |
| ENSG00000145757 | SPATA9 | 0.15745 | 0.362308 | 1.76E-08 |
| ENSG00000118655 | DCLRE1B | 0.689009 | 0.404574 | 0.000218 |
| ENSG00000177133 | LINC00982 | 0.603058 | 0.370527 | 0.001422 |
| ENSG00000163624 | CDS1 | 0.220092 | 0.499052 | 2.28E-10 |
| ENSG00000102780 | DGKH | 0.670295 | 0.470104 | 1.84E-08 |
| ENSG00000177425 | PAWR | 0.272009 | 0.499802 | 4.92E-05 |
| ENSG00000036672 | USP2 | 0.21013 | 0.426038 | 2.26E-06 |
| ENSG00000140009 | ESR2 | 0.13909 | 0.37822 | 0.009173 |
| ENSG00000147231 | CXorf57 | 0.104013 | 0.365961 | 2.39E-07 |
| ENSG00000157765 | SLC34A2 | 0.130607 | 0.368418 | 5.85E-06 |
| ENSG00000176046 | NUPR1 | 0.582521 | 0.377965 | 6.55E-05 |
| ENSG00000186806 | VSIG10L | 0.643331 | 0.422131 | 1.75E-06 |
| ENSG00000152779 | SLC16A12 | 0.081687 | 0.337289 | 4.93E-05 |
| ENSG00000132465 | JCHAIN | 0.262555 | 0.475857 | 0.025105 |
| ENSG00000235072 | AC012074.1 | 0.148111 | 0.397824 | 0.018597 |
| ENSG00000180098 | TRNAU1AP | 0.794171 | 0.493472 | 1.25E-09 |
| ENSG00000179136 | LINC00670 | 0.079889 | 0.293539 | 0.045207 |
| ENSG00000144852 | NR1I2 | 0.158249 | 0.365189 | 0.001497 |
| ENSG00000115380 | EFEMP1 | 0.108793 | 0.323736 | 0.005497 |
| ENSG00000138175 | ARL3 | 0.806642 | 0.385748 | 2.67E-11 |
| ENSG00000087301 | TXNDC16 | 0.255267 | 0.506003 | 2.05E-15 |
| ENSG00000196417 | ZNF765 | 0.767283 | 0.479033 | 0.047462 |
| ENSG00000142173 | COL6A2 | 0.636935 | 0.417679 | 0.005169 |
| ENSG00000125744 | RTN2 | 0.648732 | 0.445443 | 3.36E-08 |
| ENSG00000154479 | CCDC173 | 0.085848 | 0.371457 | 0.000188 |
| ENSG00000135749 | PCNX2 | 0.335301 | 0.58084 | 2.19E-24 |
| ENSG00000114757 | PEX5L | 0.213604 | 0.421069 | 2.63E-12 |
| ENSG00000197748 | CFAP43 | 0.090408 | 0.35721 | 0.048595 |
| ENSG00000101596 | SMCHD1 | 0.893643 | 0.663344 | 0.010086 |
| ENSG00000204136 | GGTA1P | 0.856988 | 0.514301 | 1.44E-07 |
| ENSG00000083520 | DIS3 | 0.864323 | 0.57688 | 2.04E-06 |
| ENSG00000109686 | SH3D19 | 0.216499 | 0.476107 | 6.25E-09 |
| ENSG00000140006 | WDR89 | 0.760768 | 0.492122 | 1.53E-27 |
| ENSG00000165209 | STRBP | 0.68561 | 0.469514 | 0.009631 |
| ENSG00000256660 | CLEC12B | 0.116532 | 0.329583 | 7.43E-06 |
| ENSG00000103021 | CCDC113 | 0.146866 | 0.397068 | 0.005164 |
| ENSG00000173535 | TNFRSF10C | 0.199173 | 0.418634 | 0.02447 |
| ENSG00000176635 | HORMAD2 | 0.070308 | 0.302904 | 0.049747 |
| ENSG00000186714 | CCDC73 | 0.177544 | 0.466752 | 0.019798 |
| ENSG00000136425 | CIB2 | 0.266428 | 0.482613 | 1.27E-05 |
| ENSG00000171560 | FGA | 0.072468 | 0.326491 | 0.023571 |
| ENSG00000128342 | LIF | 0.222504 | 0.454763 | 1.85E-06 |
| ENSG00000158186 | MRAS | 0.741567 | 0.412705 | 1.66E-05 |
| ENSG00000004660 | CAMKK1 | 0.317595 | 0.535049 | 1.79E-15 |
| ENSG00000126267 | COX6B1 | 0.977267 | 0.63178 | 2.61E-06 |
| ENSG00000214300 | SPDYE3 | 0.167099 | 0.389513 | 0.025565 |
| ENSG00000145075 | CCDC39 | 0.144145 | 0.445828 | 0.007144 |
| ENSG00000155530 | LRGUK | 0.148891 | 0.399699 | 0.000225 |
| ENSG00000165626 | BEND7 | 0.112685 | 0.32892 | 0.037528 |
| ENSG00000260339 | HEXA-AS1 | 0.172595 | 0.401675 | 0.000256 |
| ENSG00000120925 | RNF170 | 0.753327 | 0.362398 | 2.53E-09 |
| ENSG00000178796 | RIIAD1 | 0.137129 | 0.353805 | 4.60E-05 |
| ENSG00000231160 | KLF3-AS1 | 0.202532 | 0.448261 | 1.33E-10 |
| ENSG00000145777 | TSLP | 0.095096 | 0.312662 | 1.05E-06 |
| ENSG00000148483 | TMEM236 | 0.620431 | 0.302465 | 2.94E-13 |
| ENSG00000100336 | APOL4 | 0.690052 | 0.395379 | 3.15E-11 |
| ENSG00000188176 | SMTNL2 | 0.55919 | 0.304709 | 0.002838 |
| ENSG00000144395 | CCDC150 | 0.186861 | 0.391607 | 0.022131 |
| ENSG00000180938 | ZNF572 | 0.139642 | 0.355692 | 0.002529 |
| ENSG00000138336 | TET1 | 0.128276 | 0.396464 | 0.000445 |
| ENSG00000148935 | GAS2 | 0.150517 | 0.366089 | 0.008592 |
| ENSG00000169418 | NPR1 | 0.67967 | 0.360239 | 9.73E-18 |
| ENSG00000177383 | MAGEF1 | 0.778083 | 0.507363 | 0.003308 |
| ENSG00000132855 | ANGPTL3 | 0.082271 | 0.371456 | 5.68E-08 |
| ENSG00000196876 | SCN8A | 0.181836 | 0.398293 | 8.95E-07 |
| ENSG00000125888 | BANF2 | 0.100461 | 0.307999 | 0.007605 |
| ENSG00000115290 | GRB14 | 0.056961 | 0.312131 | 0.023339 |
| ENSG00000170807 | LMOD2 | 0.191173 | 0.397921 | 3.68E-12 |
| ENSG00000182704 | TSKU | 0.641137 | 0.193539 | 4.54E-11 |
| ENSG00000184860 | SDR42E1 | 0.130802 | 0.365822 | 0.017764 |
| ENSG00000151773 | CCDC122 | 0.217702 | 0.43232 | 2.49E-05 |
| ENSG00000173597 | SULT1B1 | 0.120345 | 0.415646 | 2.93E-13 |
| ENSG00000133985 | TTC9 | 0.6507 | 0.349387 | 0.037108 |
| ENSG00000054654 | SYNE2 | 0.222613 | 0.494235 | 7.99E-05 |
| ENSG00000147669 | POLR2K | 0.911722 | 0.512212 | 5.06E-07 |
| ENSG00000120526 | NUDCD1 | 0.638216 | 0.398178 | 3.87E-06 |
| ENSG00000008517 | IL32 | 0.659218 | 0.349827 | 0.000321 |
| ENSG00000185046 | ANKS1B | 0.156929 | 0.447733 | 0.001436 |
| ENSG00000099840 | IZUMO4 | 0.644374 | 0.395864 | 0.003321 |
| ENSG00000075223 | SEMA3C | 0.719886 | 0.495679 | 3.42E-10 |
| ENSG00000276759 | AL353753.1 | 0.067973 | 0.304456 | 0.010483 |
| ENSG00000056736 | IL17RB | 0.691989 | 0.343225 | 2.28E-16 |
| ENSG00000198205 | ZXDA | 0.745192 | 0.471351 | 0.000981 |
| ENSG00000064652 | SNX24 | 0.567747 | 0.309922 | 1.58E-11 |
| ENSG00000163053 | SLC16A14 | 0.178834 | 0.389021 | 4.83E-06 |
| ENSG00000114670 | NEK11 | 0.155235 | 0.370717 | 0.020234 |
| ENSG00000138135 | CH25H | 0.619897 | 0.323068 | 0.037676 |
| ENSG00000116580 | GON4L | 0.773625 | 0.573486 | 0.006166 |
| ENSG00000198498 | TMA16 | 0.670203 | 0.451948 | 0.024031 |
| ENSG00000185414 | MRPL30 | 0.786609 | 0.454884 | 0.004293 |
| ENSG00000123342 | MMP19 | 0.688311 | 0.451935 | 7.88E-11 |
| ENSG00000139350 | NEDD1 | 0.720092 | 0.495549 | 0.000183 |
| ENSG00000105855 | ITGB8 | 0.559783 | 0.243407 | 7.24E-05 |
| ENSG00000082556 | OPRK1 | 0.087379 | 0.337282 | 0.014423 |
| ENSG00000171385 | KCND3 | 0.197428 | 0.474442 | 3.05E-15 |
| ENSG00000161911 | TREML1 | 0.67133 | 0.377917 | 2.66E-09 |
| ENSG00000143401 | ANP32E | 0.787421 | 0.559207 | 0.00064 |
| ENSG00000161992 | PRR35 | 0.549056 | 0.347394 | 0.034243 |
| ENSG00000101335 | MYL9 | 0.659602 | 0.391209 | 7.17E-05 |
| ENSG00000156218 | ADAMTSL3 | 0.109166 | 0.32323 | 0.048773 |
| ENSG00000221923 | ZNF880 | 0.589183 | 0.384598 | 1.75E-11 |
| ENSG00000244588 | RAD21L1 | 0.055333 | 0.359695 | 0.005003 |
| ENSG00000178429 | RPS3AP5 | 0.792911 | 0.410738 | 0.014605 |
| ENSG00000109047 | RCVRN | 0.214747 | 0.434478 | 3.08E-09 |
| ENSG00000130988 | RGN | 0.100213 | 0.329401 | 0.018252 |
| ENSG00000106683 | LIMK1 | 0.793345 | 0.579625 | 0.046014 |
| ENSG00000137168 | PPIL1 | 0.647759 | 0.384768 | 0.000478 |
| ENSG00000137726 | FXYD6 | 0.741153 | 0.308947 | 1.53E-06 |
| ENSG00000102970 | CCL17 | 0.889069 | 0.201742 | 2.26E-14 |
| ENSG00000133048 | CHI3L1 | 0.603164 | 0.359658 | 2.98E-11 |
| ENSG00000184698 | OR51M1 | 0.08693 | 0.304749 | 0.001162 |
| ENSG00000100867 | DHRS2 | 0.593446 | 0.281914 | 2.39E-14 |
| ENSG00000185900 | POMK | 0.683404 | 0.475826 | 2.94E-05 |
| ENSG00000154447 | SH3RF1 | 0.227082 | 0.447133 | 1.53E-20 |
| ENSG00000172057 | ORMDL3 | 0.704502 | 0.417495 | 1.68E-06 |
| ENSG00000183648 | NDUFB1 | 0.90678 | 0.497795 | 2.65E-22 |
| ENSG00000176927 | EFCAB5 | 0.120553 | 0.36748 | 0.028508 |
| ENSG00000162997 | PRORSD1P | 0.587688 | 0.257987 | 7.12E-05 |
| ENSG00000175105 | ZNF654 | 0.72865 | 0.517367 | 0.000169 |
| ENSG00000128040 | SPINK2 | 0.10673 | 0.316943 | 0.000891 |
| ENSG00000159884 | CCDC107 | 0.7797 | 0.513323 | 0.000298 |
| ENSG00000141753 | IGFBP4 | 0.785846 | 0.34921 | 1.40E-07 |
| ENSG00000115414 | FN1 | 0.653988 | 0.427383 | 1.69E-08 |
| ENSG00000146733 | PSPH | 0.598385 | 0.379562 | 1.23E-09 |
| ENSG00000120333 | MRPS14 | 0.721866 | 0.432321 | 4.45E-07 |
| ENSG00000170522 | ELOVL6 | 0.126448 | 0.334577 | 0.002431 |
| ENSG00000159199 | ATP5MC1 | 0.895464 | 0.520304 | 0.000347 |
| ENSG00000186891 | TNFRSF18 | 0.620956 | 0.366139 | 2.19E-05 |
| ENSG00000164659 | KIAA1324L | 0.183701 | 0.39392 | 0.001806 |
| ENSG00000198780 | FAM169A | 0.192615 | 0.432139 | 5.66E-09 |
| ENSG00000204007 | GLT6D1 | 0.082382 | 0.291692 | 0.001005 |
| ENSG00000011405 | PIK3C2A | 0.772204 | 0.565832 | 2.49E-05 |
| ENSG00000181374 | CCL13 | 0.826085 | 0.202291 | 1.87E-26 |
| ENSG00000148803 | FUOM | 0.794428 | 0.493115 | 0.003489 |
| ENSG00000198812 | LRRC10 | 0.073294 | 0.295771 | 6.34E-05 |
| ENSG00000102225 | CDK16 | 0.794793 | 0.576743 | 0.009069 |
| ENSG00000103061 | SLC7A6OS | 0.812081 | 0.602463 | 0.003523 |
| ENSG00000151366 | NDUFC2 | 0.939912 | 0.544433 | 9.61E-08 |
| ENSG00000147650 | LRP12 | 0.752806 | 0.335123 | 4.50E-17 |
| ENSG00000127083 | OMD | 0.102737 | 0.342986 | 0.000306 |
| ENSG00000108599 | AKAP10 | 0.804904 | 0.530298 | 2.80E-06 |
| ENSG00000100368 | CSF2RB | 0.962984 | 0.635846 | 0.001461 |
| ENSG00000120555 | SEPT7P9 | 0.67381 | 0.35051 | 0.014597 |
| ENSG00000167720 | SRR | 0.701014 | 0.436895 | 0.000269 |
| ENSG00000185972 | CCIN | 0.141507 | 0.376089 | 6.38E-18 |
| ENSG00000144824 | PHLDB2 | 0.149096 | 0.373909 | 5.00E-08 |
| ENSG00000127774 | EMC6 | 0.749747 | 0.509894 | 1.92E-11 |
| ENSG00000106244 | PDAP1 | 0.858643 | 0.588931 | 4.03E-06 |
| ENSG00000117594 | HSD11B1 | 0.429791 | 0.217906 | 5.14E-10 |
| ENSG00000088836 | SLC4A11 | 0.605393 | 0.349018 | 3.07E-10 |
| ENSG00000197302 | ZNF720 | 0.723506 | 0.434418 | 5.52E-09 |
| ENSG00000150637 | CD226 | 0.887885 | 0.492171 | 0.047258 |
| ENSG00000197653 | DNAH10 | 0.146868 | 0.366864 | 1.22E-05 |
| ENSG00000163435 | ELF3 | 0.192377 | 0.417337 | 2.91E-17 |
| ENSG00000185519 | FAM131C | 0.569748 | 0.317074 | 0.040788 |
| ENSG00000174903 | RAB1B | 0.902907 | 0.65337 | 1.37E-05 |
| ENSG00000178125 | PPP1R42 | 0.142643 | 0.384648 | 0.001151 |
| ENSG00000249774 | AC025458.1 | 0.656973 | 0.412174 | 0.001616 |
| ENSG00000103056 | SMPD3 | 0.294997 | 0.536544 | 2.85E-20 |
| ENSG00000163960 | UBXN7 | 0.772655 | 0.568303 | 0.010848 |
| ENSG00000141255 | SPATA22 | 0.067929 | 0.276403 | 0.001365 |
| ENSG00000146243 | IRAK1BP1 | 0.191757 | 0.416215 | 0.000209 |
| ENSG00000188778 | ADRB3 | 0.489348 | 0.284001 | 6.73E-05 |
| ENSG00000115457 | IGFBP2 | 0.561194 | 0.322393 | 1.06E-08 |
| ENSG00000071677 | PRLH | 0.511636 | 0.302939 | 0.00037 |
| ENSG00000177700 | POLR2L | 0.931069 | 0.492674 | 0.002611 |
| ENSG00000172728 | FUT10 | 0.630427 | 0.415534 | 8.65E-10 |
| ENSG00000108700 | CCL8 | 0.587502 | 0.209744 | 1.97E-11 |
| ENSG00000105429 | MEGF8 | 0.743225 | 0.507044 | 0.006904 |
| ENSG00000188725 | SMIM15 | 0.81875 | 0.478014 | 4.85E-23 |
| ENSG00000140961 | OSGIN1 | 0.60149 | 0.372513 | 0.000395 |
| ENSG00000115159 | GPD2 | 0.782824 | 0.469655 | 3.45E-08 |
| ENSG00000165066 | NKX6-3 | 0.554467 | 0.348515 | 0.002092 |
| ENSG00000104946 | TBC1D17 | 0.788319 | 0.549559 | 0.000211 |
| ENSG00000223803 | RPS20P14 | 0.778145 | 0.370314 | 0.016111 |
| ENSG00000074071 | MRPS34 | 0.763702 | 0.500776 | 0.01099 |
| ENSG00000168802 | CHTF8 | 0.897509 | 0.567138 | 0.000244 |
| ENSG00000121446 | RGSL1 | 0.073747 | 0.288652 | 0.004538 |
| ENSG00000172366 | MCRIP2 | 0.66439 | 0.458518 | 1.36E-05 |
| ENSG00000138660 | AP1AR | 0.71621 | 0.462799 | 2.62E-05 |
| ENSG00000151612 | ZNF827 | 0.648633 | 0.412688 | 0.017486 |
| ENSG00000001626 | CFTR | 0.092643 | 0.305115 | 0.03954 |
| ENSG00000155052 | CNTNAP5 | 0.072852 | 0.303826 | 0.022865 |
| ENSG00000164346 | NSA2 | 0.884211 | 0.562421 | 0.004314 |
| ENSG00000119673 | ACOT2 | 0.647604 | 0.419972 | 0.044136 |
| ENSG00000132694 | ARHGEF11 | 0.710154 | 0.420596 | 6.06E-05 |
| ENSG00000016490 | CLCA1 | 0.036703 | 0.289547 | 1.39E-09 |
| ENSG00000125434 | SLC25A35 | 0.691307 | 0.476201 | 1.08E-05 |
| ENSG00000042286 | AIFM2 | 0.563495 | 0.358105 | 1.20E-06 |
| ENSG00000127920 | GNG11 | 0.577349 | 0.303561 | 7.08E-15 |
| ENSG00000108702 | CCL1 | 0.627185 | 0.257677 | 1.56E-13 |
| ENSG00000131951 | LRRC9 | 0.059716 | 0.279852 | 0.024047 |
| ENSG00000186577 | SMIM29 | 0.724327 | 0.469909 | 7.28E-07 |
| ENSG00000067646 | ZFY | 0.602011 | 0.398432 | 1.19E-06 |
| ENSG00000121931 | LRIF1 | 0.782421 | 0.420591 | 6.20E-24 |
| ENSG00000148848 | ADAM12 | 0.574281 | 0.356087 | 9.85E-11 |
| ENSG00000150627 | WDR17 | 0.081581 | 0.361848 | 2.36E-06 |
| ENSG00000197520 | FAM177B | 0.110787 | 0.312813 | 0.001088 |
| ENSG00000163263 | C1orf189 | 0.095335 | 0.304704 | 0.001679 |
| ENSG00000172023 | REG1B | 0.043241 | 0.250434 | 0.021333 |
| ENSG00000164142 | FAM160A1 | 0.224077 | 0.512551 | 1.30E-08 |
| ENSG00000165895 | ARHGAP42 | 0.137723 | 0.371251 | 4.86E-05 |
| ENSG00000131831 | RAI2 | 0.498023 | 0.289581 | 9.88E-07 |
| ENSG00000204283 | LINC01973 | 0.095922 | 0.340651 | 1.47E-09 |
| ENSG00000200169 | RNU5D-1 | 0.044194 | 0.284317 | 0.00301 |
| ENSG00000204116 | CHIC1 | 0.199703 | 0.422968 | 2.36E-15 |
| ENSG00000044012 | GUCA2B | 0.52419 | 0.290659 | 0.000649 |
| ENSG00000075391 | RASAL2 | 0.803969 | 0.428852 | 5.60E-05 |
| ENSG00000152683 | SLC30A6 | 0.664524 | 0.447919 | 0.010705 |
| ENSG00000160191 | PDE9A | 0.504419 | 0.26676 | 0.004841 |
| ENSG00000108433 | GOSR2 | 0.783204 | 0.483836 | 0.00206 |
| ENSG00000162601 | MYSM1 | 0.812486 | 0.598843 | 0.008766 |
| ENSG00000187172 | BAGE2 | 0.040512 | 0.259449 | 0.00444 |
| ENSG00000112851 | ERBIN | 0.859449 | 0.632517 | 9.47E-07 |
| ENSG00000162576 | MXRA8 | 0.73406 | 0.451954 | 2.07E-08 |
| ENSG00000163806 | SPDYA | 0.198869 | 0.530563 | 7.74E-08 |
| ENSG00000187173 | LCE2A | 0.542597 | 0.298436 | 0.038494 |
| ENSG00000264813 | AC113554.1 | 0.604931 | 0.355111 | 1.63E-06 |
| ENSG00000101182 | PSMA7 | 0.977562 | 0.58959 | 7.80E-07 |
| ENSG00000103528 | SYT17 | 0.492277 | 0.22841 | 4.49E-08 |
| ENSG00000196553 | CCDC196 | 0.060998 | 0.28601 | 0.033217 |
| ENSG00000107281 | NPDC1 | 0.70077 | 0.390387 | 5.80E-09 |
| ENSG00000250317 | SMIM20 | 0.748466 | 0.428261 | 0.011129 |
| ENSG00000170509 | HSD17B13 | 0.107345 | 0.307979 | 0.000182 |
| ENSG00000166716 | ZNF592 | 0.849113 | 0.628897 | 2.26E-05 |
| ENSG00000212747 | RTL8B | 0.688822 | 0.350379 | 1.26E-13 |
| ENSG00000171658 | NMRAL2P | 0.472423 | 0.213914 | 1.66E-13 |
| ENSG00000168488 | ATXN2L | 0.868382 | 0.590178 | 0.000372 |
| ENSG00000107984 | DKK1 | 0.048297 | 0.266953 | 0.001418 |
| ENSG00000162669 | HFM1 | 0.046142 | 0.266518 | 0.041739 |
| ENSG00000105723 | GSK3A | 0.84003 | 0.608318 | 0.03818 |
| ENSG00000005379 | TSPOAP1 | 0.479008 | 0.687004 | 9.92E-09 |
| ENSG00000228526 | MIR34AHG | 0.694207 | 0.414671 | 0.000181 |
| ENSG00000211450 | SELENOH | 0.751639 | 0.449565 | 0.002437 |
| ENSG00000173928 | SWSAP1 | 0.634053 | 0.285616 | 0.000335 |
| ENSG00000140254 | DUOXA1 | 0.620795 | 0.266352 | 2.91E-13 |
| ENSG00000185246 | PRPF39 | 0.733118 | 0.499741 | 2.23E-12 |
| ENSG00000154518 | ATP5MC3 | 0.663002 | 0.453243 | 1.17E-07 |
| ENSG00000110887 | DAO | 0.065784 | 0.276261 | 0.000911 |
| ENSG00000197826 | C4orf22 | 0.045258 | 0.257151 | 0.039024 |
| ENSG00000227124 | ZNF717 | 0.707915 | 0.374634 | 0.000209 |
| ENSG00000105771 | SMG9 | 0.746788 | 0.474818 | 0.000601 |
| ENSG00000235865 | GSN-AS1 | 0.795672 | 0.421625 | 7.16E-28 |
| ENSG00000197183 | NOL4L | 0.74623 | 0.545583 | 0.000422 |
| ENSG00000135144 | DTX1 | 0.543789 | 0.342409 | 0.000134 |
| ENSG00000127399 | LRRC61 | 0.63733 | 0.361982 | 1.36E-12 |
| ENSG00000160563 | MED27 | 0.808727 | 0.442792 | 1.22E-16 |
| ENSG00000175054 | ATR | 0.772394 | 0.533826 | 0.009473 |
| ENSG00000263006 | ROCK1P1 | 0.132911 | 0.33343 | 1.48E-06 |
| ENSG00000162813 | BPNT1 | 0.599488 | 0.271237 | 2.06E-11 |
| ENSG00000037241 | RPL26L1 | 0.585525 | 0.35898 | 4.90E-11 |
| ENSG00000170629 | DPY19L2P2 | 0.096619 | 0.311853 | 0.041289 |
| ENSG00000135390 | ATP5MC2 | 0.948433 | 0.578742 | 0.026081 |
| ENSG00000104154 | SLC30A4 | 0.861998 | 0.427569 | 8.91E-09 |
| ENSG00000174946 | GPR171 | 0.765066 | 0.374571 | 5.91E-11 |
| ENSG00000129244 | ATP1B2 | 0.71024 | 0.335625 | 5.16E-12 |
| ENSG00000122824 | NUDT10 | 0.52244 | 0.294203 | 0.033429 |
| ENSG00000151704 | KCNJ1 | 0.096465 | 0.320918 | 1.07E-08 |
| ENSG00000101138 | CSTF1 | 0.737861 | 0.419686 | 1.67E-10 |
| ENSG00000177830 | CHID1 | 0.801567 | 0.488875 | 1.25E-06 |
| ENSG00000111341 | MGP | 0.052877 | 0.278542 | 0.020414 |
| ENSG00000143748 | NVL | 0.63406 | 0.431879 | 0.000435 |
| ENSG00000185668 | POU3F1 | 0.555555 | 0.344606 | 0.000239 |
| ENSG00000116353 | MECR | 0.545944 | 0.331122 | 0.005676 |
| ENSG00000235110 | AC107421.1 | 0.065755 | 0.27188 | 0.001402 |
| ENSG00000258469 | CHMP4BP1 | 0.770565 | 0.461323 | 1.49E-13 |
| ENSG00000141540 | TTYH2 | 0.846187 | 0.628116 | 4.71E-06 |
| ENSG00000099866 | MADCAM1 | 0.648653 | 0.330037 | 0.006983 |
| ENSG00000179941 | BBS10 | 0.599828 | 0.333449 | 1.73E-05 |
| ENSG00000135312 | HTR1B | 0.480166 | 0.240847 | 0.002599 |
| ENSG00000135999 | EPC2 | 0.741496 | 0.498418 | 0.00054 |
| ENSG00000183114 | FAM43B | 0.563868 | 0.3008 | 0.007079 |
| ENSG00000080573 | COL5A3 | 0.525588 | 0.273146 | 0.000203 |
| ENSG00000185798 | WDR53 | 0.639588 | 0.408953 | 1.85E-10 |
| ENSG00000155974 | GRIP1 | 0.158774 | 0.394355 | 7.97E-10 |
| ENSG00000244005 | NFS1 | 0.71792 | 0.35529 | 7.09E-10 |
| ENSG00000124172 | ATP5F1E | 0.979932 | 0.565354 | 8.51E-07 |
| ENSG00000168062 | BATF2 | 0.533912 | 0.26669 | 0.000238 |
| ENSG00000116194 | ANGPTL1 | 0.07222 | 0.306203 | 2.44E-08 |
| ENSG00000132481 | TRIM47 | 0.623383 | 0.296365 | 0.002175 |
| ENSG00000235028 | HMGN1P30 | 0.590272 | 0.341489 | 1.29E-07 |
| ENSG00000172016 | REG3A | 0.06336 | 0.28824 | 1.41E-15 |
| ENSG00000184608 | FAM167A-AS1 | 0.064521 | 0.285714 | 6.67E-07 |
| ENSG00000236062 | GSTM5P1 | 0.483744 | 0.28263 | 0.002395 |
| ENSG00000260314 | MRC1 | 0.853575 | 0.469177 | 0.001593 |
| ENSG00000105825 | TFPI2 | 0.540284 | 0.23187 | 9.79E-13 |
| ENSG00000136541 | ERMN | 0.168833 | 0.461754 | 7.20E-10 |
| ENSG00000228979 | PPIAP55 | 0.566516 | 0.249146 | 0.000197 |
| ENSG00000155254 | MARVELD1 | 0.799607 | 0.47345 | 0.027502 |
| ENSG00000123200 | ZC3H13 | 0.85244 | 0.553041 | 2.75E-08 |
| ENSG00000114767 | RRP9 | 0.632784 | 0.407855 | 0.000327 |
| ENSG00000187135 | VSTM2B | 0.523321 | 0.298302 | 0.002045 |
| ENSG00000118520 | ARG1 | 0.125838 | 0.334896 | 1.30E-08 |
| ENSG00000161217 | PCYT1A | 0.882128 | 0.572663 | 2.25E-06 |
| ENSG00000033122 | LRRC7 | 0.132486 | 0.357787 | 1.77E-08 |
| ENSG00000168702 | LRP1B | 0.083715 | 0.345744 | 4.07E-08 |
| ENSG00000150938 | CRIM1 | 0.661337 | 0.413181 | 0.00021 |
| ENSG00000163328 | GPR155 | 0.677044 | 0.425787 | 0.011393 |
| ENSG00000072518 | MARK2 | 0.751561 | 0.530084 | 0.000196 |
| ENSG00000157613 | CREB3L1 | 0.55985 | 0.250776 | 1.67E-21 |
| ENSG00000073803 | MAP3K13 | 0.741507 | 0.385529 | 4.16E-11 |
| ENSG00000154277 | UCHL1 | 0.620695 | 0.26054 | 1.84E-20 |
| ENSG00000185198 | PRSS57 | 0.584735 | 0.332828 | 5.31E-05 |
| ENSG00000197134 | ZNF257 | 0.822537 | 0.372208 | 0.000147 |
| ENSG00000116062 | MSH6 | 0.831312 | 0.533324 | 1.51E-07 |
| ENSG00000136367 | ZFHX2 | 0.490842 | 0.285431 | 0.027974 |
| ENSG00000215424 | MCM3AP-AS1 | 0.674992 | 0.431072 | 6.69E-10 |
| ENSG00000165915 | SLC39A13 | 0.784365 | 0.477188 | 2.87E-10 |
| ENSG00000128039 | SRD5A3 | 0.67133 | 0.368432 | 0.007106 |
| ENSG00000165972 | CCDC38 | 0.094773 | 0.299979 | 1.60E-07 |
| ENSG00000163933 | RFT1 | 0.706396 | 0.462125 | 0.003398 |
| ENSG00000196267 | ZNF836 | 0.608998 | 0.375849 | 4.60E-14 |
| ENSG00000141568 | FOXK2 | 0.811559 | 0.563445 | 0.004133 |
| ENSG00000162366 | PDZK1IP1 | 0.446826 | 0.246789 | 0.019853 |
| ENSG00000181826 | RELL1 | 0.679604 | 0.467801 | 0.000498 |
| ENSG00000130203 | APOE | 0.669513 | 0.447245 | 0.002613 |
| ENSG00000138434 | SSFA2 | 0.827653 | 0.59451 | 2.68E-06 |
| ENSG00000168938 | PPIC | 0.71363 | 0.245936 | 1.64E-24 |
| ENSG00000177483 | RBM44 | 0.118856 | 0.384191 | 3.93E-10 |
| ENSG00000164761 | TNFRSF11B | 0.454993 | 0.168558 | 8.71E-11 |
| ENSG00000082213 | C5orf22 | 0.682288 | 0.404503 | 0.037163 |
| ENSG00000234589 | AC125807.1 | 0.679495 | 0.323348 | 1.68E-06 |
| ENSG00000188001 | TPRG1 | 0.52483 | 0.291182 | 1.98E-12 |
| ENSG00000240225 | ZNF542P | 0.599082 | 0.346246 | 4.77E-05 |
| ENSG00000199568 | RNU5A-1 | 0.088739 | 0.353947 | 1.20E-22 |
| ENSG00000181904 | C5orf24 | 0.844983 | 0.499586 | 0.002601 |
| ENSG00000111850 | SMIM8 | 0.572584 | 0.320633 | 7.16E-06 |
| ENSG00000112062 | MAPK14 | 0.799054 | 0.495592 | 0.011445 |
| ENSG00000188994 | ZNF292 | 0.819915 | 0.559911 | 0.026971 |
| ENSG00000101966 | XIAP | 0.723823 | 0.48321 | 0.031241 |
| ENSG00000180385 | EMC3-AS1 | 0.637199 | 0.341561 | 5.34E-06 |
| ENSG00000135127 | BICDL1 | 0.579454 | 0.339434 | 0.000246 |
| ENSG00000187504 | RPL7P48 | 0.519705 | 0.265054 | 0.006167 |
| ENSG00000119401 | TRIM32 | 0.644948 | 0.343395 | 8.18E-15 |
| ENSG00000149635 | OCSTAMP | 0.452096 | 0.233904 | 0.000102 |
| ENSG00000202515 | VTRNA1-3 | 0.479966 | 0.243799 | 1.11E-08 |
| ENSG00000111224 | PARP11 | 0.629174 | 0.396791 | 0.004788 |
| ENSG00000170891 | CYTL1 | 0.551927 | 0.285619 | 2.87E-05 |
| ENSG00000096264 | NCR2 | 0.492113 | 0.273945 | 0.001416 |
| ENSG00000081870 | HSPB11 | 0.865435 | 0.429901 | 1.10E-08 |
| ENSG00000179134 | SAMD4B | 0.74957 | 0.509642 | 0.00121 |
| ENSG00000168348 | INSM2 | 0.417097 | 0.160358 | 0.013955 |
| ENSG00000137216 | TMEM63B | 0.686213 | 0.424615 | 0.014124 |
| ENSG00000077782 | FGFR1 | 0.581249 | 0.332452 | 2.16E-08 |
| ENSG00000143819 | EPHX1 | 0.695302 | 0.407499 | 3.28E-05 |
| ENSG00000072062 | PRKACA | 0.841917 | 0.563741 | 1.79E-06 |
| ENSG00000152422 | XRCC4 | 0.549218 | 0.293485 | 4.38E-10 |
| ENSG00000136542 | GALNT5 | 0.067053 | 0.277152 | 4.68E-09 |
| ENSG00000236138 | DUX4L26 | 0.525819 | 0.282621 | 8.01E-08 |
| ENSG00000099998 | GGT5 | 0.703236 | 0.247363 | 2.65E-19 |
| ENSG00000108691 | CCL2 | 0.773143 | 0.284309 | 0.00077 |
| ENSG00000125398 | SOX9 | 0.506253 | 0.260758 | 0.002993 |
| ENSG00000124429 | POF1B | 0.043888 | 0.254801 | 0.002753 |
| ENSG00000077063 | CTTNBP2 | 0.698019 | 0.296834 | 1.33E-12 |
| ENSG00000163635 | ATXN7 | 0.88149 | 0.564623 | 7.06E-06 |
| ENSG00000232150 | ST13P4 | 0.597065 | 0.323728 | 3.71E-08 |
| ENSG00000188282 | RUFY4 | 0.656001 | 0.393184 | 2.58E-05 |
| ENSG00000136546 | SCN7A | 0.117517 | 0.332463 | 0.001067 |
| ENSG00000135801 | TAF5L | 0.730563 | 0.454965 | 0.003668 |
| ENSG00000135253 | KCP | 0.708668 | 0.422131 | 0.000752 |
| ENSG00000105662 | CRTC1 | 0.641126 | 0.426523 | 1.15E-07 |
| ENSG00000122952 | ZWINT | 0.58633 | 0.31794 | 1.07E-11 |
| ENSG00000119899 | SLC17A5 | 0.763208 | 0.39123 | 2.23E-13 |
| ENSG00000161298 | ZNF382 | 0.579735 | 0.281919 | 2.47E-22 |
| ENSG00000173660 | UQCRH | 0.834397 | 0.492083 | 2.32E-10 |
| ENSG00000103152 | MPG | 0.843477 | 0.429529 | 3.79E-10 |
| ENSG00000186766 | FOXI2 | 0.464665 | 0.251869 | 0.000184 |
| ENSG00000135870 | RC3H1 | 0.847529 | 0.579204 | 0.000681 |
| ENSG00000165272 | AQP3 | 0.791364 | 0.345821 | 1.16E-05 |
| ENSG00000173769 | TOPAZ1 | 0.467382 | 0.255087 | 1.00E-07 |
| ENSG00000130643 | CALY | 0.552239 | 0.277822 | 0.027348 |
| ENSG00000167779 | IGFBP6 | 0.558089 | 0.274286 | 0.03645 |
| ENSG00000171224 | FAM241B | 0.517068 | 0.221306 | 0.001452 |
| ENSG00000167136 | ENDOG | 0.849211 | 0.400343 | 2.78E-20 |
| ENSG00000127954 | STEAP4 | 0.562699 | 0.258615 | 1.65E-06 |
| ENSG00000123411 | IKZF4 | 0.64758 | 0.374116 | 0.000407 |
| ENSG00000199038 | MIR210 | 0.636756 | 0.256097 | 0.006009 |
| ENSG00000102572 | STK24 | 0.795911 | 0.549365 | 0.002191 |
| ENSG00000198663 | C6orf89 | 0.842227 | 0.486159 | 0.015953 |
| ENSG00000151657 | KIN | 0.639611 | 0.397561 | 0.002527 |
| ENSG00000106178 | CCL24 | 0.585994 | 0.23966 | 2.85E-06 |
| ENSG00000176435 | CLEC14A | 0.501716 | 0.28566 | 0.000555 |
| ENSG00000125861 | GFRA4 | 0.491066 | 0.253587 | 0.000227 |
| ENSG00000140859 | KIFC3 | 0.57844 | 0.32256 | 8.56E-06 |
| ENSG00000216863 | LY86-AS1 | 0.563117 | 0.276317 | 0.027226 |
| ENSG00000090621 | PABPC4 | 0.867522 | 0.581338 | 1.02E-06 |
| ENSG00000237440 | ZNF737 | 0.745834 | 0.334088 | 1.58E-09 |
| ENSG00000104921 | FCER2 | 0.696935 | 0.25322 | 4.82E-07 |
| ENSG00000131023 | LATS1 | 0.768874 | 0.523762 | 0.031163 |
| ENSG00000111052 | LIN7A | 0.43702 | 0.211009 | 3.23E-08 |
| ENSG00000197860 | SGTB | 0.789616 | 0.427589 | 0.000186 |
| ENSG00000184270 | HIST2H2AB | 0.711662 | 0.487009 | 1.64E-05 |
| ENSG00000234292 | AC123595.1 | 0.569982 | 0.278189 | 0.000158 |
| ENSG00000102241 | HTATSF1 | 0.774067 | 0.4517 | 1.82E-05 |
| ENSG00000112210 | RAB23 | 0.488581 | 0.280276 | 2.58E-07 |
| ENSG00000267278 | MAP3K14-AS1 | 0.704731 | 0.410582 | 0.00357 |
| ENSG00000128512 | DOCK4 | 0.718113 | 0.353336 | 9.68E-06 |
| ENSG00000112146 | FBXO9 | 0.808679 | 0.467356 | 2.27E-09 |
| ENSG00000184731 | FAM110C | 0.464233 | 0.239294 | 0.001591 |
| ENSG00000185915 | KLHL34 | 0.422506 | 0.210423 | 0.005347 |
| ENSG00000162139 | NEU3 | 0.786098 | 0.408713 | 5.06E-07 |
| ENSG00000102158 | MAGT1 | 0.876004 | 0.48309 | 3.74E-06 |
| ENSG00000165914 | TTC7B | 0.693848 | 0.354177 | 6.66E-10 |
| ENSG00000167992 | VWCE | 0.537824 | 0.328638 | 0.000858 |
| ENSG00000120337 | TNFSF18 | 0.407096 | 0.198121 | 5.26E-13 |
| ENSG00000180438 | TPRXL | 0.472 | 0.237665 | 0.009213 |
| ENSG00000169840 | GSX1 | 0.485575 | 0.234107 | 2.79E-07 |
| ENSG00000165527 | ARF6 | 0.84613 | 0.550483 | 9.95E-07 |
| ENSG00000063515 | GSC2 | 0.451831 | 0.226646 | 5.12E-08 |
| ENSG00000149231 | CCDC82 | 0.702408 | 0.403129 | 2.81E-05 |
| ENSG00000203724 | C1orf53 | 0.497821 | 0.195797 | 2.55E-07 |
| ENSG00000164022 | AIMP1 | 0.836313 | 0.416549 | 1.74E-07 |
| ENSG00000141867 | BRD4 | 0.832886 | 0.552906 | 0.0333 |
| ENSG00000089335 | ZNF302 | 0.570548 | 0.329643 | 1.40E-09 |
| ENSG00000164512 | ANKRD55 | 0.507373 | 0.226806 | 1.09E-09 |
| ENSG00000167904 | TMEM68 | 0.520017 | 0.295657 | 2.08E-22 |
| ENSG00000064545 | TMEM161A | 0.660112 | 0.43586 | 0.024244 |
| ENSG00000198455 | ZXDB | 0.648575 | 0.430395 | 4.02E-11 |
| ENSG00000155313 | USP25 | 0.846661 | 0.489934 | 0.000164 |
| ENSG00000127377 | CRYGN | 0.45229 | 0.228099 | 0.000254 |
| ENSG00000130340 | SNX9 | 0.850096 | 0.50647 | 0.007085 |
| ENSG00000162520 | SYNC | 0.580656 | 0.365921 | 9.93E-10 |
| ENSG00000142082 | SIRT3 | 0.62878 | 0.412732 | 0.023942 |
| ENSG00000205795 | CYS1 | 0.564732 | 0.235432 | 0.034953 |
| ENSG00000128714 | HOXD13 | 0.411452 | 0.209801 | 2.36E-05 |
| ENSG00000212402 | SNORA74B | 0.618677 | 0.277357 | 0.000321 |
| ENSG00000174032 | SLC25A30 | 0.775269 | 0.429078 | 0.018058 |
| ENSG00000113638 | TTC33 | 0.50131 | 0.253427 | 1.05E-16 |
| ENSG00000137073 | UBAP2 | 0.817803 | 0.507478 | 0.000702 |
| ENSG00000146360 | GPR6 | 0.440206 | 0.22001 | 0.000105 |
| ENSG00000267384 | SMCO4P1 | 0.592064 | 0.206126 | 0.00272 |
| ENSG00000162062 | TEDC2 | 0.585537 | 0.310582 | 0.040709 |
| ENSG00000100842 | EFS | 0.446222 | 0.220539 | 0.040572 |
| ENSG00000106086 | PLEKHA8 | 0.613018 | 0.377429 | 0.000908 |
| ENSG00000130382 | MLLT1 | 0.697962 | 0.454361 | 6.90E-05 |
| ENSG00000162129 | CLPB | 0.654823 | 0.430165 | 0.00773 |
| ENSG00000145819 | ARHGAP26 | 0.771166 | 0.474849 | 0.039854 |
| ENSG00000171159 | C9orf16 | 0.85891 | 0.452425 | 0.011489 |
| ENSG00000136695 | IL36RN | 0.402272 | 0.185006 | 5.45E-12 |
| ENSG00000105379 | ETFB | 0.842328 | 0.428492 | 0.000606 |
| ENSG00000089048 | ESF1 | 0.63943 | 0.33637 | 1.69E-09 |
| ENSG00000130517 | PGPEP1 | 0.60417 | 0.387896 | 9.01E-05 |
| ENSG00000167548 | KMT2D | 0.779005 | 0.519843 | 3.26E-05 |
| ENSG00000143632 | ACTA1 | 0.571455 | 0.27654 | 0.003216 |
| ENSG00000087245 | MMP2 | 0.418061 | 0.210899 | 4.85E-05 |
| ENSG00000168944 | CEP120 | 0.769266 | 0.442392 | 0.010472 |
| ENSG00000072840 | EVC | 0.526556 | 0.277045 | 0.000124 |
| ENSG00000169397 | RNASE3 | 0.5632 | 0.228994 | 0.004585 |
| ENSG00000204612 | FOXB2 | 0.526342 | 0.227875 | 5.46E-07 |
| ENSG00000179271 | GADD45GIP1 | 0.823178 | 0.437167 | 9.04E-06 |
| ENSG00000174307 | PHLDA3 | 0.603154 | 0.323997 | 0.000211 |
| ENSG00000075131 | TIPIN | 0.550176 | 0.28384 | 2.01E-05 |
| ENSG00000134595 | SOX3 | 0.582152 | 0.220375 | 5.03E-05 |
| ENSG00000168298 | HIST1H1E | 0.816724 | 0.567217 | 0.000414 |
| ENSG00000179933 | C14orf119 | 0.911685 | 0.361771 | 3.51E-51 |
| ENSG00000168282 | MGAT2 | 0.843518 | 0.425168 | 4.14E-16 |
| ENSG00000142538 | PTH2 | 0.558005 | 0.226434 | 0.027917 |
| ENSG00000162408 | NOL9 | 0.64513 | 0.383799 | 0.001143 |
| ENSG00000164933 | SLC25A32 | 0.773586 | 0.420202 | 3.28E-05 |
| ENSG00000168505 | GBX2 | 0.441935 | 0.213595 | 1.07E-11 |
| ENSG00000180535 | BHLHA15 | 0.464614 | 0.229013 | 3.67E-09 |
| ENSG00000100147 | CCDC134 | 0.507205 | 0.294931 | 0.015917 |
| ENSG00000262180 | OCLM | 0.56036 | 0.303715 | 0.022688 |
| ENSG00000205439 | KRTAP12-3 | 0.413754 | 0.201866 | 0.042186 |
| ENSG00000172428 | COPS9 | 0.864075 | 0.362459 | 1.49E-20 |
| ENSG00000167157 | PRRX2 | 0.544815 | 0.188196 | 0.020119 |
| ENSG00000245680 | ZNF585B | 0.552144 | 0.28723 | 0.002942 |
| ENSG00000101096 | NFATC2 | 0.850262 | 0.484305 | 3.20E-09 |
| ENSG00000105088 | OLFM2 | 0.616504 | 0.272422 | 0.001461 |
| ENSG00000186847 | KRT14 | 0.448859 | 0.194785 | 0.003902 |
| ENSG00000100393 | EP300 | 0.942193 | 0.508675 | 0.027243 |
| ENSG00000124217 | MOCS3 | 0.465825 | 0.2623 | 7.73E-11 |
| ENSG00000226415 | TPI1P1 | 0.844182 | 0.420859 | 0.038816 |
| ENSG00000133935 | ERG28 | 0.894701 | 0.405118 | 8.42E-06 |
| ENSG00000121152 | NCAPH | 0.724383 | 0.342969 | 1.42E-07 |
| ENSG00000198680 | TUSC1 | 0.482767 | 0.258762 | 1.22E-08 |
| ENSG00000205426 | KRT81 | 0.41347 | 0.17257 | 0.04833 |
| ENSG00000269001 | AC092070.2 | 0.563495 | 0.234094 | 1.23E-22 |
| ENSG00000087076 | HSD17B14 | 0.429213 | 0.214769 | 8.94E-05 |
| ENSG00000162367 | TAL1 | 0.530936 | 0.223413 | 1.70E-09 |
| ENSG00000101132 | PFDN4 | 0.657431 | 0.298228 | 1.66E-12 |
| ENSG00000197588 | KLKP1 | 0.358351 | 0.150171 | 0.002616 |
| ENSG00000127561 | SYNGR3 | 0.575243 | 0.224526 | 9.63E-10 |
| ENSG00000106723 | SPIN1 | 0.708646 | 0.446052 | 7.34E-06 |
| ENSG00000131094 | C1QL1 | 0.531935 | 0.214216 | 0.003385 |
| ENSG00000164976 | MYORG | 0.555367 | 0.254323 | 2.24E-09 |
| ENSG00000165476 | REEP3 | 0.874554 | 0.380026 | 3.82E-06 |
| ENSG00000186567 | CEACAM19 | 0.5424 | 0.257698 | 5.02E-14 |
| ENSG00000125046 | SSUH2 | 0.417732 | 0.191759 | 0.001424 |
| ENSG00000106993 | CDC37L1 | 0.682595 | 0.289999 | 5.58E-21 |
| ENSG00000154265 | ABCA5 | 0.662653 | 0.336267 | 1.41E-05 |
| ENSG00000131653 | TRAF7 | 0.767089 | 0.45542 | 0.000194 |
| ENSG00000253831 | ETV3L | 0.357738 | 0.144432 | 4.17E-06 |
| ENSG00000107159 | CA9 | 0.400729 | 0.195184 | 0.005422 |
| ENSG00000161692 | DBF4B | 0.554251 | 0.293001 | 1.93E-08 |
| ENSG00000181656 | GPR88 | 0.514104 | 0.221325 | 6.09E-05 |
| ENSG00000104140 | RHOV | 0.549335 | 0.219457 | 0.040896 |
| ENSG00000110844 | PRPF40B | 0.742451 | 0.36134 | 3.92E-05 |
| ENSG00000151715 | TMEM45B | 0.612386 | 0.247013 | 5.14E-07 |
| ENSG00000145246 | ATP10D | 0.685655 | 0.394604 | 0.000211 |
| ENSG00000205364 | MT1M | 0.390044 | 0.145011 | 9.97E-09 |
| ENSG00000164379 | FOXQ1 | 0.522377 | 0.169699 | 2.43E-05 |
| ENSG00000188779 | SKOR1 | 0.498472 | 0.266163 | 2.36E-19 |
| ENSG00000033011 | ALG1 | 0.643191 | 0.310893 | 0.014637 |
| ENSG00000105523 | FAM83E | 0.478 | 0.209489 | 4.15E-06 |
| ENSG00000100234 | TIMP3 | 0.706413 | 0.229348 | 1.53E-13 |
| ENSG00000178860 | MSC | 0.724263 | 0.200215 | 3.06E-17 |
| ENSG00000099377 | HSD3B7 | 0.62022 | 0.287296 | 8.56E-05 |
| ENSG00000205899 | BHLHA9 | 0.444201 | 0.193561 | 2.14E-08 |
| ENSG00000134668 | SPOCD1 | 0.44454 | 0.180404 | 8.20E-05 |
| ENSG00000081386 | ZNF510 | 0.526009 | 0.286352 | 0.001073 |
| ENSG00000165506 | DNAAF2 | 0.6057 | 0.288469 | 2.38E-06 |
| ENSG00000196510 | ANAPC7 | 0.839762 | 0.370979 | 1.17E-21 |
| ENSG00000161677 | JOSD2 | 0.72664 | 0.323768 | 4.75E-08 |
| ENSG00000145391 | SETD7 | 0.707859 | 0.439679 | 0.005085 |
| ENSG00000187189 | TSPYL4 | 0.54855 | 0.345748 | 9.80E-11 |
| ENSG00000007923 | DNAJC11 | 0.76418 | 0.375139 | 2.55E-09 |
| ENSG00000174407 | MIR1-1HG | 0.482632 | 0.176755 | 0.046285 |
| ENSG00000188735 | TMEM120B | 0.824746 | 0.430391 | 4.16E-17 |
| ENSG00000184423 | RPL23AP38 | 0.451991 | 0.191025 | 0.00017 |
| ENSG00000159958 | TNFRSF13C | 0.536791 | 0.32944 | 0.04068 |
| ENSG00000162971 | TYW5 | 0.621244 | 0.326958 | 8.17E-06 |
| ENSG00000105329 | TGFB1 | 0.77601 | 0.50703 | 7.99E-19 |
| ENSG00000204335 | SP5 | 0.45125 | 0.185281 | 2.99E-06 |
| ENSG00000105221 | AKT2 | 0.845943 | 0.468505 | 5.19E-13 |
| ENSG00000147804 | SLC39A4 | 0.657743 | 0.313499 | 0.00323 |
| ENSG00000127870 | RNF6 | 0.722328 | 0.421078 | 0.019844 |
| ENSG00000130856 | ZNF236 | 0.642691 | 0.413195 | 0.000281 |
| ENSG00000188763 | FZD9 | 0.496108 | 0.197403 | 0.012533 |
| ENSG00000169169 | CPT1C | 0.384887 | 0.179028 | 0.003345 |
| ENSG00000160396 | HIPK4 | 0.428353 | 0.225009 | 0.008333 |
| ENSG00000088827 | SIGLEC1 | 0.789945 | 0.371333 | 0.001443 |
| ENSG00000182141 | ZNF708 | 0.626009 | 0.310825 | 0.001537 |
| ENSG00000196867 | ZFP28 | 0.616452 | 0.232553 | 4.93E-17 |
| ENSG00000171794 | UTF1 | 0.550311 | 0.293353 | 0.000136 |
| ENSG00000175229 | GAL3ST3 | 0.384012 | 0.169409 | 2.46E-09 |
| ENSG00000109586 | GALNT7 | 0.749839 | 0.382199 | 0.020959 |
| ENSG00000073067 | CYP2W1 | 0.419892 | 0.210984 | 2.32E-16 |
| ENSG00000179331 | RAB39A | 0.581654 | 0.26348 | 0.001079 |
| ENSG00000089902 | RCOR1 | 0.826496 | 0.47533 | 3.54E-14 |
| ENSG00000028277 | POU2F2 | 0.641777 | 0.422035 | 2.11E-13 |
| ENSG00000171992 | SYNPO | 0.609309 | 0.238945 | 2.45E-09 |
| ENSG00000148488 | ST8SIA6 | 0.446398 | 0.188393 | 2.94E-05 |
| ENSG00000111665 | CDCA3 | 0.556386 | 0.258587 | 7.83E-07 |
| ENSG00000123444 | KBTBD4 | 0.712493 | 0.338708 | 4.33E-06 |
| ENSG00000105732 | ZNF574 | 0.632713 | 0.327921 | 0.001183 |
| ENSG00000173218 | VANGL1 | 0.59918 | 0.297658 | 2.55E-08 |
| ENSG00000119638 | NEK9 | 0.637305 | 0.412739 | 0.001371 |
| ENSG00000139618 | BRCA2 | 0.682487 | 0.342152 | 2.79E-05 |
| ENSG00000140396 | NCOA2 | 0.774538 | 0.452756 | 3.39E-08 |
| ENSG00000094661 | OR1I1 | 0.360113 | 0.133854 | 0.014402 |
| ENSG00000006377 | DLX6 | 0.535731 | 0.174219 | 6.62E-06 |
| ENSG00000182575 | NXPH3 | 0.529542 | 0.221738 | 0.03583 |
| ENSG00000197506 | SLC28A3 | 0.383219 | 0.163148 | 0.01573 |
| ENSG00000188784 | PLA2G2E | 0.363544 | 0.144737 | 0.02262 |
| ENSG00000070388 | FGF22 | 0.496001 | 0.192033 | 0.006873 |
| ENSG00000141664 | ZCCHC2 | 0.75239 | 0.439995 | 1.84E-12 |
| ENSG00000164935 | DCSTAMP | 0.457346 | 0.141939 | 4.26E-09 |
| ENSG00000186919 | ZACN | 0.652885 | 0.376769 | 0.013595 |
| ENSG00000112309 | B3GAT2 | 0.639384 | 0.323255 | 4.00E-12 |
| ENSG00000154144 | TBRG1 | 0.832942 | 0.42364 | 0.044832 |
| ENSG00000176769 | TCERG1L | 0.484254 | 0.187763 | 0.004927 |
| ENSG00000196811 | CHRNG | 0.525359 | 0.244294 | 9.83E-06 |
| ENSG00000105366 | SIGLEC8 | 0.422534 | 0.154747 | 9.63E-08 |
| ENSG00000132821 | VSTM2L | 0.575124 | 0.191915 | 0.025782 |
| ENSG00000149609 | C20orf144 | 0.558938 | 0.219016 | 1.08E-07 |
| ENSG00000143942 | CHAC2 | 0.449384 | 0.216705 | 9.30E-05 |
| ENSG00000164736 | SOX17 | 0.392967 | 0.160045 | 0.003941 |
| ENSG00000126012 | KDM5C | 0.798256 | 0.452493 | 7.52E-11 |
| ENSG00000180176 | TH | 0.351381 | 0.146097 | 0.006386 |
| ENSG00000187730 | GABRD | 0.379761 | 0.160991 | 2.89E-14 |
| ENSG00000186998 | EMID1 | 0.474718 | 0.180634 | 0.001075 |
| ENSG00000180929 | GPR62 | 0.432084 | 0.200226 | 1.75E-05 |
| ENSG00000160716 | CHRNB2 | 0.496433 | 0.245365 | 0.005313 |
| ENSG00000141376 | BCAS3 | 0.654951 | 0.353457 | 0.000141 |
| ENSG00000172155 | LCE1D | 0.615139 | 0.168581 | 8.18E-05 |
| ENSG00000172171 | TEFM | 0.586834 | 0.229375 | 2.73E-31 |
| ENSG00000122254 | HS3ST2 | 0.488577 | 0.160847 | 0.002512 |
| ENSG00000100253 | MIOX | 0.443381 | 0.185092 | 3.51E-09 |
| ENSG00000087589 | CASS4 | 0.632021 | 0.396726 | 2.97E-06 |
| ENSG00000189058 | APOD | 0.339217 | 0.136482 | 0.000969 |
| ENSG00000213171 | LINGO4 | 0.411885 | 0.157616 | 0.00648 |
| ENSG00000205213 | LGR4 | 0.400927 | 0.199412 | 3.06E-05 |
| ENSG00000145725 | PPIP5K2 | 0.824018 | 0.455941 | 0.001419 |
| ENSG00000170373 | CST1 | 0.350507 | 0.127558 | 1.00E-05 |
| ENSG00000164690 | SHH | 0.397836 | 0.174993 | 0.00027 |
| ENSG00000141744 | PNMT | 0.373245 | 0.166448 | 0.001548 |
| ENSG00000197013 | ZNF429 | 0.424421 | 0.22215 | 4.60E-10 |
| ENSG00000179673 | RPRML | 0.515198 | 0.183604 | 4.14E-09 |
| ENSG00000166448 | TMEM130 | 0.472732 | 0.157632 | 1.52E-05 |
| ENSG00000277224 | HIST1H2BF | 0.540901 | 0.287349 | 9.52E-06 |
| ENSG00000170949 | ZNF160 | 0.774982 | 0.349259 | 4.72E-14 |
| ENSG00000129911 | KLF16 | 0.634706 | 0.414212 | 4.03E-23 |
| ENSG00000173166 | RAPH1 | 0.790829 | 0.29394 | 1.28E-08 |
| ENSG00000124249 | KCNK15 | 0.507529 | 0.183759 | 1.19E-05 |
| ENSG00000133101 | CCNA1 | 0.59921 | 0.165362 | 1.34E-12 |
| ENSG00000132975 | GPR12 | 0.327185 | 0.118814 | 1.32E-08 |
| ENSG00000154240 | CEP112 | 0.521632 | 0.228367 | 7.35E-12 |
| ENSG00000121053 | EPX | 0.463465 | 0.20726 | 0.000111 |
| ENSG00000176753 | C15orf56 | 0.434529 | 0.162968 | 1.71E-05 |
| ENSG00000099822 | HCN2 | 0.458723 | 0.211467 | 2.74E-10 |
| ENSG00000203722 | RAET1G | 0.580185 | 0.151552 | 7.67E-06 |
| ENSG00000187238 | LCE3B | 0.567431 | 0.150833 | 0.001453 |
| ENSG00000188710 | QRFP | 0.505163 | 0.201292 | 0.000178 |
| ENSG00000187766 | KRTAP10-8 | 0.37715 | 0.136489 | 3.26E-10 |
| ENSG00000128965 | CHAC1 | 0.446149 | 0.173926 | 0.046549 |
| ENSG00000143869 | GDF7 | 0.488242 | 0.270962 | 1.40E-15 |
| ENSG00000198844 | ARHGEF15 | 0.478866 | 0.14396 | 0.000517 |
| ENSG00000112234 | FBXL4 | 0.758394 | 0.274728 | 1.74E-19 |
| ENSG00000105605 | CACNG7 | 0.34263 | 0.140609 | 6.64E-12 |
| ENSG00000185962 | LCE3A | 0.463264 | 0.176534 | 0.019962 |
| ENSG00000173714 | WFIKKN2 | 0.413563 | 0.144021 | 3.79E-07 |
| ENSG00000083099 | LYRM2 | 0.844889 | 0.333914 | 1.86E-16 |
| ENSG00000186897 | C1QL4 | 0.389656 | 0.144618 | 4.47E-06 |
| ENSG00000235718 | MFRP | 0.389059 | 0.133569 | 9.96E-06 |
| ENSG00000180613 | GSX2 | 0.504862 | 0.158067 | 0.003135 |
| ENSG00000156575 | PRG3 | 0.421859 | 0.128481 | 0.000108 |
| ENSG00000187566 | NHLRC1 | 0.380699 | 0.166645 | 8.28E-13 |
| ENSG00000101180 | HRH3 | 0.36136 | 0.154627 | 0.001281 |
| ENSG00000183161 | FANCF | 0.461845 | 0.252194 | 0.000153 |
| ENSG00000183072 | NKX2-5 | 0.523483 | 0.152346 | 6.22E-09 |
| ENSG00000148516 | ZEB1 | 0.727018 | 0.424569 | 4.57E-05 |
| ENSG00000130711 | PRDM12 | 0.444416 | 0.151894 | 7.41E-05 |
| ENSG00000162571 | TTLL10 | 0.46698 | 0.175295 | 2.10E-05 |
| ENSG00000197296 | FITM2 | 0.486047 | 0.210663 | 4.33E-06 |
| ENSG00000002330 | BAD | 0.69288 | 0.315057 | 0.030304 |
| ENSG00000197646 | PDCD1LG2 | 0.846014 | 0.171685 | 4.11E-26 |
| ENSG00000108688 | CCL7 | 0.437221 | 0.118014 | 0.001221 |
| ENSG00000111863 | ADTRP | 0.576767 | 0.141582 | 5.69E-15 |
| ENSG00000000457 | SCYL3 | 0.518539 | 0.2824 | 0.026013 |
| ENSG00000177842 | ZNF620 | 0.541108 | 0.213662 | 9.25E-09 |
| ENSG00000244486 | SCARF2 | 0.491192 | 0.274955 | 1.01E-17 |
| ENSG00000069482 | GAL | 0.550549 | 0.130374 | 5.75E-08 |
| ENSG00000103449 | SALL1 | 0.408682 | 0.116594 | 0.006718 |
| ENSG00000132563 | REEP2 | 0.467658 | 0.149999 | 4.40E-05 |
| ENSG00000170369 | CST2 | 0.521828 | 0.140429 | 5.72E-05 |
| ENSG00000167656 | LY6D | 0.47314 | 0.131797 | 0.001263 |
| ENSG00000176108 | CHMP6 | 0.617707 | 0.281954 | 7.31E-07 |
| ENSG00000186188 | FFAR4 | 0.595206 | 0.165711 | 1.27E-08 |
| ENSG00000120057 | SFRP5 | 0.399811 | 0.175927 | 6.95E-07 |
| ENSG00000198496 | NBR2 | 0.647759 | 0.234993 | 4.82E-07 |
| ENSG00000134452 | FBXO18 | 0.686584 | 0.353921 | 0.000147 |
| ENSG00000113763 | UNC5A | 0.422259 | 0.191436 | 0.020918 |
| ENSG00000100156 | SLC16A8 | 0.498077 | 0.193861 | 0.002785 |
| ENSG00000164603 | BMT2 | 0.747022 | 0.289756 | 2.65E-09 |
| ENSG00000034693 | PEX3 | 0.530686 | 0.227821 | 1.01E-16 |
| ENSG00000197961 | ZNF121 | 0.705811 | 0.387816 | 1.50E-08 |
| ENSG00000243910 | TUBA4B | 0.446381 | 0.16751 | 0.002069 |
| ENSG00000131435 | PDLIM4 | 0.662548 | 0.179456 | 6.38E-13 |
| ENSG00000186493 | C5orf38 | 0.398125 | 0.123488 | 5.88E-14 |
| ENSG00000254550 | OMP | 0.438586 | 0.137251 | 0.017649 |
| ENSG00000136883 | KIF12 | 0.372823 | 0.124734 | 3.20E-14 |
| ENSG00000131097 | HIGD1B | 0.441283 | 0.179556 | 0.010529 |
| ENSG00000187714 | SLC18A3 | 0.323682 | 0.117288 | 0.000338 |
| ENSG00000186198 | SLC51B | 0.347152 | 0.104069 | 3.33E-05 |
| ENSG00000144583 | 4-Mar | 0.339472 | 0.115335 | 0.002611 |
| ENSG00000010310 | GIPR | 0.512318 | 0.287655 | 0.04189 |
| ENSG00000162009 | SSTR5 | 0.320231 | 0.1087 | 7.55E-06 |
| ENSG00000172201 | ID4 | 0.362502 | 0.161474 | 0.03041 |
| ENSG00000178602 | OTOS | 0.476757 | 0.133368 | 0.001021 |
